# Transcriptional reprogramming of distinct peripheral sensory neuron subtypes after axonal injury

**DOI:** 10.1101/838854

**Authors:** William Renthal, Ivan Tochitsky, Lite Yang, Yung-Chih Cheng, Emmy Li, Riki Kawaguchi, Daniel H. Geschwind, Clifford J. Woolf

**Affiliations:** Department of Neurology, Brigham and Women’s Hospital and Harvard Medical School, 60 Fenwood Rd. Boston, MA 02115; Department of Neurobiology, Harvard Medical School, 220 Longwood Ave. Boston, MA 02115; F.M. Kirby Neurobiology Center, Boston Children’s Hospital, 3 Blackfan Cir. Boston, MA 02115; Program in Neurogenetics, Department of Neurology, David Geffen School of Medicine, University of California, Los Angeles, Los Angeles, CA

**Keywords:** Nerve injury, regeneration, sensory neuron, single cell RNA-seq, gene expression, dorsal root ganglion, reprogramming, cell identity, axon growth, Atf3

## Abstract

Primary somatosensory neurons are specialized to transmit specific types of sensory information through differences in cell size, myelination, and the expression of distinct receptors and ion channels, which together define their transcriptional and functional identity. By transcriptionally profiling sensory ganglia at single-cell resolution, we find that different somatosensory neuronal subtypes undergo a remarkably consistent and dramatic transcriptional response to peripheral nerve injury that both promotes axonal regeneration and suppresses cell identity. Successful axonal regeneration leads to a restoration of neuronal cell identity and the deactivation of the growth program. This injury-induced transcriptional reprogramming requires *Atf3*, a transcription factor which is induced rapidly after injury and is necessary for axonal regeneration and functional recovery. While *Atf3* and other injury-induced transcription factors are known for their role in reprogramming cell fate, their function in mature neurons is likely to facilitate major adaptive changes in cell function in response to damaging environmental stimuli.

## Introduction

Injury to peripheral axons of primary sensory neurons whose cell bodies reside in dorsal root ganglia (DRG) leads to the induction of cell-intrinsic transcriptional programs critical both for initiating axon growth and driving the pathological neuronal hyperexcitability that underlies neuropathic pain (Chandran et al., 2016; Costigan et al., 2002; He and Jin, 2016; Mahar and Cavalli, 2018; Scheib and Höke, 2013; Serra et al., 2012; Tuszynski and Steward, 2012). Axon regeneration involves both the regrowth of the injured axon and the correct reinnervation of its target, but this process is often incomplete and can lead both to a loss of sensation and disabling chronic painful neuropathies, such as phantom limb pain, diabetic neuropathy or chemotherapy-induced neuropathy (Chapman and Vierck, 2017; Collins et al., 2018; Xie et al., 2017). The molecular changes provoked by peripheral axonal injury have been the focus of intense study (Chandran et al., 2016; Costigan et al., 2002; He and Jin, 2016; Mahar and Cavalli, 2018; Scheib and Höke, 2013; Serra et al., 2012; Tuszynski and Steward, 2012) since the identification of the molecular drivers of regeneration has the potential to promote the regeneration of injured central nervous system neurons, which, unlike neurons with axons in the PNS, lack an intrinsic regeneration capacity (He and Jin, 2016; Mahar and Cavalli, 2018; Tuszynski and Steward, 2012). Additionally, a better understanding of the mechanisms by which neuronal hyperexcitability develops after axonal injury may reveal novel targets for analgesic development.

Previous molecular studies using bulk DRG tissue have identified transcriptional networks regulated in the DRG in response to injury (Abe and Cavalli, 2008; Chandran et al., 2016; Costigan et al., 2002; LaCroix-Fralish et al., 2011; Michaelevski et al., 2010; Perkins et al., 2014; Xiao et al., 2002). However, the extensive cellular heterogeneity of DRG cell types (Usoskin et al., 2015; Zeisel et al., 2018; Zheng et al., 2019) has made it difficult to establish in which cell types these changes occur and whether these changes are uniform or distinct across different neuronal subtypes. This challenge is underscored by the fact that non-neuronal cells, including satellite glia, Schwann cells, dural cells and endothelial cells, are collectively more abundant than sensory neurons in the DRG. Moreover, peripheral sensory neurons themselves vary dramatically in size, conduction velocity, gene expression patterns and the sensory transduction receptors present on nerve terminals (Gatto et al., 2019; Le Pichon and Chesler, 2014; Usoskin et al., 2015; Zeisel et al., 2018). In addition to the cellular heterogeneity within the DRG, in most nerve injury models, only a fraction of DRG neurons are injured and bulk analyses cannot differentiate between changes in injured or non-injured neurons (Berta et al., 2017; Gosselin et al., 2010; Jessen and Mirsky, 2016; Laedermann et al., 2014; Rigaud et al., 2008).

High-throughput single-nucleus genomics enables the characterization of axonal injury response programs within distinct cell types of the DRG, without use of cell dissociation procedures that themselves induce injury-like/immediate early gene responses (Chiu et al., 2014; Frey et al., 2015; Lindwall et al., 2004; Nguyen et al., 2019). Using droplet-based single-nucleus RNA sequencing (snRNA-seq) we mapped the transcriptomes of 107,541 individual mouse DRG cells across a range of nerve injury models. Remarkably, we find that axonal injury induces a common transcriptional program across all neuronal subtypes that largely replaces the expression of their subtype-specific genes. Non-neuronal cells exhibit a much smaller, distinct, transcriptional response to injury. The response of sensory neurons to injury involves the rapid induction of many of the transcription factors associated with reprogramming fibroblasts into either pluripotent stem cells or differentiated cell types (Brouwer et al., 2016), raising the possibility that neurons may invoke an analogous intrinsic transcriptional reprogramming for generating their response to axonal injury. We further demonstrate that *Atf3*, an axonal injury-induced transcription factor (Hunt et al., 2012; Parsadanian et al., 2006; Tsujino et al., 2000) also implicated in cellular reprogramming (Duan et al., 2019; Ronquist et al., 2017), is necessary for axotomy-induced neuronal transcriptional reprogramming and for axonal regeneration and sensory recovery after injury. Finally, we present a web-based resource for exploring changes in gene expression across DRG cell types (www.painseq.com) to aid fundamental studies of sensory neuron biology and development of novel therapeutics for pain and regeneration.

## Results

### Single-nucleus RNA-seq of naive and injured DRG cell types

To characterize transcriptional responses induced by peripheral axonal injury, we performed snRNA-seq on lumbar DRGs from adult naive mice and compared their transcriptional profiles to DRGs from mice after spinal nerve transection (SpNT) (the segmental nerve that emerges directly from each DRG), sciatic nerve transection (ScNT) or sciatic nerve crush (crush), over multiple time points, ranging from hours to months after injury (Figure 1A). Full axonal regeneration with target reinnervation and functional recovery is only observed in the sciatic crush model (Navarro et al., 1994). To determine whether nerve injury response is distinct from other pain-producing insults, we also characterized gene expression changes in lumbar DRGs from two models that do not involve physical axotomy: a model of acute (1 week) chemotherapy-induced allodynia (4mg/kg paclitaxel) (Velasco and Bruna, 2015) and a model of peripheral inflammation, hindpaw injection of Complete Freund’s Adjuvant (CFA, 20 µL, 2 days) (Jaggi et al., 2011).

**Figure 1.**
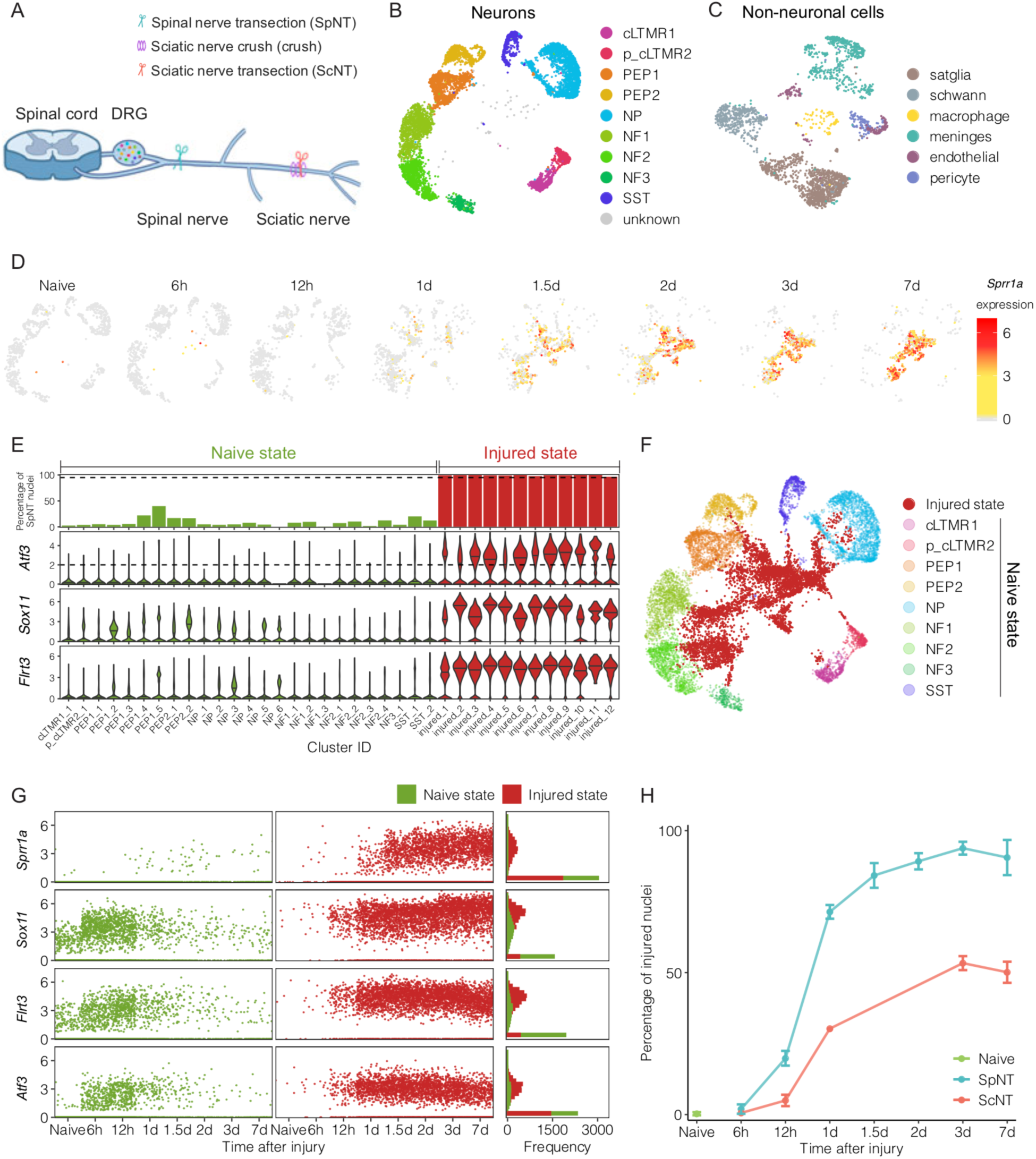
Single-nucleus RNA sequencing of DRG neurons in mouse models of peripheral axonal injury. (A) Diagram of mouse axotomy models. Spinal nerve transection (SpNT) is a proximal injury resulting in axotomy of 90+% of all neurons in a given DRG, whereas sciatic nerve transection (ScNT) and sciatic crush are distal injury models resulting in ∼50% of axotomized neurons on average across L3-L5 DRGs. (B) UMAP plot of 10,212 neuronal nuclei from naive mice. Clusters correspond to 9 neuronal subtypes and a small group of cells of unknown classification. (C) UMAP plot of 2,470 non-neuronal nuclei from naive mice representing 6 cell types. Satglia = satellite glia (D) UMAP plots displaying DRG neuronal subtypes expressing the injury-induced gene *Sprr1a* at different times after spinal nerve transection. Each time point was downsampled to display 900 nuclei. Color denotes log_2_-normalized expression of *Sprr1a;* nuclei not expressing *Sprr1a* are colored grey. (E) Bar plot showing the percent of SpNT nuclei [100 * SpNT nuclei / (naive + SpNT nuclei)] within each neuronal cluster (top row) and violin plots showing log_2_-normalized expression of selected injury-induced genes in each cluster (second to fourth rows). Fractions were calculated from a pool of 7,742 naive neuronal nuclei and 6,482 spinal nerve transection neuronal nuclei (> 1d). Cluster ID (x-axis) corresponds to cluster number assignment from Seurat (see Figure S1E, methods). Clusters are classified as “injured state” (red) if they are comprised of > 95% nuclei from SpNT mice and have a median normalized *Atf3* expression > 0.8 SD from mean (corresponding to > log_2_-normalized expression of 2). All other clusters are classified as “naive state” (green). (F) UMAP plot showing 7,000 naive neuronal nuclei and 7,000 randomly sampled SpNT neuronal nuclei. Nuclei classified as being in their “naive state” are colored by their assigned neuronal subtypes. Nuclei classified as in the “injured state” are colored red. (G) Scatter plot of the log_2_-normalized expression of four injury-induced genes (*Sprr1a, Atf3, Flrt3* and *Sox11*) in “naive state” (green) and “injured state” (red) nuclei. While there is little expression of *Atf3* and *Sprr1a* in the naive condition, there is some expression of *Flrt3* and *Sox11* in naive neurons. Within hours after injury, the expression of *Atf3, Flrt3, and Sox11* dramatically increases in neurons that are still classified as in the “uninjured state.” *Sprr1a* expression is largely absent in neurons until 1d after injury, the time point at which the “injured” transcriptional state emerges. These injury-induced genes remain increased for at least 7d. Each time point is downsampled to 900 nuclei for purposes of visualization. (H) Percentage of naive, SpNT, and ScNT neuronal nuclei that are classified as in the “injured state” at each time point after the respective injury. cLTMR = C-fiber low threshold mechanoreceptor; PEP = peptidergic nociceptor; NP = non-peptidergic nociceptor; NF = *Nefh+* A-fiber low threshold mechanoreceptors; SST = *Sst+* pruriceptors.

In total, we obtained 107,541 DRG nuclei that passed quality control (see methods). Sequenced nuclei had an average of 2,918 transcripts per nucleus representing 1,478 unique genes per nucleus (Figure S1A). For the purposes of cell type identification, DRG nuclei from naive and all experimental injury conditions were initially clustered together based on their gene expression patterns. Dimensionality reduction (uniform manifold approximation and projection [UMAP]) revealed 16 distinct groups of cells. Nuclei in clusters expressing high levels of *Rbfox3*, which encodes the pan-neuronal marker NeuN (Kim et al., 2009), were classified as neurons, and clusters expressing high levels of known non-neuronal marker genes, such as *Sparc*, were classified as non-neuronal nuclei (Figure S1B-C). We re-clustered neuronal and non-neuronal nuclei separately to better visualize their distinct subtypes and used this separate visualization in all subsequent analyses.

Focusing initially on naive DRG nuclei, the neuronal subtypes we observed include *Tac1*+ peptidergic nociceptors (PEP), *Mrgprd*+ non-peptidergic nociceptors (NP), *Sst*+ pruriceptors, *Fam19a4+/Th*+ low threshold mechano-receptive neurons with C-fibers (cLTMR), *Nefh*+ A fibers including A-LTMRs and proprioceptors (NF), (Figures 1B, S1C). Non-neuronal cells include *Apoe*+ satellite glia, *Mpz*+ Schwann cells and *Cldn5*+ endothelial cells (Figures 1C, S1C-D). The distinct neuronal and non-neuronal subtypes we identified in DRGs from naive animals were also observed in all injury models and are similar to those previously reported (Figure S1C-D) (Usoskin et al., 2015; Zeisel et al., 2018; Zheng et al., 2019). We also observed a neuronal cluster that expresses *Fam19a4*, but very low levels of *Th,* which we termed putative-cLTMR2 (p_cLTMR2). A subset of the cell type selective marker genes (Figures S1E-G), including those of p_cLTMR2 (Figure S1H), were studied by *in situ* hybridization and found largely to label distinct, non-overlapping cell populations (Usoskin et al., 2015; Zeisel et al., 2018; Zheng et al., 2019). In addition to the cell-type-specific gene expression patterns of known marker genes, we also observed distinct expression patterns of ion channels, G-protein coupled receptors (GPCRs), neuropeptides, and transcription factors (Figure S2A, Table S1, see methods). For example, we observed that PEP1 and PEP2 neurons express the ion channels *Trpv1* and *Atp2b4* and the GPCRs *Sstr2* (PEP1 only) and *Gpr26* (PEP2 only), as well as multiple neuropeptides including *Tac1, Adcyap1,* and *Calca* (PEP1 only), whereas NF1-3 neurons express the ion channels *Scn1b* and *Scn4b* and the GPCR *Adgrg2* (NF2,3 only), highlighting the molecular and functional differences between distinct subtypes of DRG neurons.

### Axonal injury induces a new transcriptional state in DRG neurons

To characterize the transcriptional programs activated in response to axonal injury, we first compared DRG nuclei from naive mice to DRG nuclei from mice 6 hours (h), 12h, 1day (d), 1.5d, 2d, 3d and 7d after transection of the spinal nerves from the respective ganglia, which results in the axotomy of >90% of DRG neurons in the affected ganglia (Shortland et al., 2006; Tsujino et al., 2000). Strikingly, we observed that new neuronal clusters emerge by 1d after SpNT, which are essentially absent in naive mice and which contain neurons that express very high levels of known injury-induced genes such as *Sprr1a* (Figure 1D). By 3 days after injury, few nuclei cluster with naive neurons, consistent with an axotomy of most DRG neurons. New injury-induced clusters of nuclei were not observed in non-neuronal cells (Figure S2B). To quantify the extent of injury among all neurons after SpNT, we defined the new neuronal clusters that emerged after the injury as an “injured state” if the cluster was comprised of greater than 95% SpNT nuclei and had a median normalized expression of *Atf3* greater than 2 (Figures 1E-F, S2C). *Atf3* is a major injury-induced gene in axon-damaged neurons (Hunt et al., 2012; Parsadanian et al., 2006; Tsujino et al., 2000). All other clusters were classified as being in a transcriptionally “naive state,” and were comprised primarily of nuclei from naive mice (∼93% of nuclei in these clusters were from naive mice) with a median *Atf3* expression of 0. “Injured state” neurons express higher levels of all canonical DRG axonal injury-induced genes such as *Atf3*, *Sox11*, *Sprr1a*, *Flrt3* (Chandran et al., 2016; Costigan et al., 2002; LaCroix-Fralish et al., 2011; Perkins et al., 2014; Xiao et al., 2002) than “naive state” neurons (Figures 1E, 1G, two-tailed Student’s t-test, *P* < 0.001) and overlap with injury gene modules previously identified from bulk microarray studies (Chandran et al., 2016) (Figure S2D). It is notable that we still observe a small number of “naive state” neurons in mice who underwent SpNT (Figure 1D), consistent with the 5-10% of neurons not axotomized in this model. Several of the canonical injury-induced transcription factors are expressed within hours after injury, well before the full emergence of the “injured state,” raising the possibility that these transcription factors are involved in establishing the later transcriptional transformation of the neurons after injury (Figure 1G).

To test the accuracy of our injured versus non-injured neuron classification, we compared the percentage of neurons classified as injured in SpNT, a proximal injury model that causes axotomy of >90% of DRG neurons in the affected DRG (Shortland et al., 2006; Tsujino et al., 2000) and in ScNT, a more distal injury model that results in axotomy of ∼50% of the affected DRGs (Laedermann et al., 2014; Rigaud et al., 2008). Three days after axotomy, the injury classification identified 93.8% of neurons sequenced as “injured” after SpNT and 53.3% after ScNT (Figure 1H, S2E). Therefore, there is good agreement between the detection of axotomized neurons from the snRNA-seq analyses and those measured by *in vivo* anatomical labeling/tracing (Rigaud et al., 2008; Shortland et al., 2006). Interestingly, a few DRG neuronal nuclei from naive mice (mean 0.34%) were classified as being in an “injured state,” which may be explained by neurons injured from occult fight wounds that often occur in group-housed mice, and is consistent with the rare detection by *in situ* hybridization of *Atf3*+ neurons in naive mice (see Figure 2).

**Figure 2.**
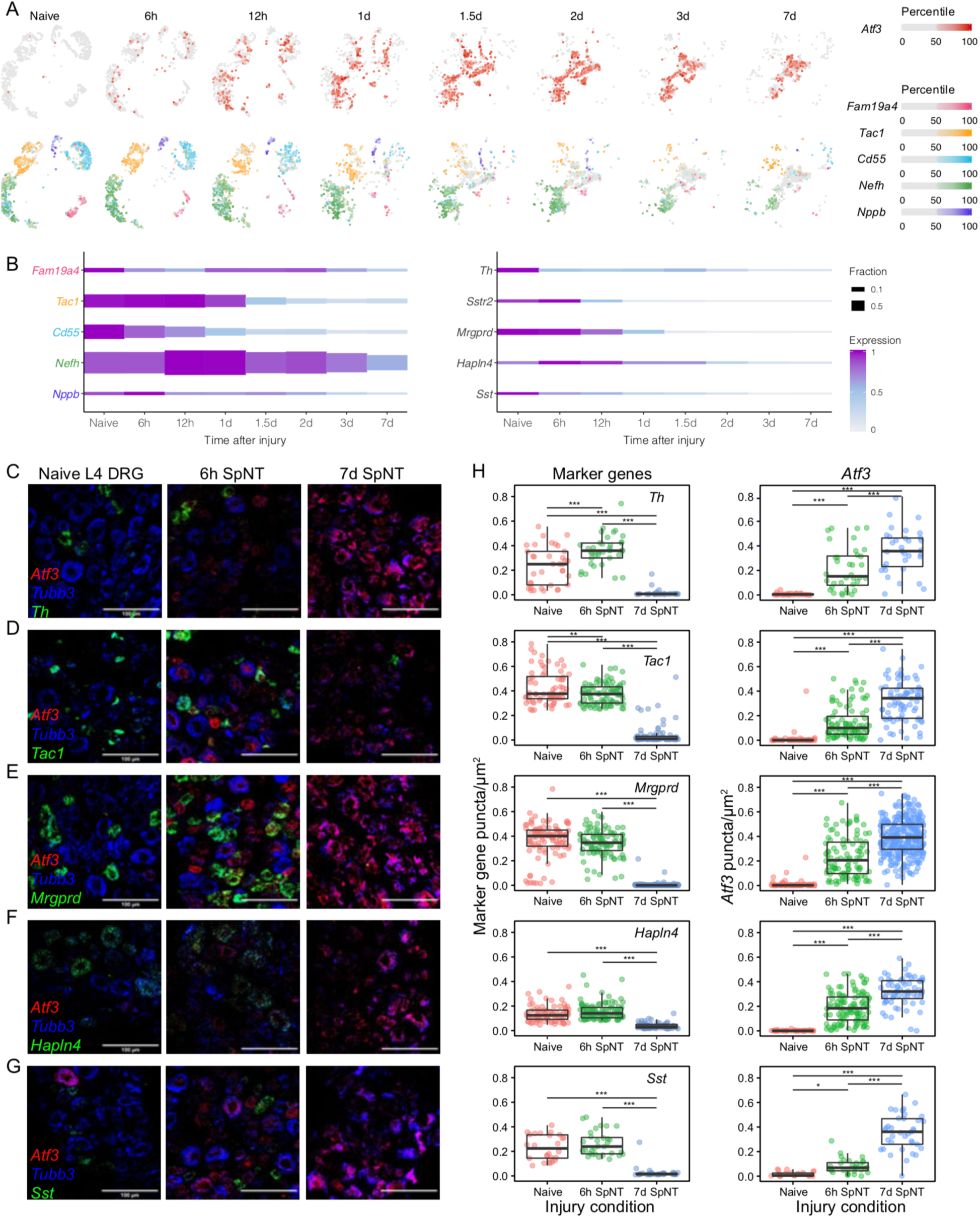
Loss of neuronal marker gene expression after DRG axonal injury. **(A)** UMAP plots displaying DRG neuronal subtypes after spinal nerve transection (SpNT). Nuclei are colored by *Atf3* (top) or by subtype-specific marker genes (bottom). For each gene, the color of nuclei represents the percentile of gene expression within SpNT neurons above the median (50^th^ percentile) of nuclei with > 0 counts of the corresponding gene; nuclei with expression below the median and no expression are colored gray. For cell-type-specific marker genes, 4.5% of nuclei that had expression above the median for multiple markers and their colors were overlaid. Time points were downsampled to the number of nuclei at the time point with the fewest number of nuclei sequenced (900 neuronal nuclei). Marker genes: *Atf3* (injury), *Fam19a4* (C-fiber low threshold mechanoreceptor), *Tac1* (peptidergic nociceptor), *Cd55* (non-peptidergic nociceptor), *Nefh* (*Nefh+* A-fiber low threshold mechanoreceptors), *Nppb* (*Sst+* pruriceptors). **(B)** Plot showing expression level of neuronal subtype-specific marker genes across neuronal nuclei and the fraction of naive or SpNT nuclei that express each gene (rows) over time. Fraction of nuclei is calculated as the number of nuclei expressing each gene (>0 counts) divided by the total number of nuclei at each time point. Expression at each time point is calculated as the mean scaled counts of a marker gene relative to the highest mean-scaled counts of that gene across time points. **(C-G)** Fluorescence *in situ* hybridization (FISH) images of L4 mouse DRGs stained with probes against *Atf3* (I-M, injury marker, red), *Tubb3* (I-M, neuronal marker, blue) and cell type markers: *Mrgprd* (C, green), *Hapln4* (D, green), *Tac1* (E, green), *Th* (F, green) or *Sst* (G, green). Representative sections from naive DRGs (left), DRGs 6 hours (middle) and 1 week (right) after SpNT are shown. **(H)** Quantification of *Atf3* and DRG neuronal subtype-specific marker gene expression from naive DRGs, DRGs 6 hours and 7 days after SpNT as measured by *in situ* hybridization (n = 3-6 L4 DRGs from different mice for each probe combination). Each dot on the boxplot represents gene expression within an individual cell, boxes indicate quartiles and whiskers are 1.5-times the interquartile range (Q1-Q3). The median is a black line inside each box. Significance testing by 1-way ANOVAs were all *P <* 0.001: *Th* (n = 36 [naive], 36 [6h], 33 [7d]), *F*(2, 102) = 74.70, *Atf3*(on *Th* slides), *F*(2, 102) = 52.87; *Tac1* (n = 68 [naive], 93 [6h], 78 [7d]), *F*(2, 236) = 332.33, *Atf3*(on *Tac1* slides), *F*(2, 236) = 112.56; *Mrgprd* (n = 100 [naive], 102 [6h], 308 [7d]), *F*(2, 507) = 1210.87, *Atf3*(on *Mrgprd* slides), *F*(2, 507) = 315.33; *Hapln4* (n = 80 [naive], 114 [6h], 64 [7d]), *F*(2, 255) = 85.52, *Atf3*(on *Hapln4* slides), *F*(2, 255) = 192.61; *Sst* (n = 26 [naive], 31 [6h], 37 [7d]), *F*(2, 91) = 82.98, *Atf3*(on *Sst* slides), *F*(2, 91) = 110.91; Tukey HSD post-hoc testing (***: p < 0.001, **: p < 0.01, *: p < 0.05).

### Classification of neuronal subtypes after axotomy

A primary goal of this study was to determine whether the intrinsic axonal injury transcriptional program differs between the distinct sensory neuronal subtypes and if these differences could inform differential phenotypes after injury. Efforts to address this question are complicated by the downregulation of the neuronal subtype-specific marker genes that classify neuronal subtypes that begins less than a day after axotomy (Figure 2A). Three to seven days after injury, expression of the marker genes used to classify neuronal subtypes was reduced by 65-97% compared to levels in naive DRGs, with a more pronounced downregulation of small diameter neuron marker genes (*e.g. Tac1, Mrgprd*) than those in large diameter neurons (e.g. *Nefh, Hapln4*) (Figure 2B). *In situ* hybridization for several neuronal subtype marker genes, including *Th, Tac1*, *Mrgprd, Hapln4, Sst* (Figures 2C-G) confirmed the significantly reduced marker gene expression. The coupling of marker gene downregulation with the profound changes in cluster identity after injury makes it difficult to classify injured neuronal subtypes, even if injury-induced genes are omitted when clustering (Figure S3A). To overcome this, we used multiple consecutive timepoints after SpNT to capture the transition between “naive” and “injured” states for each neuronal subtype. When neighboring time points after injury were co-clustered, residual cell-type-specific transcriptional signatures in injured nuclei led them to co-cluster with nuclei classified prior to marker gene downregulation. The defined subtypes were then projected onto the “unknown” injured nuclei with which they co-clustered (Figures 3A-B, S3B) (see methods). As a complementary informatic approach for classifying injured neuronal subtypes, we used a vector of injury-induced genes as a measurement of injury progression (see methods), and removed the variation in each gene that can be explained by the injury signal prior to clustering. Cell type assignments from the two approaches had 99% concordance for naive cell types and 91% for injured cell types (Figure S3C). To test the accuracy of the bioinformatic classification of neuronal subtypes after injury we performed lineage tracing of non-peptidergic (*Mrgprd*+) nociceptors after injury using *Mrgprd-Cre^ERT2^*;*Gcamp6f* reporter mice. SnRNA-seq of DRGs from injured and naive reporter mice identified reporter-positive nuclei in the same clusters as those classified informatically by pair-wise clustering and projection (estimated error = 2.93% in “injured state” nuclei and 1.88% in “naive state” nuclei, Figure S3D). The ability to classify neuronal subtypes at each time point after axonal injury (Figures 3B, S3E) provides an opportunity to characterize cell-type-specific molecular adaptions to axonal injury.

**Figure 3.**
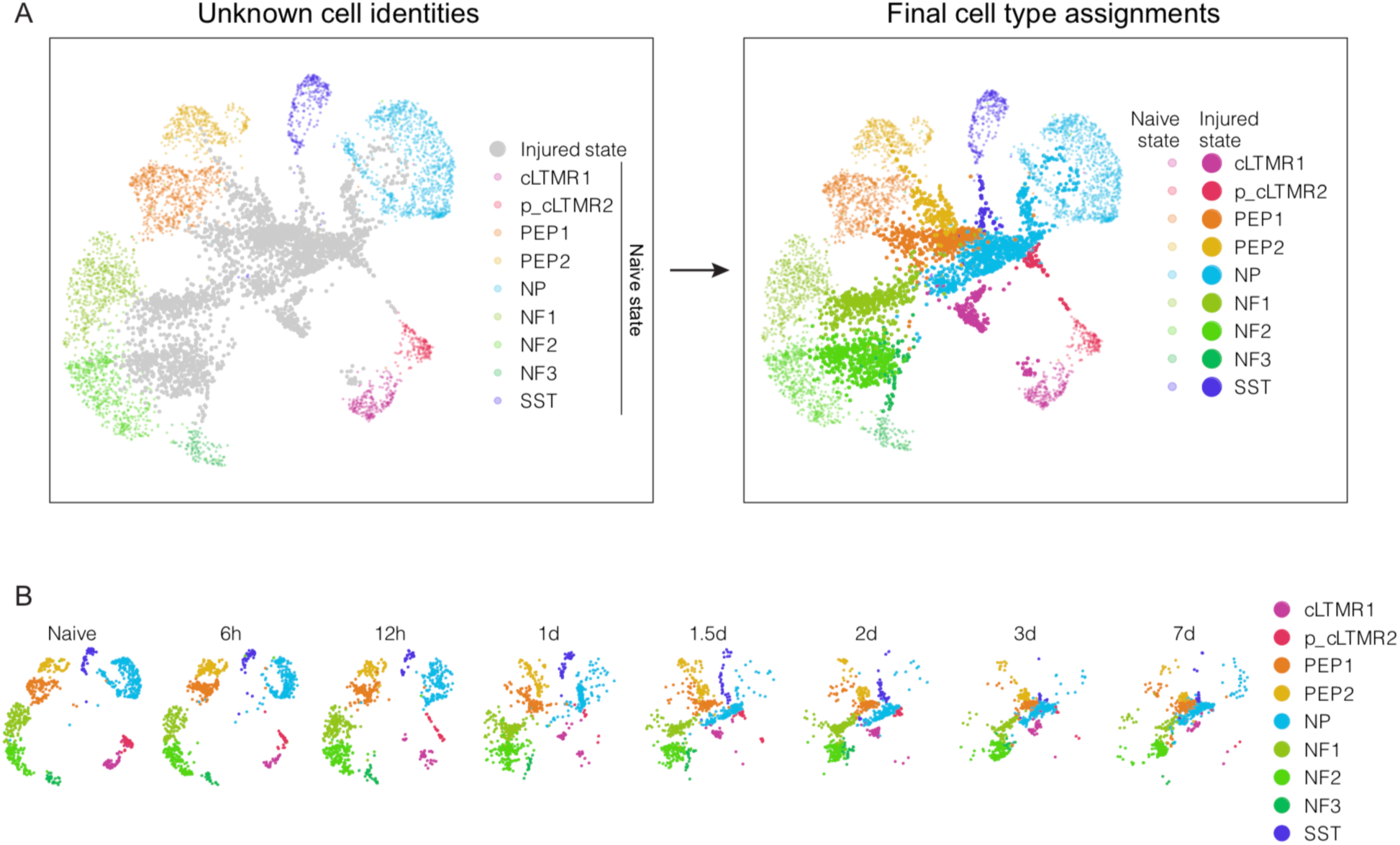
Classification of DRG neuronal subtypes after axotomy. **(A)** Classification of injured neuronal subtypes after spinal nerve transection (SpNT) by pair-wise clustering and projection. UMAP plots showing 7,000 naive and 7,000 SpNT neurons that were randomly sampled for purposes of visualization. Prior to pair-wise clustering and projection, neurons that are classified in the “naive state” are colored by their respective neuronal subtype, and neurons in the “injured state” are gray (left). After injured neuronal subtype classification by pair-wise clustering and projection, injured-state neurons (bold) are colored by their subtype (right). Naive-state neurons are also colored by their subtype (faint). **(B)** UMAP plots displaying the progression from naive to injured-state for each neuronal subtype after pair-wise projection and clustering. DRG neurons from naive and each time point after SpNT are shown (900 randomly-sampled neuronal nuclei per time point). Color represents neuronal subtype. cLTMR = C-fiber low threshold mechanoreceptor; PEP = peptidergic nociceptor; NP = non-peptidergic nociceptor; NF = *Nefh+* A-fiber low threshold mechanoreceptors; SST = *Sst+* pruriceptors.

### Characterization of cell-type-specific transcriptional responses to injury reveals a common program

After classifying the neuronal subtypes of axotomized neurons following SpNT, we performed differential gene expression analyses (defined as FDR<0.01 and log_2_FC>|1|) for each cell type and time point. For all DRG cell types except p_cLTMR2, the total number of genes significantly regulated by axotomy increased over time until 3 to 7 days after injury (Figure 4A), an effect that is observed when keeping the number of nuclei or UMI constant over time (Figure S4A). However, the rate of gene induction after injury varied across cell types (Figures 4A, S4B, Table S2). Small diameter neurons (e.g. NP and PEP) induce more genes at earlier time points than large diameter neurons (e.g. *Nefh*+ A-LTMRs) (Figures 4A, S4B), while Schwann cells induce very few genes after injury. The genes upregulated in each neuronal subtype in response to injury significantly overlap with those induced by injury in other neuronal subtypes, indicating a largely common neuronal response to injury (Figures 4B, S4C). Indeed, between 74-94% of genes induced in neuronal subtypes after injury are induced across multiple neuronal subtypes (Figure 4C). The genes that are upregulated in response to injury in p_cLTMR2 or glial subtypes are notably distinct from those that are commonly upregulated in the other neuronal subtypes (Figures 4B, S4C).

**Figure 4.**
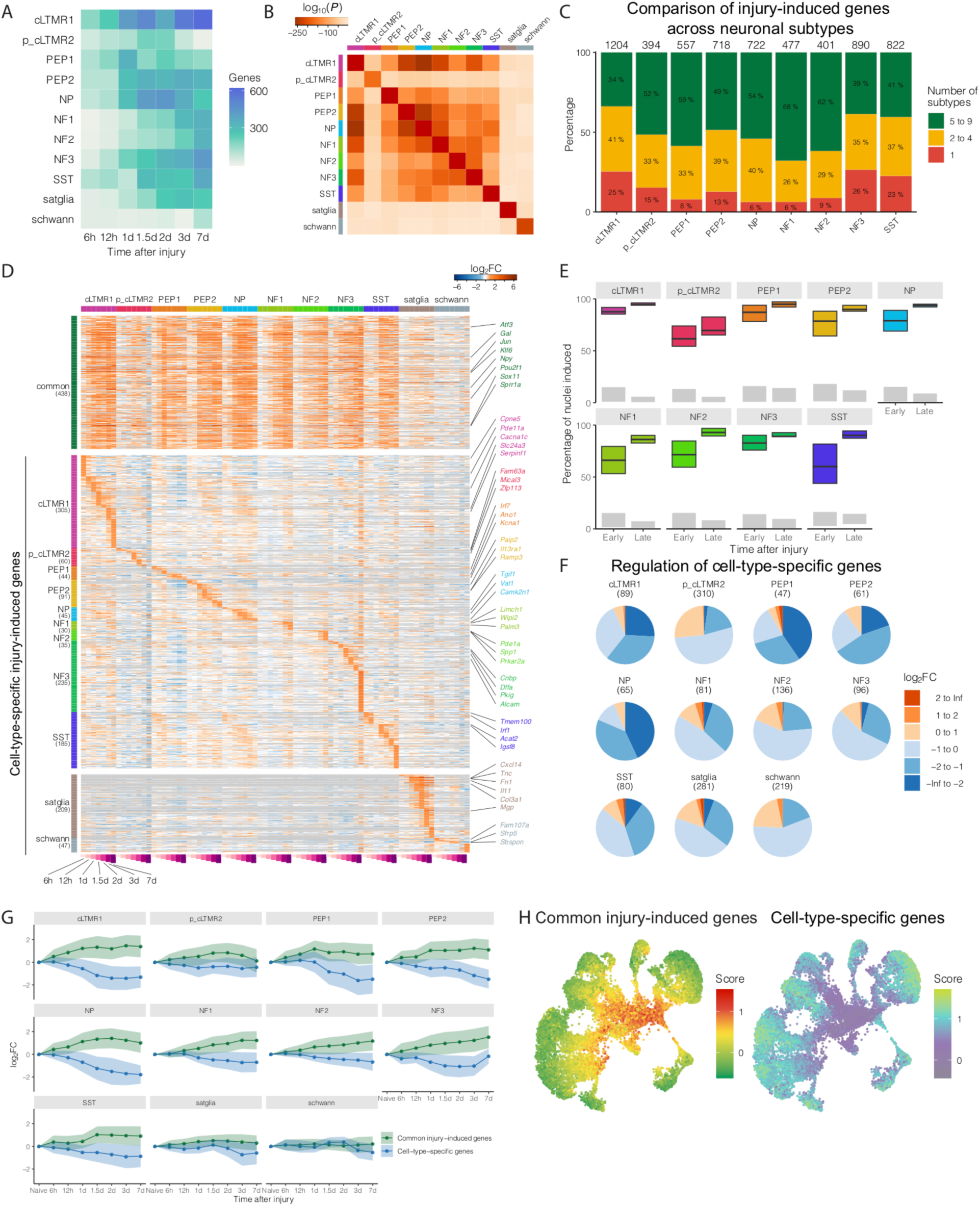
Characterization of cell-type-specific transcriptional responses to peripheral nerve injury. **(A)** Heatmap of the number of significant injury-induced genes for each cell type and time point after spinal nerve transection (SpNT) compared to their respective cell types in naive mice (FDR < 0.01, log_2_FC > 1). **(B)** Pair-wise comparison of overlapping injury-induced genes (FDR < 0.01, log_2_FC > 1; 3 and 7d after SpNT vs. naive) between the specified cell types after SpNT. Each square is colored by the *P*-value for the overlap between each comparison (hypergeometric test). Note that comparisons between the same gene list will always have 100% overlap but will have different hypergeometric p-values depending on list size. **(C)** Comparison of gene expression changes after SpNT compared to naive DRG neurons. Significantly upregulated genes after SpNT (FDR < 0.01, log_2_FC>1 SpNT vs. naive) in each neuronal subtype were aggregated across time points and compared to other neuronal subtypes to determine how many injury-induced genes are cell-type-specific (red), shared between 2-4 neuronal subtypes (yellow), or commonly shared between ≥ 5 other neuronal subtypes (green). Percentage of all significant injury-induced genes that are cell-type-specific, shared between 2-4 subtypes, or shared commonly across ≥ 5 subtypes are displayed on the bar plot. The total number of significantly-induced genes by SpNT in each subtype is shown on top of each bar. See Tables S3-4 for gene lists. **(D)** Heatmap displaying the change in expression over time after SpNT of regulated common genes (significantly upregulated by SpNT in ≥ 5 neuronal subtypes) and cell-type-specific genes (significantly upregulated by SpNT in 1 cell type) as defined in 4C. Genes are rows and cell types at each time point after SpNT are columns. Log_2_FC (SpNT vs. naive) for each time point and cell type is displayed. Genes are colored gray if they are not expressed in a cell type or at a certain time point. Select genes of interest are labeled. **(E)** Estimate of the fraction of nuclei that induce early injury-response genes (6h, 12h, and 1d) or late injury-response genes (3 and 7 days) after SpNT. A nucleus was classified as induced by injury if it expressed a threshold number of injury-response genes at the respective time point. Nuclei at 6h/12h/1d were classified using injury-induced genes from these time points, and 3d/7d nuclei were classified using a set of injury-induced genes at these time points. The boxes are defined by the fraction of injury-induced nuclei using different thresholds for the number of injury-response genes required for classification as induced by injury. The upper bar is the fraction of injury-induced nuclei using 2 injury genes/nucleus threshold, central line uses a 3 injury genes/nucleus threshold, and the lower bar uses a 4 injury genes/nucleus threshold. Grey rectangles show the fraction of nuclei from naive animals that are classified as induced by injury with the upper box boundary corresponding to a 2 injury gene/nucleus threshold and the lower boundary corresponding to a 4 injury gene/nucleus threshold. The set of injury-induced genes used to classify nuclei as “injury-induced” was chosen from the 10 common injury genes from Figure 4C with greatest fold-change between SpNT at 6h/12h/1d (early) or 3d/7d (late) and naive. An injury gene was counted towards the injury induction threshold in each nucleus if its Log_2_-normalized expression was > 90^th^ percentile of all nuclei of the same cell type from naive animals. **(F)** Regulation of cell-type-specific genes by SpNT in each cell type. Cell-type-specific genes are genes that are expressed significantly higher in one naive cell type compared to all other naive cell types (see methods). For each cell type, their respective cell-type-specific genes are grouped by log_2_FC after injury (SpNT at 3/7 days vs. naive within each subtype). Pie charts show the fraction of cell-type-specific genes within each neuronal subtype that are regulated by SpNT to the fold-change magnitude indicated. Total number of cell-type-specific genes for each subtype are shown in the header. **(G)** Line plots showing upregulation of common injury-induced genes (≥ 5 subtypes, from C) and downregulation of cell-type-specific genes (from F) for each cell type after SpNT. Each line represents the average log_2_FC of common injury-induced genes (green) or cell-type-specific genes (blue) over time. The ribbon represents standard deviation. **(H)** UMAP plots of 19,184 naive and SpNT DRG neurons. Nuclei are colored by either an aggregate injury score calculated from expression of 438 commonly induced genes after axotomy (left, see methods) or an aggregate cell-type-specificity score (right see methods). Aggregate cell-type-specificity scores are calculated for each neuronal type separately based on their respective cell-type-specific genes (see F). Higher scores indicate greater injury-induced or cell-type-specific gene expression. cLTMR = C-fiber low threshold mechanoreceptor; PEP = peptidergic nociceptor; NP = non-peptidergic nociceptor; NF = *Nefh+* A-fiber low threshold mechanoreceptors; SST = *Sst+* pruriceptors.

The common gene program induced after neuronal axotomy is enriched for genes involved in axon guidance, axonogenesis and regulation of cell migration (Figure S4D), and significantly overlaps (p = 8×10^-33^, hypergeometric test) with the injury-induced magenta gene module identified from a gene co-expression network analysis of bulk DRG microarray data (Chandran et al., 2016). This common neuronal transcriptional program includes genes previously identified in studies of axonal injury from bulk DRG tissue, such as *Atf3*, *Gal*, *Jun*, *Npy*, *Sox11* and *Sprr1a* (Figure 4D, Table S3) (Chandran et al., 2016; Costigan et al., 2002; LaCroix-Fralish et al., 2011; Perkins et al., 2014; Xiao et al., 2002). In addition to the common neuronal regeneration-associated program, there were also common changes in the expression of genes that impact neuronal excitability in all neuronal subtypes, including downregulation of multiple potassium channels and upregulation of the calcium channel*, Cacna2d1* (Figure S4E). These ion channel gene expression changes may contribute to the ectopic activity observed in injured neurons after axotomy (Liu et al., 2000; Patel et al., 2018; Serra et al., 2012; Tsantoulas and McMahon, 2014).

Single-nucleus profiling provides an opportunity to quantify the fraction of neurons within a DRG that induce the common transcriptional response to injury. We found that ∼50% percent of the neurons in each neuronal subtype show induction of the common injury gene program within hours after SpNT and this population increases to >90% 3-7 days after injury (Figure 4E), closely approximating the fraction of neurons physically axotomized in this model.

We also identified a smaller population of genes selectively induced only in specific neuronal subtypes after injury (Figures 4C-D, Table S4-5). These include genes involved in chloride homeostasis, cGMP signaling and integrin signaling pathways, some of which may contribute to cell-type-specific forms of axonal regeneration. For example, cLTMR1 neurons selectively induce *Serpinf1*, which has a pro-regenerative function in DRG neurons (Stevens et al., 2019) and NP neurons selectively induce *Vat1*, which also enhances DRG axon growth (Jia et al., 2018). Other cell-type-specific gene alterations may contribute to the neuropathic pain phenotype, as NF1 neurons selectively induce *Wipi2*, which is involved in autophagy in DRG neurons (Stavoe et al., 2019), a process argued to reduce the pain associated with sciatic nerve injury (Chen et al., 2018). PEP1 neurons selectively induce *Ano1*, which promotes pain hypersensitivity (Lee et al., 2014). These cell-type-specific gene expression changes in response to injury may also contribute to differences in axonal regeneration and/or excitability between distinct cell types.

Axonal regeneration and neuropathic pain appear to involve the participation of non-neuronal cells, such as the satellite glia which surround the somata of DRG neurons and the Schwann cells found around DRG axons (Gosselin et al., 2010; Jessen and Mirsky, 2016; Ji et al., 2016), but it has been difficult to isolate these cells and analyze their injury-induced gene expression changes (Jager et al., 2018). We find that while satellite glia induce a large number of genes in response to axonal injury, Schwann cells induce comparatively few genes (Figures 4A, 4D, Table S5). Several neuronal regeneration-associated genes, including *Atf3* and *Sox11*, are upregulated after axotomy in satellite glia and Schwann cells, but the induction is smaller in magnitude and more transient compared to neurons. A number of genes are selectively induced in glia but not in axotomized neurons. Satellite glia specifically induce tenascin C (*Tnc)* and fibronectin 1 *(Fn1),* both major components of the extracellular matrix, raising questions about the functional consequences of a potential change in the extracellular matrix in the immediate vicinity of neuronal cell bodies and their axons. Schwann cells uniquely induce complement C1q-like protein 3 (*C1ql3)* and *Tmem130*, a poorly characterized gene, although again the consequences of these changes require further study.

While many of the genes induced in sensory neurons after injury may promote regeneration-associated regrowth, the reprogramming of the injured neurons’ transcriptome extends beyond regeneration-associated genes and includes the downregulation of genes that define the identity and functional specialization of the neuron (Figure S4B).

### Profound transcriptional reprogramming after axotomy

To determine how cell-type-specific genes are regulated after axonal injury, we first compared gene expression in each neuronal cell type to that of all other neuronal subtypes to identify the cell-type-specific genes that are preferentially expressed in specific DRG cell types (FDR<0.01, log_2_FC>1, Table S6). More than 73% of the “cell-type-specific genes” in each DRG neuronal subtype were downregulated after axotomy (Figure 4F) and this downregulation occurred over the same time frame as the induction of the common neuronal injury genes (Figure 4G). By contrast, cell-type-specific markers in satellite glia, and Schwann cells were less affected by injury (Figure 4G). To determine whether the downregulation of cell-type-specific genes in neurons was specific to these genes or more broadly observed across the transcriptome, we compared the expression of cell-type-specific genes after injury to a set of randomly selected, expression-matched genes. We found that cell-type-specific genes were significantly more downregulated after injury than a set of randomly selected expression-matched genes in each neuronal subtype, except p_cLTMR2 (Figure S4F), indicating that there is a preferential downregulation of cell-type-specific genes in neurons after injury rather than a global redirection of transcriptional activators from all genes to injury response genes or a computational artifact of normalization. To quantify the extent of transcriptional reprogramming within each neuron, we generated scores using the average counts of common injury genes (injury score) or cell-type-specific genes (cell-type-specificity score). Projecting these scores onto each neuron in the UMAP plot accurately labeled the neurons as injured, with high injury scores and low cell-type-specificity scores (Figure 4H).

### Time course of injury-induced transcriptional reprogramming

To investigate the kinetics of injury-induced transcriptional reprogramming from initial injury through complete axonal regeneration, we turned to the sciatic nerve crush model in which full axonal regeneration, target reinnervation, and functional recovery occur within weeks to months after injury (Navarro et al., 1994; Vogelaar et al., 2004).

Similar to SpNT and ScNT, nuclei from mice who underwent sciatic crush injury began to adopt a transcriptional profile consistent with nerve injury within a day after sciatic crush, with injured nuclei displaying maximal injury scores and minimal cell-type-specificity scores 3-7 days after injury (Figure 5A). Similar injury-induced transcriptional changes were observed in both male and female DRG neurons after sciatic crush (Figure S5A). Between 2 weeks and 3 months following sciatic crush injury, the injured clusters of neurons gradually disappear (Figures 5A-B) in parallel with functional recovery (Figure S5B).

**Figure 5.**
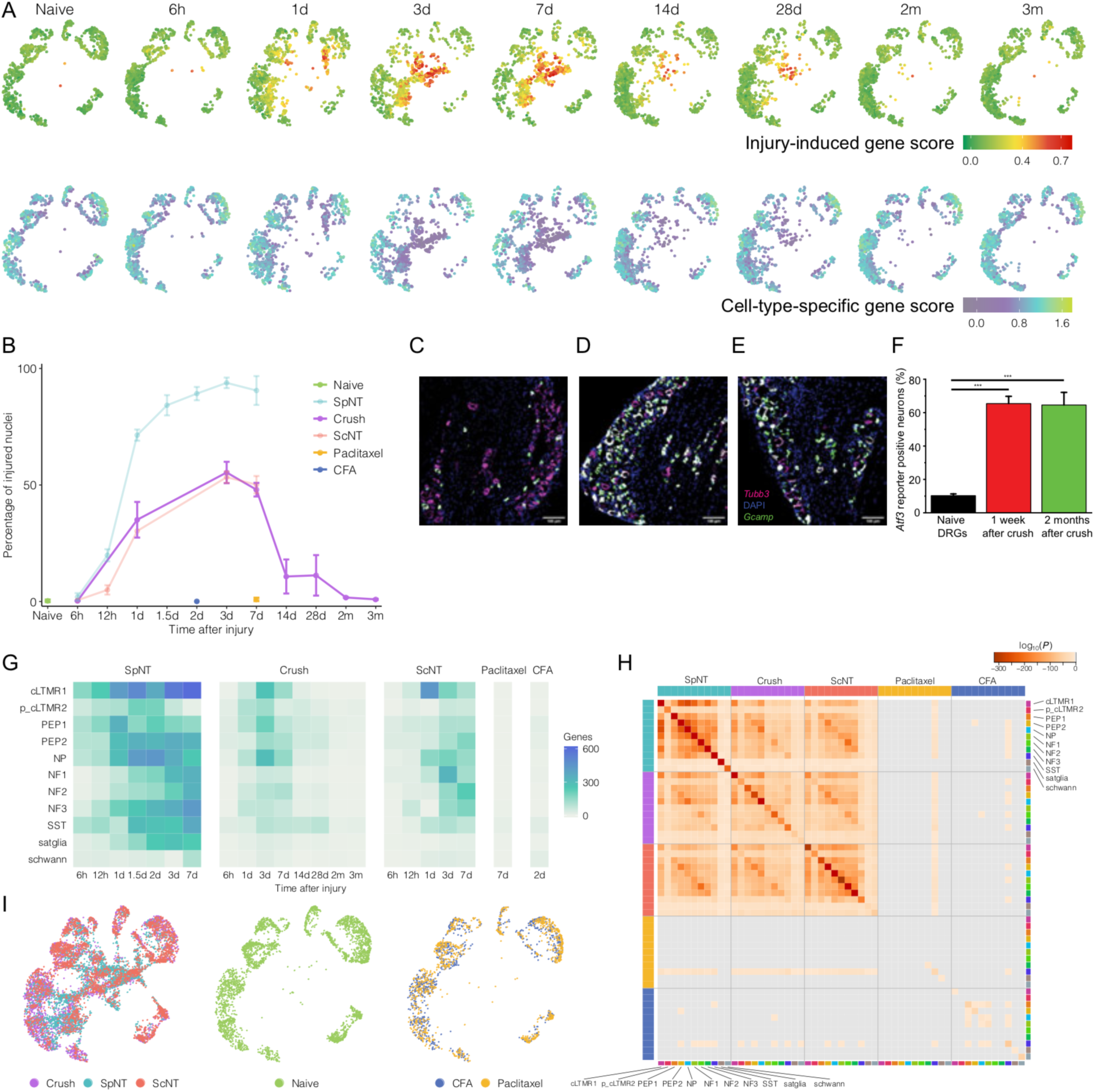
Transcriptional reprogramming of DRG neurons after axotomy. **(A)** UMAP plots displaying DRG neurons from either naive mice or mice who received sciatic nerve crush followed by the indicated amount of time prior to harvesting. Each time point down is sampled to the number of nuclei at which the fewest number of nuclei were sequenced (1000 neuronal nuclei). Nuclei are colored by the common injury score (top) or cell-type-specificity score (bottom) as in Fig 4H. Higher scores indicate greater injury-induced or cell-type-specific gene expression. **(B)** Percentage of naive, spinal nerve transection (SpNT), sciatic crush, sciatic nerve transection (ScNT), paclitaxel-treated, Complete Freund’s Adjuvant-treated (CFA) neuronal nuclei that are classified as in the “injured state” at each time point after the respective injury. Colors represent injury models; naive, crush, paclitaxel, and CFA are bolded, SpNT and ScNT are faded. **(C-E)** Fluorescence *in situ* hybridization (FISH) images of ipsilateral L4 *Atf3-Cre^ERT2^;Gcamp6f* DRG sections from a naive mouse (C), 1 week after sciatic crush (D) and 2 months after sciatic crush (E). Sections stained for the neuronal marker *Tubb3* (magenta), DAPI (blue) and the reporter, *Gcamp6* (green). The *Atf3*-driven *Gcamp6* reporter is upregulated after sciatic crush and persists for months after injury. **(F)** Quantification of *Gcamp6* reporter gene expression in L4 *Atf3-Cre^ERT2^;Gcamp6f* DRGs after sciatic crush measured by FISH. N = 3-4 DRG sections from different mice per group, one-way ANOVA, *F*(2, 8) = 37.4, *P* = 8.7×10^-5^. Sciatic nerve crush injury causes an increase in *Gcamp* reporter positive neurons 1 week after crush (Bonferroni post-hoc, *P* = 2.9 x 10^-4^), which persists for two months after sciatic crush injury (Bonferroni post-hoc, *P* = 1.9 x 10^-4^). **(G)** Heatmap of the number of significant injury-induced genes for each cell type and time point after SpNT, sciatic crush, ScNT, paclitaxel, or CFA compared to their respective cell types in naive mice (FDR < 0.01, log_2_FC > 1). **(H)** Pair-wise comparison of overlapping injury-induced genes between the specified cell types 3/7 days after SpNT, sciatic crush, ScNT, or paclitaxel or 2 days after CFA (FDR < 0.01, log_2_FC > 1, compared to naive nuclei of the respective cell type). Each square is colored by the *P*-value for the overlap between each comparison (hypergeometric test). Note that comparisons between the same gene list will always have 100% overlap but will have different hypergeometric p-values depending on list size. **(I)** UMAP plots show neuronal nuclei after different injury models (left, 3,000 nuclei randomly sampled equally from crush, SpNT, ScNT [total = 9,000 nuclei]; middle, 3000 nuclei randomly sampled from naive; right, 1,000 nuclei randomly sampled from paclitaxel and CFA [total = 2,000 nuclei]). Each nucleus is colored by the injury model to which it was exposed. cLTMR = C-fiber low threshold mechanoreceptor; PEP = peptidergic nociceptor; NP = non-peptidergic nociceptor; NF = *Nefh+* A-fiber low threshold mechanoreceptors; SST = *Sst+* pruriceptors.

The reduction in the number of “injured state” neurons 2-3 months after crush injury could be explained either by the reversal of their transcriptional reprogramming due to successful regeneration, or by the selective cell death of this neuronal population, both of which have been suggested as possibilities in the literature (Hart et al., 2002; Kataoka et al., 2007; Tandrup et al., 2000). To test the latter possibility, we generated an injury reporter mouse (*Atf3-Cre^ERT2^;Gcamp6f*) in which *Atf3* induction drives Cre-dependent expression of the *Gcamp6f* reporter gene (Figure S5C). This reporter efficiently marks injured *Atf3*+ DRG neurons 1 week after sciatic crush injury (Figure S5D-F). The percentage of reporter positive neurons was unchanged from 1 week to two months after crush, when the injury program has disappeared (Figures 5C-F), indicating that injured neurons do not die but rather return to their naive transcriptional profiles. This result is consistent with previous studies which reported minimal to no DRG neuron death after sciatic crush in rodents (Swett et al., 1995). Therefore, injury-induced transcriptional reprogramming reverses if axonal regeneration and reinnervation is complete.

Because sciatic crush, like ScNT, only results in the physical injury of ∼50% of L3-5 DRG axons (Chang and Namgung, 2013), there is a mixture of neurons with injured or uninjured axons in these ganglia. This can be observed both in the UMAP plots 3 and 7 days after sciatic crush (Figure 5A) as well as in the percentage of nuclei within clusters classified as injured (Figure 5B). To identify whether injury-induced gene expression also occurs in unaxotomized neurons, we performed differential expression analysis between neurons classified as uninjured in animals who underwent sciatic crush and the same cell type in naive animals. We found a transient induction of some common injury-induced genes in uninjured neurons after sciatic crush, but the magnitude of these changes was very small in comparison to injured neurons from the same mice (Figures S5G-H). The transient induction of common injury response genes like *Atf3* or *Nts* could be due to surgical injury-induced inflammation and stress, or paracrine signaling between injured and non-injured neurons (Berta et al., 2017; Fukuoka et al., 2012).

Cell-type-specific marker genes were downregulated in injured neurons after sciatic crush (Figures 5A, S5I-N), but we could assign neuronal subtypes to all nuclei, including those injured by crush or ScNT, because they co-clustered with the classified SpNT injured neuronal subtypes (Figure S6A). Differential gene expression analysis comparing injured neuronal subtypes after sciatic crush or ScNT at each time point after injury with their respective naive subtypes, revealed a peak of gene induction 3-7 days after injury, similar to that observed for SpNT (Figure 5G). Moreover, there is significant overlap between the genes induced in a given cell type across all the axotomy models (Figure 5H, Figure S6B-C), indicating that a common transcriptional program is induced by axotomy in most peripheral sensory neuron subtypes regardless of injury location (proximal or distal) or the fraction of injured DRG neurons. It should be noted that the small number of gene expression changes in crush and ScNT compared to SpNT was primarily a consequence of the smaller number of axotomized neurons in the distal injury models than SpNT. The extent and composition of the gene expression changes were quite similar across the distal and proximal axotomy models when specifically comparing neurons in the “injured state” with their naive controls (Figure S6D).

### Inflammatory and chemotherapy-induced transcriptional changes

The high correlation between the transcriptional programs induced by three different physical axotomy models led us to test whether the same reprogramming is engaged in a model of acute painful peripheral neuropathy caused by paclitaxel treatment and an inflammatory pain model produced by intraplantar injection of Complete Freund’s Adjuvant (CFA). Paclitaxel treatment causes mechanical allodynia 1 week after treatment (Figure S6E) and causes peripheral neuropathy by 4 weeks after treatment (Toma et al., 2017; Velasco and Bruna, 2015), while injection of CFA into the hindpaw leads to inflammation and mechanical allodynia within 24 hours after treatment (Figure S6F) (Ghasemlou et al., 2015; Jaggi et al., 2011). SnRNAseq was performed on L3-5 DRGs from mice treated with paclitaxel or CFA and compared with naive and axotomized mice. Over 99% of neurons from paclitaxel-treated mice and CFA-treated mice clustered together with naive nuclei (Figure 5I). Cell-type-specific differential expression analysis between paclitaxel- or CFA-treated and naive-treated mice displayed few statistically significant genes (Figure 5G, S6G) and those which were significantly regulated had little overlap with axotomy models (Figure 5H, I). The presence of pain is thus independent of injury-induced transcriptional reprogramming in DRG neurons.

### Transcriptional mechanisms underlying injury-induced transcriptional reprogramming of sensory neurons

Transcription factors that mediate the injury-induced transcriptional reprogramming must be induced very rapidly (≤ 1 day) after injury and have consensus DNA binding sites enriched in the set of genes that are induced several days after injury when the injury score plateaus (Figure 6A). Within hours of injury, we identified 24 transcription factors commonly upregulated after SpNT across neuronal subtypes and whose target gene regulation is enriched in DRG nuclei (Figure 6B, see methods). Over half of these 24 transcription factors have been previously detected after axonal injury (e.g. *Atf3, Jun, Jund*) (Chandran et al., 2016; Herdegen et al., 1992; Mahar and Cavalli, 2018; Patodia and Raivich, 2012; Tsujino et al., 2000). After identifying transcription factor binding motifs that are significantly enriched compared to all motifs in the set of commonly-induced injury genes, we ranked each early injury-induced transcription factor by the number of these enriched motifs they bind. We observed that the activating protein 1 (AP-1) family members such *Jun, Jund,* and *Fosl2* as well as *Atf3* were associated with the highest number of enriched motifs, an effect that was not observed for motifs identified in random sets of genes (Figure 6C, permutation test, *P* < 0.001). We chose to focus on *Atf3* because it is the transcription factor most strongly upregulated within hours after injury across neuronal subtypes whose consensus binding motifs are also enriched in the common set of genes that are upregulated after injury compared to naive neurons. Indeed, there is a strong and significant correlation between the level of *Atf3* mRNA and its predicted activity on its target genes in individual neurons (Figures 6D, S7A, Pearson’s r = 0.48, permutation test, *P* < 0.001), indicating that Atf3 is likely to play an important role in injury-induced transcriptional reprogramming.

**Figure 6.**
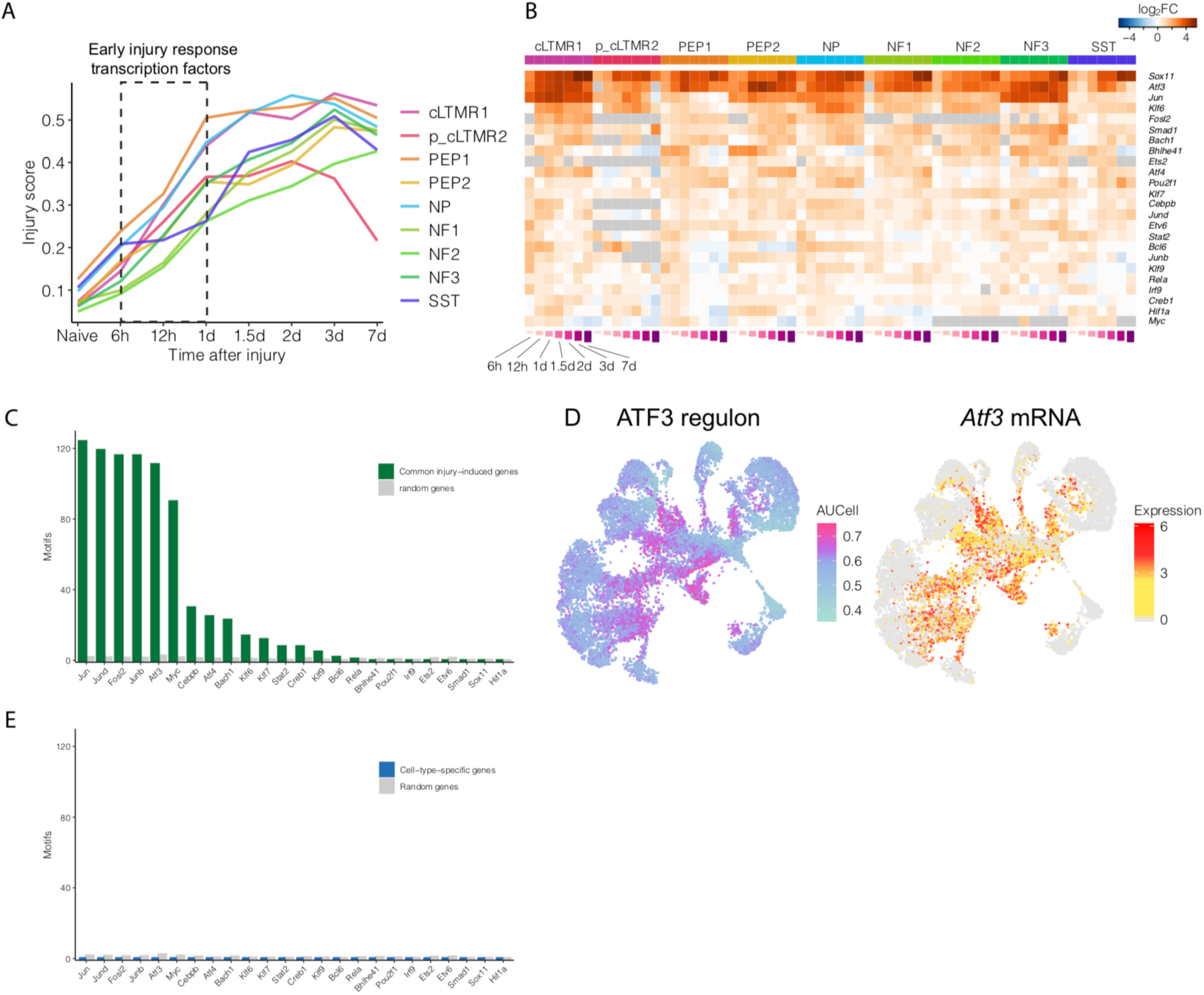
Induction of a common set of transcription factors across sensory neuronal subtypes after axotomy. **(A)** Mean common injury score for specific neuronal subtypes at each spinal nerve transection (SpNT) time point. Dotted box highlights the time points at which transcription factors that are significantly upregulated (FDR < 0.01, log_2_FC > 0.5, SpNT vs. naive) early after injury were identified. **(B)** Heatmap of 24 transcription factors (rows) that are significantly induced ≤ 1 day after SpNT (FDR < 0.01, log_2_FC > 0.5) in ≥ 5 neuronal subtypes. Heatmap is colored by log_2_FC (SpNT vs. naive) for each neuronal subtype and injury time point (columns). **(C)** Bar graph showing the number of significantly-enriched transcription factor binding motifs in 438 common injury-induced genes to which each early injury-induced transcription factor binds. Gray bars show the average number of transcription factor binding motifs enriched in 1000 sets of 438 randomly-selected expressed genes. **(D)** UMAP of neuronal nuclei from naive and SpNT mice colored by their degree of ATF3 regulon enrichment (left, AUCell score, see methods) or log_2_-normalized expression of *Atf3* (right). Nuclei with no *Atf3* expression colored gray. **(E)** Bar graph showing the number of significantly-enriched motifs in 1240 cell-type-specific genes that each early injury-induced transcription factor binds (green bars). Gray bars show the average number of transcription factor binding motifs enriched across 1000 sets of 1240 randomly-selected expressed genes. cLTMR = C-fiber low threshold mechanoreceptor; PEP = peptidergic nociceptor; NP = non-peptidergic nociceptor; NF = *Nefh+* A-fiber low threshold mechanoreceptors; SST = *Sst+* pruriceptors.

While many of the injury-induced transcription factors are known to have both transcriptional activating and repressing roles (Aguilera et al., 2011; Renthal et al., 2008), the absence of their motif enrichment in the set of cell-type-specific genes compared to random sets of genes (Figure 6E) suggests alternative mechanisms are likely to contribute to the downregulation of cell-type-specific genes after injury (see Discussion).

To determine if *Atf3* in sensory neurons is necessary for injury-induced transcriptional reprogramming and sensory neuronal regeneration after injury, we generated a floxed *Atf3* mouse and crossed it with *Vglut2-Cre* mice (Figures 7A), resulting in a conditional knockout (cKO) of *Atf3* from >95% of sensory neurons (*Atf3* WT: 89 ± 1% of DRG neurons 1 week after SpNT are ATF3+ Nissl+; *Atf3* cKO: 4 ± 2% of DRG neurons 1 week after SpNT are ATF3+ Nissl+; n=4 DRG sections, p<0.001, two-tailed Student’s t-test) (Figure 7B). Consistent with a role for *Atf3* in axonal regeneration (Gey et al., 2016; Jing et al., 2012; Seijffers et al., 2006), the deletion of *Atf3* in sensory neurons resulted in a significant delay in functional sensory recovery after sciatic crush injury (Figure 7C), an effect that we also observed using a tamoxifen-inducible cKO approach in the adult mouse (Figures S7B-C).

**Figure 7.**
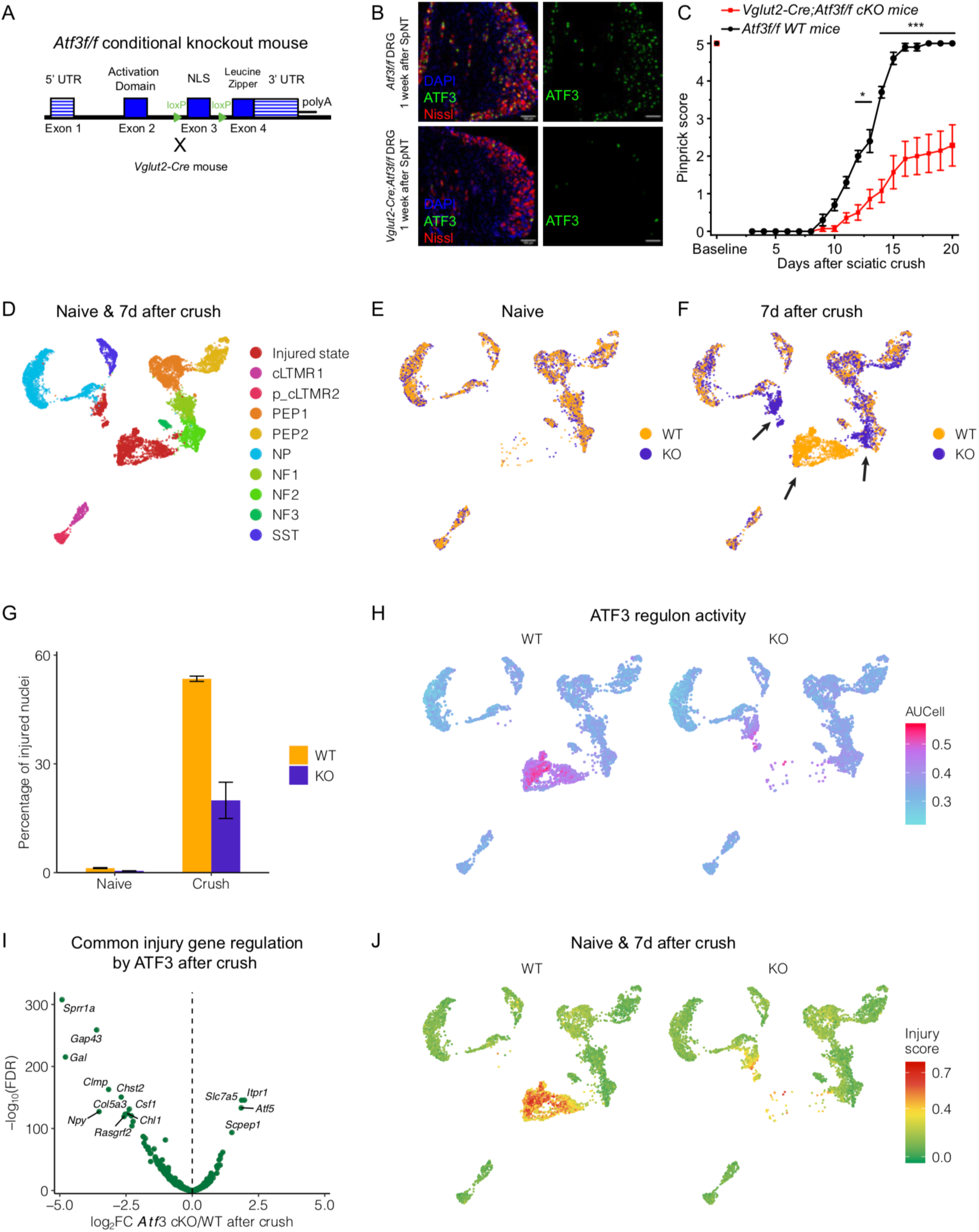
*Atf3* is required for axon regeneration. **(A)** Strategy used to create *Atf3* conditional knockout (cKO) mice. Transgenic mice carrying a floxed allele of *Atf3*, where loxP sites surround exon 3 (nuclear localization element) of *Atf3* were generated. These mice were crossed with *Vglut2-Cre* mice, which express *Cre* in >95% of sensory neurons (Kupari et al., 2019). **(B)** Representative images of *Vglut2-Cre;Atf3f/f* (cKO, bottom) or *Atf3f/f* (WT, top) 1 week after SpNT injury. DRGs are stained with antibodies against ATF3 (green), DAPI (blue) and neurons are counterstained with Nissl. There is a clear loss of ATF3 staining in the cKO compared to the WT. **(C)** Recovery of sensory function as measured by the pinprick assay in *Atf3f/f* (WT) and *Vglut2-Cre;Atf3f/f* (*Atf3* cKO) mice after sciatic nerve crush. Sciatic nerve crush causes a loss of sensory responses in the ipsilateral hindpaw, followed by a recovery over time associated with sensory neuron regeneration. The pinprick responses of *Atf3f/f* WT mice (n=10, black line) recover to baseline within 15 days after sciatic nerve crush (1-way repeated measures within subjects ANOVA, lower bound *F*(1, 9) = 388, *P* = 1.0×10^-8^). The pinprick responses of *Atf3* cKO mice (n=14, red line) show a significant delay in the time course of sensory function recovery (2-way repeated measures between subjects ANOVA, *F*(1, 22) = 33.7, *P* = 7.7×10^-6^, Bonferroni post-hoc, * *P* < 0.05, *** *P* < 0.001), suggesting a slower rate of sensory neuron regeneration. **(D)** UMAP plot displaying 6,410 WT and 5,601 *Atf3* cKO DRG neurons from naive mice and mice 7d after sciatic crush. Neurons are colored by their neuronal subtype. **(E)** UMAP plot displaying 2,653 WT and 2,489 *Atf3* cKO DRG neurons from naive mice. Neurons are colored by genotype. **(F)** UMAP plot displaying 3,487 WT and 3,112 *Atf3* cKO DRG neurons from mice 7d after sciatic crush. Neurons are colored by genotype. Arrows point to novel neuronal clusters observed in the sciatic nerve crush samples. **(G)** Bar plot indicating the percent of nuclei classified as in the “injured state” in each condition (naive or 7d after crush) and genotype (WT or *Atf3* cKO). There is a significant reduction in the number of “injured state” neurons in *Atf3* cKO compared to WT 7d after sciatic crush (one-way ANOVA: *F*(3, 4) = 192.96, *P* < 0.001; Tukey HSD post-hoc testing *P* > 0.05 for naive cKO vs. naive WT, *P* < 0.01 for naive cKO vs crush cKO, and naive WT vs. crush cKO, *P* < 0.001 all other pair-wise comparisons). **(H)** UMAP of WT (left) or *Atf3* cKO (right) DRG neuronal nuclei from naive mice and mice 7d after sciatic crush colored by their degree of ATF3 regulon enrichment (AUCell score, see methods). **(I)** Volcano plot displaying differential expression of 436 common injury-induced genes between *Atf3* cKO and WT neuronal nuclei that are classified as in the “injured state.” The common injury-induced genes are obtained from the 438 genes described in Figures 4C-D; 2 genes were not expressed in *Atf3* WT and cKO mice. **(J)** UMAP plots displaying 6,410 WT (left) and 5,601 *Atf3* cKO (right) neuronal nuclei from naive mice and mice 7d after sciatic crush. Neurons are colored by the common injury score (see methods). cLTMR = C-fiber low threshold mechanoreceptor; PEP = peptidergic nociceptor; NP = non-peptidergic nociceptor; NF = *Nefh+* A-fiber low threshold mechanoreceptors; SST = *Sst+* pruriceptors.

To determine if *Atf3* is required for injury-induced transcriptional reprogramming, we performed snRNA-seq on *Atf3f/f* (WT) and *Vglut2-Cre;Atf3f/f* cKO DRGs that are either naive or 7 days after sciatic nerve crush. We clustered WT and *Atf3* cKO neuronal nuclei together and found that the naive neuronal subtypes from these mice cluster together and express the same subtype-specific marker genes (Figures 7D-E, S7D), indicating a high degree of transcriptional similarity between naive WT and *Atf3* cKO DRG neurons. To compare transcriptional responses between WT and *Atf3* cKO after sciatic crush, we first identified the clusters of neurons from these mice that have high common injury scores and exhibit the “injured” transcriptional state (Figures 7D, S7E). Consistent with a central role of *Atf3* in driving injury-induced transcriptional reprogramming, we observed significantly fewer *Atf3* cKO DRG neurons in the “injured” transcriptional state 7 days after sciatic nerve crush than WT neurons (Figures 7E-G), an effect that is not explained by neuronal cell loss (Figure S7F). The attenuation of injury-induced transcriptional reprogramming in *Atf3* cKO DRG neurons is associated with significantly less putative *Atf3* target gene induction than is observed in WT neurons after injury (Figures 7H, S7G). Moreover, the clusters of “injured state” *Atf3* cKO neurons express most common injury genes at significantly lower levels than “injured state” WT neurons (e.g. *Sprr1a, Gal, Gap43*) (Figures 7I-J), which likely contributes to the axonal regeneration deficit in these mice (Schmid et al., 2014; Woolf et al., 1990). Together, these findings implicate *Atf3*, and possibly other transcription factors that are induced rapidly after injury, in the transcriptional reprogramming and subsequent axonal regeneration that occurs after nerve injury.

## Discussion

Peripheral nerve injury initiates a cascade of events that result in the conversion of sensory neurons from a non-growing to an active regenerating state. While previous studies have generated a number of mechanistic insights into this process, they have largely relied on bulk DRG gene expression studies which mask heterogeneous response to axonal injury (Chandran et al., 2016; Costigan et al., 2002; Xiao et al., 2002) or the dissociation or sorting of a small number of DRG neurons (Chiu et al., 2014; Sakuma et al., 2016; Usoskin et al., 2015; Zeisel et al., 2018), a process which itself induces many injury-related transcriptional changes (Hrvatin et al., 2018; Lacar et al., 2016; Wu et al., 2017). To avoid these confounders, and to identify cell type specific changes, we used snRNA-seq to generate a DRG cell atlas, with gene expression profiles of 107,541 DRG nuclei derived from naive and injured mice. Using these data, we interrogated the transcriptional mechanisms by which injury initiates axonal regeneration and may also contribute to neuropathic pain (Cattin and Lloyd, 2016; Ji et al., 2016).

One of the most dramatic findings in our study is that peripheral axonal injury results in a profound transcriptional reprogramming of DRG neurons, one involving both the induction of a common set of injury-response genes across neuronal subtypes and the coincident downregulation of their cell-type-specific genes. This transcriptional reprogramming is reversible, as the transcriptional states of injured neuronal nuclei return to their naive states within weeks, when the axons successfully reinnervate their targets, (Figure S5B) (Navarro et al., 1994; Vogelaar et al., 2004). An analogous process also occurs in the trigeminal ganglion after infraorbital nerve injury (Nguyen et al., 2019). Injury-induced transcriptional reprogramming leads to a new transcriptional state in which neuronal subtypes become difficult to distinguish because of the upregulation of a common set of injury-response genes and the attenuation of cell-type-specific genes after injury. However, we were able to classify each injured neuronal subtype by developing an informatic approach, validated by lineage tracing, that extracted the subtle cell-type-specific gene expression signatures that remained after injury. This ability to classify injured neuronal subtypes then enabled us to determine which components of the nerve injury response are common or cell-type-specific. While cell-type-specific gene expression changes do manifest after axonal injury (e.g. p_cLTMR2) and may contribute to distinct injury responses between cell types (Figure 4D, Table S4), the most striking observation was that the majority of injury-induced gene expression changes are common across neuronal subtypes and that the differences in gene expression between highly specialized DRG neuronal subtypes are lost.

The profound transcriptional reprogramming that occurs after axotomy is associated with the rapid induction of transcription factors within hours after injury. Many of these transcription factors (e.g. *Atf3, Jun, Klf6*) have their consensus DNA binding sites enriched in regions upstream of the genes induced days later after axotomy. *Atf3* has previously been implicated in peripheral neuron regeneration (Gey et al., 2016; Seijffers et al., 2007; Tsujino et al., 2000), but the mechanisms by which *Atf3* function have remained unclear. Consistent with these prior reports, we observed that *Atf3* was one of the most prominent neuronal injury-induced transcription factors identified in our study, as defined by its rapid induction after injury and the extent of its motif enrichment in the pool of injury-induced genes days after injury (Figure 6B). We also found that conditional deletion of *Atf3* in sensory neurons resulted in a substantial impairment of sciatic nerve regeneration and limited the ability of DRG neurons to activate the common neuronal injury gene program (Figures 7C, 7G, 7I-J). *Atf3* is likely to act in concert with other injury-induced transcription factors, such as *Jun* and *Klf6* (Chandran et al., 2016; Raivich et al., 2004), to produce the transcriptional and functional metamorphosis from mature neurons devoted to sensory transduction to injured neurons devoted to axonal growth and target re-innervation, which is also accompanied by pain-producing ectopic neuronal activity.

It has been previously hypothesized that axonal injury may reactivate an embryonic development program to drive regeneration (Harel and Strittmatter, 2006; Lisi et al., 2017). We do observe a limited induction of genes after injury that are also regulated during embryonic DRG development (Figure S7H, Table S8), but there is no statistically significant overlap between these two programs. Rather, many of the injury-induced transcription factors are related to the families of transcription factors capable of reprogramming differentiated cells into induced pluripotent stem cells or in the transdifferentiation of a mature cell into a distinct other cell type. This overlap suggests that strong environmental stimuli, such as axonal injury, may invoke transcriptional reprogramming mechanisms similar to those required to convert cells from one transcriptional identity to another, in order to change the primary function of somatosensory neurons from sensory transduction to axonal regeneration (Duan et al., 2019; Ronquist et al., 2017). Unlike stem cell reprogramming, however, injury-induced reprogramming is self-limited, only affecting the cell’s transcriptional state until axonal regeneration is complete. The mechanisms governing the timing and mechanisms of the deactivation of injury-induced transcriptional reprogramming will be the subject of future investigations.

While *Atf3* is a major driver of the common injury gene program and there are fewer neurons in the “injured state” after axotomy in *Atf3* cKO compared to WT (Figure 7G), *Atf3* binding sites are not enriched in the cell-type-specific genes that are downregulated after injury (Figure 6E). Thus, the downregulation of cell-type-specific genes after injury may be an indirect consequence of *Atf3* or of another transcription factor that is rapidly induced after injury and/or the redirection of RNA polymerase/co-activators from cell-type-specific genes to the common injury response genes. The downregulation of cell-type-specific genes after injury is likely to have functional implications, as many of these downregulated genes are ion channels involved in maintaining neuronal excitability (Figure S4E). For example, there is a broad downregulation of voltage-gated potassium channels after peripheral axotomy, which has been reported previously in bulk gene expression studies (Bangash et al., 2018; Chandran et al., 2016; Tsantoulas and McMahon, 2014) and this is associated with the neuronal hyperexcitability linked to injury-induced neuropathic pain (Colloca et al., 2017; Haroutounian et al., 2014; Serra et al., 2012).

Non-neuronal cells such as satellite glia and Schwann cells do not exhibit the same massive transcriptional reprogramming after nerve injury that sensory neurons do, but several transcription factors (e.g. *Srebf1* and *Nr3c1*) are induced after injury and have consensus binding sites enriched in the injury-induced genes in these cell types (Figure S7I). Paracrine signaling from injured neurons must produce these changes but interestingly our data indicate that non-injured neurons show only small and transient alterations. Similarly, we did not observe the same magnitude of injury-induced transcriptional reprogramming genes in non-axotomy models such as paclitaxel-induced painful neuropathy or CFA-induced inflammatory pain, at least not at the time points we investigated. These findings are consistent with observations from bulk gene expression studies (Bangash et al., 2018; Zhang and Dougherty, 2014) and support the hypothesis that distinct mechanisms are likely to drive nociceptor sensitization in these pain models.

We expect that single-cell sensory neuron atlases from both mice and humans will catalyze the identification of novel therapeutic targets for nerve repair and/or pain. Towards this goal, we have created an online resource at www.painseq.com which enables facile access to and visualization of the snRNA-seq datasets presented and analyzed in this study. This resource can be used to further explore the many gene expression changes that occur in response to nerve injury, paclitaxel-induced neuropathy, or inflammatory pain in animal models of these conditions.

## Supporting information

High-resolution Figures 1-7

High-resolution Figures S1-S7

## Acknowledgements

We would like to thank Michael Tetreault, Daniel Taub, and Nick Andrews for assistance with behavioral experiments; the Harvard Single Cell Core and Neurobiology Imaging Facility for technical assistance; Sinisa Hrvatin, Aurel Nagy, Rory Kirshner, and Michael Greenberg for analytic guidance and helpful discussions. W.R. is supported by NINDS K08NS101064 and the Migraine Research Foundation. C.J.W. is supported by NINDS R35NS105076 and the Bertarelli Foundation, Dr. Miriam and Sheldon G. Adelson Medical Research Foundation, and the DARPA Panacea program (HR0011-19-2-0022)

## Author Contributions

W.R. and I.T. designed, performed, and analyzed data for most experiments in this study. L.Y. performed data analysis and designed the website. Y.C. performed and analyzed experiments related to *Atf3* and generated the *Atf3-Cre^ERT2^* mice. E.L. assisted with experiments. R.K. and D.G. contributed to *Atf3KO* gene profiling. W.R., I.T., L.Y., and C.J.W. wrote the manuscript. W.R., I.T. and C.J.W. supervised all aspects of the study.

## Methods

### Animals

Male and female 8-12-week-old C57 mice were obtained from Jackson Labs (strain #000667) and used in most behavioral and snRNA-seq experiments. Unless stated otherwise, male mice were used in all experiments. *The Atf3-Cre^ERT2^* mice were generated by inserting an IRES_Cre^ERT2^_pA cassette at the 3’UTR of the mouse *Atf3* locus in order to preserve endogenous *Atf3* expression. CRISPR guide RNAs were designed to produce a defined double-strand break (DSB) at the 3’UTR in order to enable homology-directed repair (HDR). The HDR donor sequence consisted of IRES_Cre^ERT2^_pA cassette flanked by two homologous arms 1 kb (left-arm) and 4 kb (right-arm) in length. We mixed synthetic sgRNA targeting at 3’UTR of mouse *Atf3*, Cas9 protein and HDR donor, and then injected the mixture directly into single-cell mouse embryos. *Atf3-Cre^ERT2^*;*Gcamp6f f/f* mice were generated by crossing the *Atf3-Cre^ERT2^* transgenic mice with *Gcamp6f f/f* mice from Jackson Labs (strain #024105) and bred to homozygosity for both alleles. *Gcamp6f* reporter expression was induced in injured *Atf3-Cre^ERT2^*;*Gcamp6f f/f* mice 24 hrs after injury by intraperitoneal (i.p.) tamoxifen injection at the same time as in naive *Atf3-Cre^ERT2^*;*Gcamp6f f/f* mice. *Atf3f/f* mice were generated by inserting loxP sites around exon 3 of the mouse *Atf3* gene. *Vglut2-Cre;Atf3f/f* and *Brn3a-Cre^ERT2^*;*Atf3f/f* mice were generated by crossing the *Atf3f/f* mice with *Vglut2-ires-Cre* (strain #016963) or *Brn3a-Cre^ERT2^* (strain #032594) mice from Jackson Labs. These mice were bred as homozygotes for *Atf3f/f* and heterozygotes for *Vglut2-Cre* or *Brn3a-Cre^ERT2^*. Littermate controls were used for experiments involving transgenic mice. Injured *Vglut2-Cre;Atf3f/f* cKO DRG neurons express *Atf3* mRNA as measured by FISH (data not shown) and snRNA-seq (Table S8), but do not express nuclear ATF3 protein in sensory neurons (Figure 7B). *Mrgprd-Cre^ERT2^;Gcamp6f* mice were generated by crossing the *Mrgprd-Cre^ERT2^* transgenic mice from Jackson Labs (strain #031286) with *Gcamp6f f/f* mice (strain #024105) and bred to homozygosity for both alleles. All animal experiments were conducted according to institutional animal care and safety guidelines at Boston Children’s Hospital and Harvard Medical School.

### Surgical Procedures

Sciatic nerve crush and ScNT were performed as previously described (Ma et al., 2011), and the SpNT protocol was modified from previous reports (Ogawa et al., 2014; Vilceanu et al., 2010). Briefly, mice were anesthetized by administration of 2.5% isoflurane. Sciatic nerve crush and ScNT were performed by exposing the left sciatic nerve at the mid-thigh level and crushing with smooth forceps for 30 s or cutting a 2mm segment with a pair of scissors followed by a tight ligation of the proximal end to prevent regeneration, respectively. SpNT was performed by making a midline incision of mouse back skin, exposing the left L3 and L4 spinal nerves on the visual field and cutting them with a pair of scissors. These two ganglia were selected in order to maximize the number of transected sensory axons in the sciatic nerve. Intraperitoneal injections of 4mg/kg paclitaxel every other day for 6 days (total of 4 injections) were performed as previously described (Toma et al., 2017). A single intraplantar injection of 20µl CFA was performed into the left hindpaw as previously described (Ghasemlou et al., 2015). Naive and treated mice were euthanized by CO_2_ asphyxiation and decapitation. Ipsilateral lumbar L3-L5 ganglia from naive, crush, ScNT, paclitaxel or CFA-treated mice and ipsilateral L3-L4 ganglia from SpNT mice were collected at various time points after treatment. Ganglia were from 3-5 mice per sample were immediately frozen on dry ice, then pooled for subsequent snRNA-seq profiling or histology. There were 2-7 biological replicates of each pooled condition, as indicated in Figure S1. Two biological replicates were used in snRNA-seq experiments of *Atf3* cKO mice. Each replicate of a specific condition (naive or crush) or genotype (*Atf3* cKO; *Vglut2-Cre;Atf3f/f* or littermate WT controls; *Atf3f/f*) contained L3-L5 DRGs pooled from 1 male mouse and 1 female mouse.

### Single-nuclei isolation from mouse DRG

Single-nuclei suspensions of lumbar DRGs from naive or injured/treated mice were collected using a modified protocol from that described previously (Renthal et al., 2018). Briefly, DRGs were removed from dry ice and placed into homogenization buffer (0.25 M sucrose, 25 mM KCl, 5 mM MgCl_2_, 20 mM tricine-KOH, pH 7.8, 1 mM DTT, 5 μg/mL actinomycin, and 0.04% BSA). After a brief incubation on ice, the samples were briefly homogenized using a tissue tearer and transferred to a Dounce homogenizer for an additional ten strokes with a tight pestle in a total volume of 5mL homogenization buffer. After ten strokes with a tight pestle, a 5% IGEPAL (Sigma) solution was added to a final concentration of 0.32% and five additional strokes with the tight pestle were formed. The tissue homogenate was then passed through a 40-μm filter, and diluted 1:1 with OptiPrep (Sigma) and layered onto an OptiPrep gradient as described previously (Mo et al., 2015). After ultracentrifugation, nuclei were collected between the 30 and 40% OptiPrep layers. This layer contains DRG nuclei as well as some membrane fragments likely from Schwann cells that have the same density as nuclei. We diluted this layer in 30% OptiPrep to a final concentration of 80-90,000 nuclei+fragments/mL for loading into the inDrops microfluidic device. All buffers and gradient solutions for nuclei extraction contained RNAsin (Promega) and 0.04% BSA.

### Single-nucleus RNA sequencing (inDrops)

Single-nuclei suspensions were encapsulated into droplets and the RNA in each droplet was reverse transcribed using a unique oligonucleotide barcode for each nucleus as described previously (Klein et al., 2015). Nuclei encapsulation was performed in a blinded fashion and the order of sample processing was randomized. After encapsulation, the sample was divided into pools of approximately 3,000 droplets and library preparation was performed as described previously (Hrvatin et al. 2017). Libraries were sequenced on an Illumina Nextseq 500 to a depth of 500 million reads per 30,000 droplets collected, resulting in at least 5 reads per UMI on average per sample. Sequencing data was processed and mapped to the mouse genome GRCm38 (modified by the addition of 3’ regions of *Gcamp6f*-WPRE and *Cre*) using the pipeline described in https://github.com/indrops/indrops (Klein et al., 2015). Counts tables from each library were then combined and processed as described below.

### Initial quality control, clustering and visualization of snRNA-seq

To be included for analysis, nuclei were required to contain counts for greater than 600 unique genes, fewer than 15,000 total UMI, and fewer than 10% of the counts deriving from mitochondrial genes. There were 171,827 nuclei that met these criteria. We used the Seurat package (version 2.3.4) in R to perform clustering of these nuclei as previously described (Satija et al., 2015). Raw counts were scaled to 10,000 transcripts per nucleus to control the sequencing depth between nuclei. Counts were centered and scaled for each gene. The effects of total UMI and percent of mitochondrial genes in each nucleus, as well as the batch in which the library was prepared were regressed out using a linear model in Scaledata() function. Highly variable genes were identified using the MeanVarPlot()with default settings. The top 20 principal components were retrieved with the RunPCA() function using default parameters. Nuclei clustering was performed using FindClusters() based on the top 20 principal components, with resolution at 1.5 for the initial clustering of all nuclei and the sub-clustering of non-neuronal nuclei except where otherwise specified. For dimension reduction and visualization, Uniform Manifold Approximation and Projection (UMAP) coordinates were calculated in the PCA space by using the implemented function runUMAP() in Seurat.

### Doublet identification and classification of neuronal and non-neuronal nuclei

After clustering all DRG nuclei that passed initial quality control metrics as above, we next excluded nuclei from downstream analysis that were likely to be doublets. Specifically, nuclei that expressed marker genes (> 0.5 standard deviations away from the mean of the nuclei included for clustering) from multiple cell types were classified as doublets and excluded from downstream analysis. After doublet removal, 145,338 nuclei were included for downstream analysis (97,137 neuronal nuclei and 48,201 non-neuronal nuclei). The marker genes used to make doublet calls were neurons = *Rbfox3*, endothelial = *Cldn5*, macrophages = *Mrc1*, glia = *Mbp*, and meninges = *Mgp*). A nucleus was also classified as a doublet if it expressed multiple neuronal subtype marker genes (peptidergic nociceptors (PEP) = *Tac1*, non-peptidergic nociceptors (NP) = *Cd55*, pruriceptors (SST) = *Sst*, cLTMR = *Fam19a4*, A-LTMR (NF) = *Nefh*. Clusters enriched for the expression of the neuronal marker gene *Rbfox3* were classified as neuronal clusters, and clusters enriched for the expression of the non-neuronal marker genes *Cldn5, Mrc1, Mbp,* or *Mgp* were classified as non-neuronal clusters.

### Annotation of non-neuronal DRG cell types

Non-neuronal subtypes (defined by low *Rbfox3* expression and expression of any non-neuronal marker) were clustered separately as described above to facilitate classification of non-neuronal subtypes. Doublet removal was performed again with higher stringency to remove nuclei from downstream analysis that expressed marker genes from multiple cell types (marker gene expression > 1 standard deviation away from the mean of non-neuronal nuclei). The same genes were used as above to make doublet calls. Significant enrichment (FDR< 0.01, log_2_FC > 0.5) of known non-neuronal marker genes within a cluster of nuclei compared to all other nuclei was used to assign the respective non-neuronal cell type to each cluster (satellite glia = *Apoe*, Schwann cells = *Mpz*, meninges = *Mgp*, endothelial cells = *Cldn5*, and pericytes/endothelial = *Flt1*). The final non-neuronal dataset after quality control contains 34,108 nuclei, with 33 clusters corresponding to 6 cell types.

### Annotation of neuronal DRG subtypes

Neuronal nuclei (classified as above) were clustered separately as described above to facilitate neuronal subtype classification. Doublet removal was performed again with higher stringency to remove nuclei from downstream analysis that expressed marker genes from multiple neuronal subtypes (marker gene expression > 1 standard deviation away from the mean of the neuronal nuclei). The same neuronal subtype marker genes were used as above to make doublet calls. Significant enrichment (FDR< 0.01, log_2_FC > 0.5) of known neuronal subtype marker genes within a cluster of nuclei compared to all other neuronal nuclei was used to assign the neuronal subtype to each cluster. Specifically, peptidergic nociceptors (PEP)1 = *Tac1,Gpx3*; PEP2 = *Tac1,Hpca*; non-peptidergic nociceptors (NP) = *Cd55*; non-peptidergic/itch receptors (SST) = *Sst*; cLTMR1 = *Fam19a4,Th+*; p_cLTMR2 = *Fam19a4*, *Th-*; A-LTMR (NF1) = *Nefh, Cadps2*; proprioceptors (NF2) = *Nefh, Pvalb*; A-LTMR (NF3) = *Nefh, Cplx2*. Each of these subtypes was confirmed by FISH (see Figure S1). We removed 4 neuronal clusters that were significantly enriched for *Rgs11* after being unable to confirm this cell population by FISH. The final neuronal dataset after quality control contains 73,433 high-quality nuclei, with 37 clusters corresponding to 9 neuronal subtypes and “injured state” neurons (see below).

### Classification of naive and injured transcriptional states

To quantitatively classify neurons as being in either a transcriptionally “naive state“ or “injured state,” we calculated the percent of nuclei that were derived from naive mice or SpNT mice within each neuronal cluster. Percentages were calculated with all 7,742 naive neuronal nuclei and 6,482 SpNT neuronal nuclei > 1 day after injury. Clusters of neuronal nuclei were classified as in the “injured state” if >95% of the nuclei in that cluster were derived from SpNT mice and median log_2_-normalized expression of injury induced genes *Atf3* greater than 2. All other clusters were classified as “naive,” which on average had ∼7% (roughly the percent of un-axotomized neurons after SpNT) of their nuclei from SpNT mice and a median *Atf3* expression of 0.

### Classification of injured neuronal subtypes

The “injured state” neurons lose most of the distinguishing gene expression features used for classifying neuronal subtypes (e.g. *Tac1* expression for PEP). Thus, to classify “injured state” neuronal subtypes, we aimed to identify more subtle gene expression signatures that could be used to distinguish between neuronal subtypes after injury. To do this, we co-clustered nuclei from two consecutive time points after SpNT, reasoning that if we had sufficient temporal resolution of the transition states between “naive” and “injured” neurons, we could project remaining neuronal subtype-specific transcriptional signatures from one time point to the next even after the primary marker genes are downregulated. Each pairwise co-clustering was pairwise as follows: naive and 6h after SpNT, 6h and 12h 6h after SpNT, 12h and 1d 6h after SpNT, 1d and 1.5d 6h after SpNT, 1.5d and 2d 6h after SpNT, 2d and 3d 6h after SpNT, and 3d and 7d 6h after SpNT. The neuronal subtype classifications of naive neuronal clusters were then projected onto “injured”/unknown neuronal nuclei from 6h after SpNT that were present in the same cluster. We then used the new neuronal subtype classifications of 6h SpNT nuclei to guide the classification of “injured”/unknown nuclei 12h after SpNT, and continued in this fashion until nuclei from all SpNT time points were classified.

For each pairwise clustering and projection step, if > 50% of the total nuclei (classified + unknown) in a cluster were already assigned to a specific neuronal subtype, either from the initial clustering above using marker gene expression or projection from an earlier pairwise clustering step, this subtype classification was projected to all nuclei in the same cluster. If ≤ 50% of the total nuclei in a cluster had a known subtype classification, we determined whether the classified nuclei in these clusters were all from the same subtype or multiple subtypes. If they were from the same subtype, we next used the FindMarkers() function in Seurat to identify cluster-specific gene expression patterns as described previously. If known subtype-specific marker genes were significantly enriched in a specific cluster (FDR<0.01, log_2_FC > 0.5), we assigned this cluster the corresponding subtype as described above (e.g. *Tac1*+ clusters are peptidergic nociceptors). If multiple previously-classified neuronal subtypes were present in a cluster, we re-clustered these nuclei separately to maximize the potential to separate neuronal subtypes into biologically meaningful clusters. After re-clustering, the FindMarkers() function in Seurat was performed on each cluster as described previously to identify cluster-specific gene expression patterns. If known subtype-specific marker genes were significantly enriched in a specific cluster (FDR<0.01, log_2_FC > 0.5), we assigned this cluster the corresponding subtype as described above. If known marker genes were not enriched in a cluster even after re-clustering, we classified these clusters as unknown (1.9% of SpNT nuclei).

To assign the neuronal subtypes of “injured state” nuclei from crush, ScNT, paclitaxel, CFA, naive, and the “unknown” SpNT nuclei above, we clustered all “injured state” neuronal nuclei in the study together. Having classified most SpNT nuclei previously, we were able to project those neuronal subtypes onto the “injured state” nuclei from other models. We assigned clusters to the neuronal subtype of the most abundant SpNT neuronal subtype in that cluster if it was more than 3X more abundant than the next most abundant subtype in that cluster (88.5% of nuclei classified this way). Otherwise, nuclei from the remaining clusters were separately clustered and each new cluster was assigned to a neuronal subtype depending on the number proportion of previously classified neurons in that cluster. A neuronal subtype was then assigned to the new cluster if > 80% of previously-classified SpNT nuclei in the new cluster were of the same neuronal subtype (on average ∼1/3 of the nuclei within a cluster were previously-classified SpNT nuclei and ∼2/3 were of unknown subtype) (7.5% of nuclei classified this way). If ≤ 80% of the previously-classified SpNT nuclei in the new cluster were of the same neuronal subtype, we assigned the new cluster to the most abundant subtype in that cluster (4% of nuclei classified this way).

We also used an independent bioinformatic approach in which injury-induced gene expression within each cell is regressed out prior to subtype. To do this, we used the FindMarkers() function in Seurat to identify differential gene expression (FDR<0.01 and log_2_FC > 1) between “injured state” clusters and “naive state” clusters across all injury models. Seventy-five genes were identified, and a score was generated with these genes using the AddModuleScore() function in Seurat. This function calculates the mean normalized expression of the specified gene set, subtracted by the mean normalized expression of a random gene set for each single nucleus. We then scaled the counts matrix using the Scaledata() function in Seurat, including the injury score along with UMI, % mitochondrial genes, and batch to the linear regression. The regressed counts matrix was then clustered with default settings described above. Regressing out the injury score resulted in “injured state” nuclei clustering with their “naive state” counterparts, which enabled cell types to be assigned to each cluster based on their marker gene expression as described above. Neuronal subtypes assigned by the regression method were compared to the neuronal subtypes assigned by pairwise clustering and projection, and the concordance rate was 99% for naive nuclei and 91% for injured nuclei.

### Lineage tracing of injured non-peptidergic neurons

Neuronal DRG nuclei from tamoxifen-treated *Mrgprd-Cre^ERT2^* mice (naive and 7d after crush) were co-clustered with neuronal nuclei from our injury time course with default clustering settings in Seurat. Neuronal subtypes were identified by pairwise clustering and projection described above. We then calculated the fraction of nuclei in each neuronal subtype that expresses the *Gcamp6f* reporter of *Mrgprd+* NP neurons greater than the threshold. The threshold was set as the median *Gcamp6f* expression of all *Gcamp6f-* expressing nuclei from *Mrgprd-Cre^ERT2^* mice. The error rate (1.88 for “naive state” nuclei, 2.93% for “injured state” nuclei), for neuronal classification by pairwise clustering and projection was reported as the fraction of non-NP neuronal nuclei expressing *Gcamp6f* greater than the threshold.

### Differential expression analysis

Differential expression analysis was done with edgeR (version 3.24.3) similar to that described for single-cell analysis in (Soneson and Robinson, 2018). Briefly, edgeR uses the raw counts as input, and genes detected in fewer than 5% of nuclei selected for each comparison were excluded from analysis. Counts within each nucleus were normalized by the trimmed mean of M-values (TMM) method to adjust for total RNA differences between nuclei. Dispersion was estimated by fitting a quasi-likelihood negative binomial generalized log-linear model (glmQLFit) with the conditions being analyzed. The QL F-test was used to determine statistical significance between differentially expressed genes in the experimental and control groups. For each experimental condition (e.g. NP neurons 6h after SpNT), the control group used for each comparison was the corresponding cell type from naive animals, unless otherwise specified. Differentially regulated genes are defined as genes with FDR<0.01 and log_2_FC > |1|.

### Cell-type-specificity score

“Cell-type-specific” genes in naive animals were identified using the FindMarkers() function in Seurat to compare gene expression in nuclei of each cell type to all other nuclei (FDR<0.01 and log_2_FC > 1). These “cell-type-specific” genes for each cell type were used to generate cell-type-specificity scores using the AddModuleScore() function in Seurat, which resulted in a distinct cell-type-specificity score for each cell type. Each nucleus was assigned to the cell-type-specificity score of its respective cell type.

### Common injury score

The 438 injury-induced genes that are present in ≥ 5 neuronal subtypes (see common genes in Figure 4D, Table S3) are used to generate the common injury score. The injury score was calculated for each nucleus by using the AddModuleScore() function in Seurat as described above.

### Random gene selection

To generate expression-matched control gene lists, genes in each cell type were first ranked by their level of expression, and then for each cell-type-specific gene, the gene either above or below it was selected randomly. Random gene lists for motif enrichment analysis were generated as described in that section.

### Gene ontology (GO) analysis

GO analysis was performed using topGO (version 2.34.0) in R. Expressed genes (≥ 5% of SpNT+naive nuclei with the mean log_2_-normalized expression > 0.1 from edgeR analysis in any neuronal subtype) were used as the background list. The common injury-induced genes described above were used as the input gene list. R package org.Mm.eg.db (version 3.7.0) was used as the genome wide annotation database for *Mus musculus*. Genes were annotated for their biological process and associated gene ontology terms with > 10 annotated genes and enrichment *P*-value < 0.05 were returned. Enrichment is defined as the number of annotated genes observed in the input list divided by the number of annotated genes expected from the background list.

PANTHER was used to categorize the molecular function of cell-type-specific genes (Figure S2A) using default settings for *Mus musculus*. Genes containing the molecular function of transcription factors, ion channels, and GPCRs were selected and used for plotting. Neuropeptide gene lists were obtained from http://www.neuropeptides.nl/tabel%20neuropeptides%20linked.htm.

### Transcription factor analysis

We used SCENIC package (version 1.1.1-9) (Aibar et al., 2017) to conduct gene regulatory network analysis and transcription factor assessment. For inclusion in this analysis, genes needed to be detected in at least 5% of nuclei and have a mean log_2_-normalized expression > 0.1. To identify potential transcription factor targets, SCENIC first performs a co-expression network analysis to identify the genes whose expression is positively correlated (Pearson’s r > 0.01) with each transcription factor expressed in the dataset. For each transcription factor and its corresponding module of genes that are positively correlated with it, SCENIC uses RcisTarget to perform motif enrichment analysis to identify the putative regulon for each transcription factor. RcisTarget was run with default settings and motif enrichment was calculated based on regions 500 bp upstream and 20 kb centered (10 kb upstream + 10kb downstream) around the transcription start site of each gene. Once a regulon is assigned for each transcription factor, SCENIC then calculates a score (AUCell) that represents the “activity” of each transcription factor within each cell based on the expression of the transcription factor and its target genes. Only transcription factors that were identified by SCENIC and also present in the list of annotated mouse transcription factors from AnimalTFDB database (http://bioinfo.life.hust.edu.cn/AnimalTFDB/) were included in the study.

### Gene set motif enrichment analysis

To identify motifs that are significantly enriched in a gene set, motif enrichment analysis was run with RcisTarget (version 1.3.5). Motif analysis was performed for 20 kb regions centered (10 kb upstream + 10kb downstream) around the transcription start site of each gene. RcisTarget assigns an enrichment score for each motif based on its frequency near the transcription start site of our input gene list compared to its average frequency in the genome. Enrichment scores for each motif were then normalized and motifs with normalized enrichment scores > 3SD are considered enriched. The relative activity of the injury-induced transcription factors (see Figure 6B) was predicted by counting the motifs they are known to bind within the set of enriched motifs within a given input gene list (e.g. 438 common injury-induced genes). Motif enrichment was performed on the set of common injury-induced genes (see Table S3) and cell-type-specific genes (see Table S6) as well as random gene sets. To calculate motif enrichment for random gene sets, motif analysis was averaged across 1000 sets of either 438 randomly selected expressed genes (to compare with common injury-induced genes) or 1240 randomly selected expressed genes (to compare with cell-type-specific genes).

### Bulk RNA-seq library preparation and sequencing

Total RNA was extracted from DRG tissue samples using TRIzol (ThermoFisher), and then purified using total RNA mini kit (Qiagen). Quality control assessment of these purified RNA samples was conducted using Bioanalyzer (Agilent) and the RNA integrity numbers (RIN) of all RNA samples submitted for sequencing were > 7. RNA-sequencing was carried out using the NuGEN Ovation RNA Ultra Low Input kit and TruSeq Nano. Libraries were indexed and sequenced by HiSeq2500/HiSeq4000 with 69-bp paired end reads. Quality control (QC) was performed on base qualities and nucleotide composition of sequences, to identify problems in library preparation or sequencing. Reads were trimmed if necessary after the QC before input to the alignment stage. Reads were aligned to the Mouse mm10 reference genome (GRCm38.75) using the STAR spliced read aligner (ver 2.4.0). Average input read counts were 63.7M per sample and average percentage of uniquely aligned reads were 76.5%. Total counts of read-fragments aligned to known gene regions within the mouse (mm10) refSeq (refFlat ver 07.24.14) reference annotation are used as the basis for quantification of gene expression. Fragment counts were derived using HTSeq program (ver 0.6.0). Batch effect was removed using Bioconductor package ComBat and RUV (removal of unwanted variation). Differentially expressed genes were identified using the Bioconductor package edgeR (FDR ≤ 0.1). Scripts used in the RNA sequencing analyses are available at https://github.com/icnn/RNAseq-PIPELINE.git.

### Behavioral Experiments

Mouse behavior experiments were performed as previously described (Ghasemlou et al., 2015; Latremoliere et al., 2018; Sakuma et al., 2016). Briefly, von Frey filaments were used to measure the mechanical sensitivity of ipsilateral mouse hindpaws by blinded experimenters. 50% von Frey thresholds were calculated using the Up-Down Reader (Gonzalez-Cano et al., 2018). Responses to pinprick stimulation of different parts of the ipsilateral hindpaw were recorded in the same animals by blinded experimenters at different time points following sciatic nerve crush as previously described (Sakuma et al., 2016).

### RNAScope *in situ* histochemistry

RNAscope fluorescence *in situ* hybridization (FISH) experiments were performed according to the manufacturer’s instructions, using the RNAscope Multiplex Fluorescent kit (Advanced Cell Diagnostics (ACD)) for fresh frozen tissue, as previously described (Zeisel et al., 2018). Briefly, fresh frozen ipsilateral naive or injured L4 lumbar DRGs were dissected at various points after injury, fresh frozen and sectioned into 12 µm sections using a cryostat. *In situ* probes were ordered from ACD and multiplexed in the same permutations across quantified sections. Following FISH, some sections were imaged using a 20x widefield objective on an Olympus Slide Scanner microscope. In order to quantify marker gene expression, high resolution images of a single z-plane were obtained using a 60x oil immersion objective on a Perkin Elmer UltraView Spinning Disk confocal microscope.

### Fluorescence *in situ hybridization* quantification

L4 DRG section images from 3-6 mice per probe were used for quantification. All in-focus neurons were manually segmented by blinded scorers using *Tubb3* fluorescence. Images were then thresholded, puncta were automatically quantified using ImageJ and puncta counts per µm^2^ per neuron compared across conditions. For sciatic crush sections (Fig S5N), cutoffs were set to the mean of *Atf3* puncta density in naive neurons plus 2 standard deviations, and neurons after crush are divided into *Atf3* high (injured, *Atf3* puncta density > cutoff) and *Atf3* low (uninjured, *Atf3* puncta density ≤ cutoff) populations; for SpNT slides (Fig 2H), neurons were analyzed as one population. Then neurons with the most marker puncta density in each condition were selected for visualization and statistical tests in accordance with the relative abundance of naive neuronal cell types in the snRNA-seq data (top 9.28% of neurons were selected for marker *Th* (cLTMR), top 18.33% for *Tac1* (PEP), top 22.43% for *Mrgprd* (NP), top 21.51% for *Hapln4* (NF), and top 4.25% for *Sst* (SST). One-way analysis of variance (ANOVA) was carried out by calling anova() function in R to compare means in different conditions. As the ANOVA test is significant, Tukey Multiple Comparisons are conducted to compare between conditions by calling TukeyHSD() function in R.

### Western Blot

*Brn3a-Cre^ERT2^;Atf3f/f* mice were injected intraperitoneally with tamoxifen or vehicle. Two weeks after induction, the mice underwent sciatic nerve crush. Ipsilateral L3-5 DRGs were harvested from 4 mice (12 DRGs/mouse) 1 week after crush and pooled for protein extraction. The protein lysates were extracted in presence of a protease cocktail tablet (Roche Diagnostics) using Cell Lysis buffer (ThermoFisher). Cell debris was removed by centrifugation (4°C, 10 min) after homogenization. Protein concentrations were determined using the BCA protein assay kit (ThermoFisher). Equivalent amounts of protein were loaded and separated by 4-12% gradient SDS-PAGE and subsequently transferred to an Immobilon-P PVDF transfer membrane (EMD Millipore). Blots were blocked in 5% blotting-grade blocker (Bio-rad) in PBS for 20 min at room temperature (RT) and incubated with rabbit polyclonal antibodies against ATF3 (Santa Cruz, 1:500, RRID: AB_1078233), and Horseradish peroxidase (HRP)-conjugated mouse monoclonal antibody against GAPDH (Cell Signaling, 1:5000, RRID:AB_1642205) overnight. After washing 3 times with TBST (1% Tween-20), HRP-conjugated secondary antibody (anti-rabbit, ThermoFisher, 1: 20,000), a SuperSignal West Femto Maximum Sensitivity chemiluminescence ECL kit (ThermoFisher), and Amersham Hyperfilm ECL (GE Healthcare Life Sciences) were used for signal detection. Image signals were analyzed and quantified using ImageJ software (NIH).

### Immunohistochemistry

*Vglut2-Cre;Atf3f/f* and *Atf3f/f* mice underwent SpNT. Ipsilateral L4 DRGs were harvested 1 week after SpNT from injured mice, immediately fixed with 4% PFA for 1 hr at 25°C and cryoprotected with 30% sucrose in PBS overnight. DRGs were sectioned into 12μm sections, which were blocked and permeabilized with 5% normal goat serum in 0.25% Triton X-100 in PBS (Roche Diagnostics) for 30 min at 25°C. Sections were incubated with rabbit polyclonal antibody against ATF3 (Sigma Aldrich; HPA001562; 1:1000) at 4°C overnight and then incubated with Alexa Fluor 488 goat antibody against rabbit IgG and Alexa Fluor 488 goat antibody against chicken IgG for 40 min at 25^0^C. Sections were then stained with 1:200 NeuroTrace 640/660 Deep-Red Fluorescent Nissl Stain (Thermo Fisher, N21483, RRID: AB_2572212) for 10 min and mounted with ProLong Gold Antifade Mountant with DAPI (Thermo Fisher, P36931). Slides were imaged using a 20x widefield objective on an Olympus Slide Scanner microscope. Images were thresholded and ATF3+ neurons quantified in ImageJ. Nissl+ DRG neurons were manually counted by blinded scorers. To quantify Nissl+ DRG neuron density, representative 360000 µm^2^ sections of each *Vglut2-Cre;Atf3f/f* and *Atf3f/f* DRG image were selected for quantification.

### Data obtained from other sources

Embryonic DRG development data were obtained from GEO Accessions GSE98592, GSE77892, GSE77891, deposited by the GUDMAP Database Group. We performed differential expression analysis similar to that described above in edgeR to compare the expression profiles of RET+ E12.5, E14.5, E18.5 DRG neurons to adult RET+ DRGs. Briefly, genes with counts <10 were removed from differential expression. Differential expression was otherwise performed using the default settings (calcNormFactors, estimateCommonDisp(y), and estimateTagwiseDisp(y), and exacTest(“adult DRG”, “each embryonic time point”). Regeneration associated gene modules were obtained from (Chandran et al., 2016). Gene names were cleaned up by removing suffix, and genes not detected in our snRNA-seq data were excluded.

### Data Visualization

Plots were generated using R version 3.5.0 with ggplot2 package (version 3.2.0). Heatmaps were generated using gplots package (version 3.0.1.1).

### Statistics

Statistics were performed using R version 3.5.0. Hypergeometric tests were used to test the significance of overlap between two gene sets. It was conducted by calling phyper() function in R version 3.5.0. Permutation tests were used to estimate a *P* value for transcription factor motif enrichment by calculating the number of times out of 1000 the ATF3 motif enrichment was greater in a random set of genes than the experimental set of genes divided by 1000.

## Data availability

Processed data are available at www.painseq.com. Raw and processed data were also deposited within the Gene Expression Omnibus (GEO) repository (www.ncbi.nlm.nih.gov/geo) with an accession number (GSExxxxx). Custom R scripts are available upon request.

## Supplemental Tables

- Table S1: Cell-type-specific gene expression in naive DRG nuclei.
- Table S2: Differential expression analysis between injury/neuropathy models and naive nuclei at each time point after injury.
- Table S3: Genes that are commonly upregulated across ≥ 5 neuronal subtypes after spinal nerve transection compared to their respective naive cell types.
- Table S4: Genes that are upregulated in only 1 neuronal subtype after spinal nerve transection compared to their respective naive cell types.
- Table S5: Common and cell-type-specific gene induction after spinal nerve transection, corresponding to heatmap in Figure 4D.
- Table S6: Genes that are enriched in specific DRG neuronal subtypes in naive mice. These genes are used for cell-type-specificity score.
- Table S7: *Atf3*-dependent gene regulation after sciatic nerve crush
- Table S8: Differential gene expression between embryonic (E12.5, E14.5, E18.5) and adult DRG neurons

---

**Figure S1, related to Figure 1.**
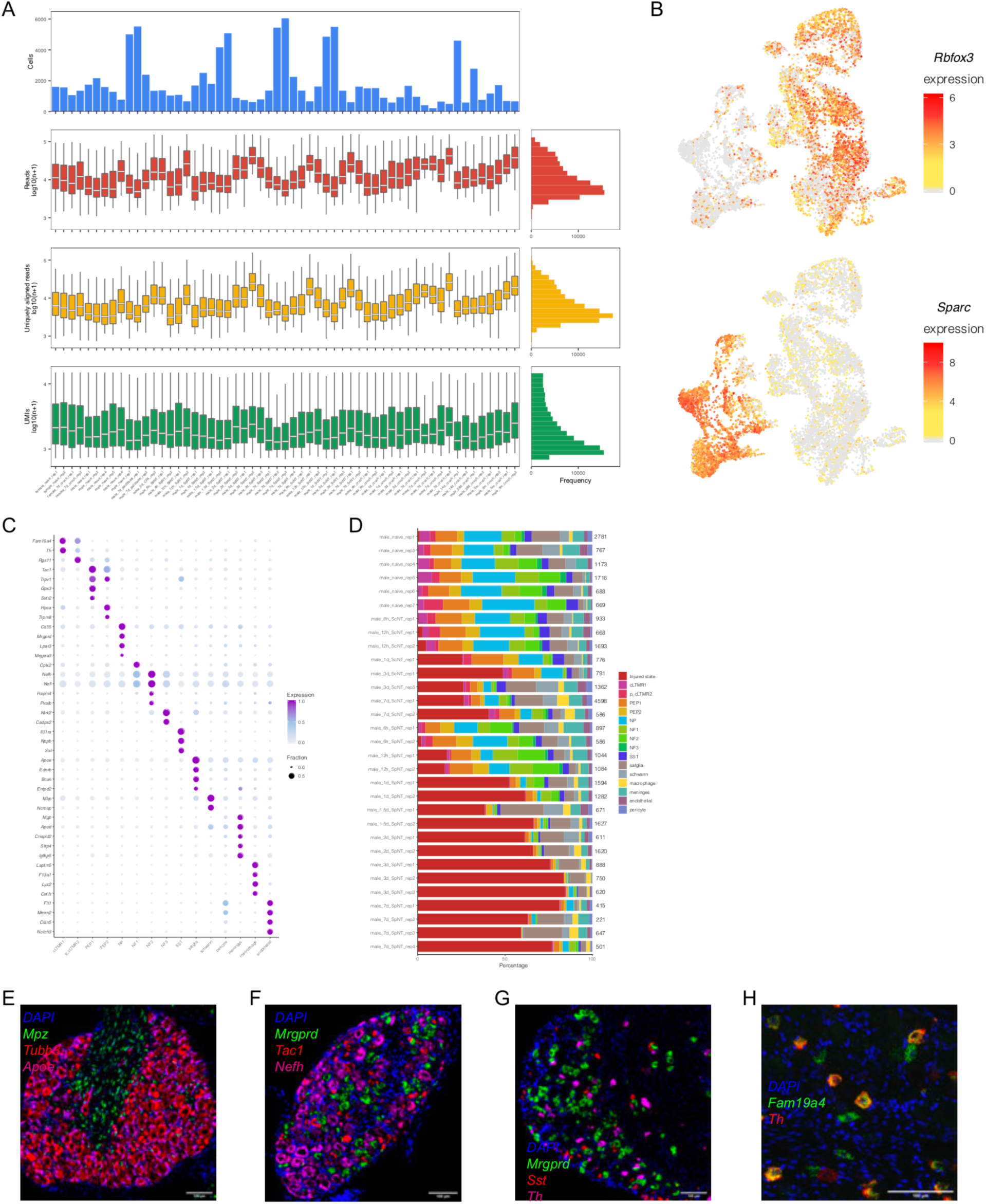
Single-nucleus RNA-seq of mouse DRG before and after injury. **(A)** Sequencing and mapping metrics of 107,541 nuclei that passed quality control and were analyzed in the study. Boxes indicate quartiles and whiskers are 1.5-times the interquartile range (Q1-Q3). Data outside 1.5-times the interquartile range are labeled as dots. The median is a white line inside each box. The distribution is aggregated across all samples and displayed on the horizontal histogram. Number of nuclei collected by sample (top), distribution of reads per sample (log_10_ transformed, second), distribution of uniquely mapped reads per sample (log_10_ transformed, third), distribution of number of unique molecular identifiers (UMI) per sample (log_10_ transformed, bottom). **(B)** UMAP plots of 10,000 randomly sampled nuclei from the 107,541 nuclei passing quality control in the study. Color shows log_2_-normalized expression of the neuronal marker gene *Rbfox3* (top) and non-neuronal marker gene, *Sparc* (bottom). **(C)** Dot plot of cell-type-specific marker genes (rows) in each cell type (columns) of nuclei from naive DRGs. The fraction of nuclei expressing a marker gene is calculated as the number of nuclei in each cell type that express a gene (> 0 counts) divided by the total number of naive nuclei in the respective cell type. Expression in each cell type is calculated as the mean scaled counts of the marker gene relative to the highest mean-scaled counts of that gene across cell types. **(D)** Percentage of nuclei from each biological sample (naive, spinal nerve transection [SpNT], sciatic nerve transection [ScNT]) that were classified into the respective DRG cell types. Neurons that were classified as in the “injured state” are shown in red. The number on the right of each bar shows total number of nuclei that passed quality control for each sample. **(E-G)** Fluorescent *in situ* hybridization (FISH) images of naive L4 mouse DRGs stained with DAPI (blue), *Mpz* (Schwann cell marker, green), *Tubb3* (neuronal marker, red) and *Apoe* (satellite glia marker, magenta) (E); *Mrgprd* (NP [non-peptidergic] DRG neuronal marker, green), *Tac1* (PEP [peptidergic] DRG neuronal marker, red) and *Nefh* (NF [neurofilament+] DRG neuron marker, magenta) (F); *Mrgprd* (NP DRG neuron marker, green), *Sst* (*Sst*+ pruriceptive DRG neuron marker, red) and *Th* (cLTMR DRG neuron marker, magenta) (G). There is minimal overlap between marker gene fluorescence, suggesting these genes are expressed in distinct cell types. **(H)** Representative FISH images of naive L4 mouse DRGs stained with DAPI (blue), *Fam19a4* (cLTMR1 and p_cLTMR2 marker, green) and *Th* (c-LTMR1 marker, red). Some cells express both *Th* and *Fam19a4* at high levels (cLTMR1), while others express *Fam19a4* with little to no *Th* expression (p_cLTMR2). cLTMR = C-fiber low threshold mechanoreceptor; PEP = peptidergic nociceptor; NP = non-peptidergic nociceptor; NF = *Nefh+* A-fiber low threshold mechanoreceptors; SST = *Sst+* pruriceptors.

**Figure S2, related to Figure 2.**
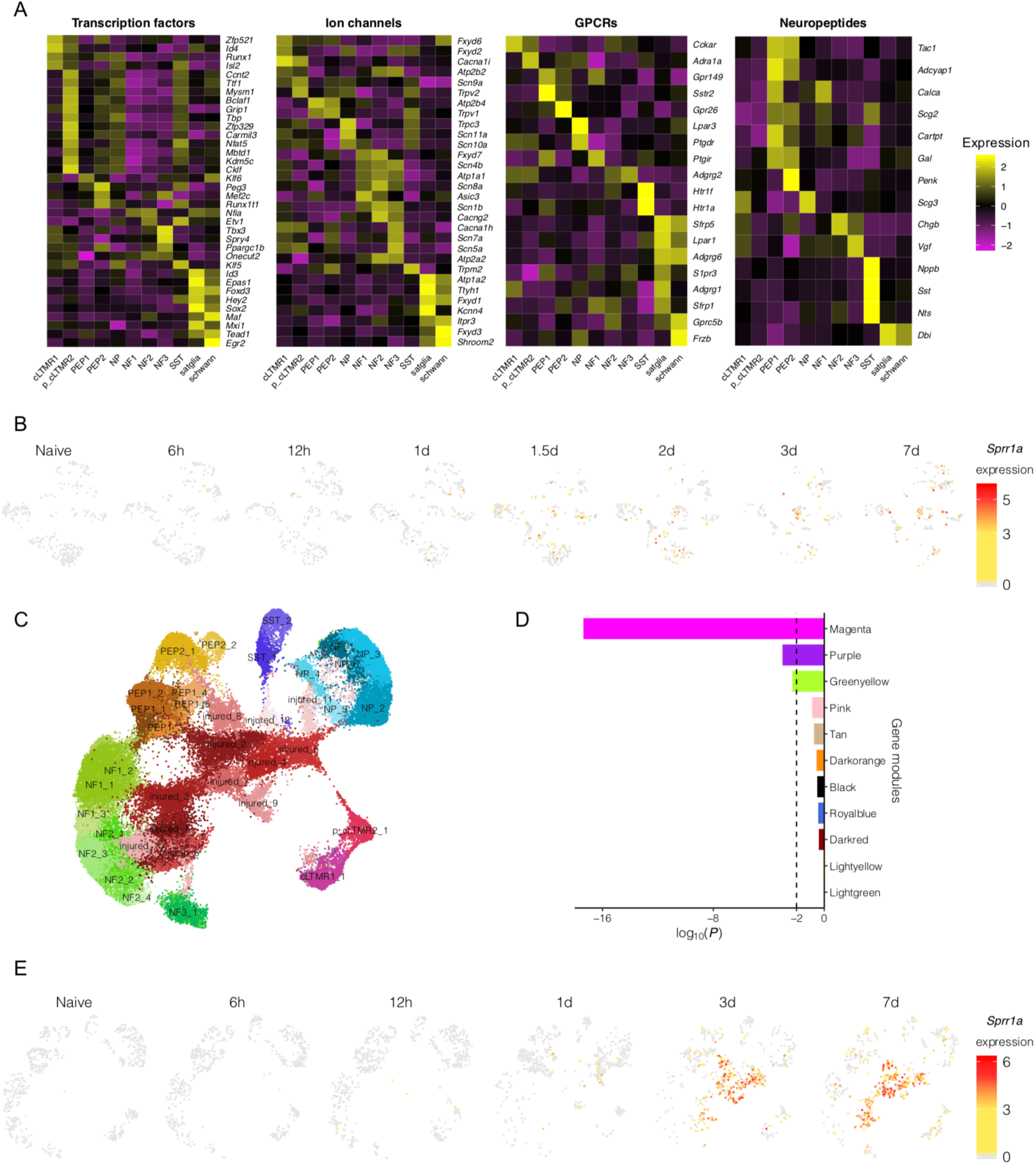
Characterization of DRG neuronal gene expression before and after axonal injury. **(A)** Heatmaps of cell-type-specific gene expression patterns in naive DRG cell types. Genes were included in the heatmap if they demonstrated significant cell type enrichment (FDR < 0.01, log_2_FC > 1) using FindMarkers in Seurat and matched the displayed gene ontology annotations. Heatmaps show scale.data from Seurat, which is the row-normalized and centered mean expression of each gene in a given cell type. **(B)** UMAP plots displaying DRG non-neuronal subtypes at different times after spinal nerve transection. Each time point was randomly sampled to display 300 nuclei. Color denotes log_2_-normalized expression of *Sprr1a,* nuclei not expressing *Sprr1a* are colored grey. **(C)** UMAP plot of all 73,433 neurons that passed quality control from naive mice and mice from each injury model. Cluster IDs that were assigned by Seurat are overlaid onto the plot. Colors denote each cluster ID. **(D)** Comparisons of the overlap between spinal nerve transection (SpNT) injury-induced genes from our single-nucleus RNA-seq data (FDR < 0.01 and log_2_FC > |1|, injured state nuclei after SpNT vs. nuclei from naive animals) and the gene modules identified from microarrays of bulk DRG tissue (Chandran et al., 2016). The magenta module was the predominant injury-induced gene module in the Chandran et al. dataset. Horizontal bars show the log_10_ transformed *P*-values from hypergeometric tests. Vertical dashed line is at *P* = 0.01. **(E)** UMAP plots displaying DRG neurons after sciatic nerve transection. Each time point was randomly sampled to the number of nuclei at the time point with the fewest number of nuclei sequenced (650 neuronal nuclei). Nuclei are colored by their log_2_-normalized expression of the injury-induced gene, *Sprr1a*. cLTMR = C-fiber low threshold mechanoreceptor; PEP = peptidergic nociceptor; NP = non-peptidergic nociceptor; NF = *Nefh+* A-fiber low threshold mechanoreceptors; SST = *Sst+* pruriceptors.

**Figure S3, related to Figure 3.**
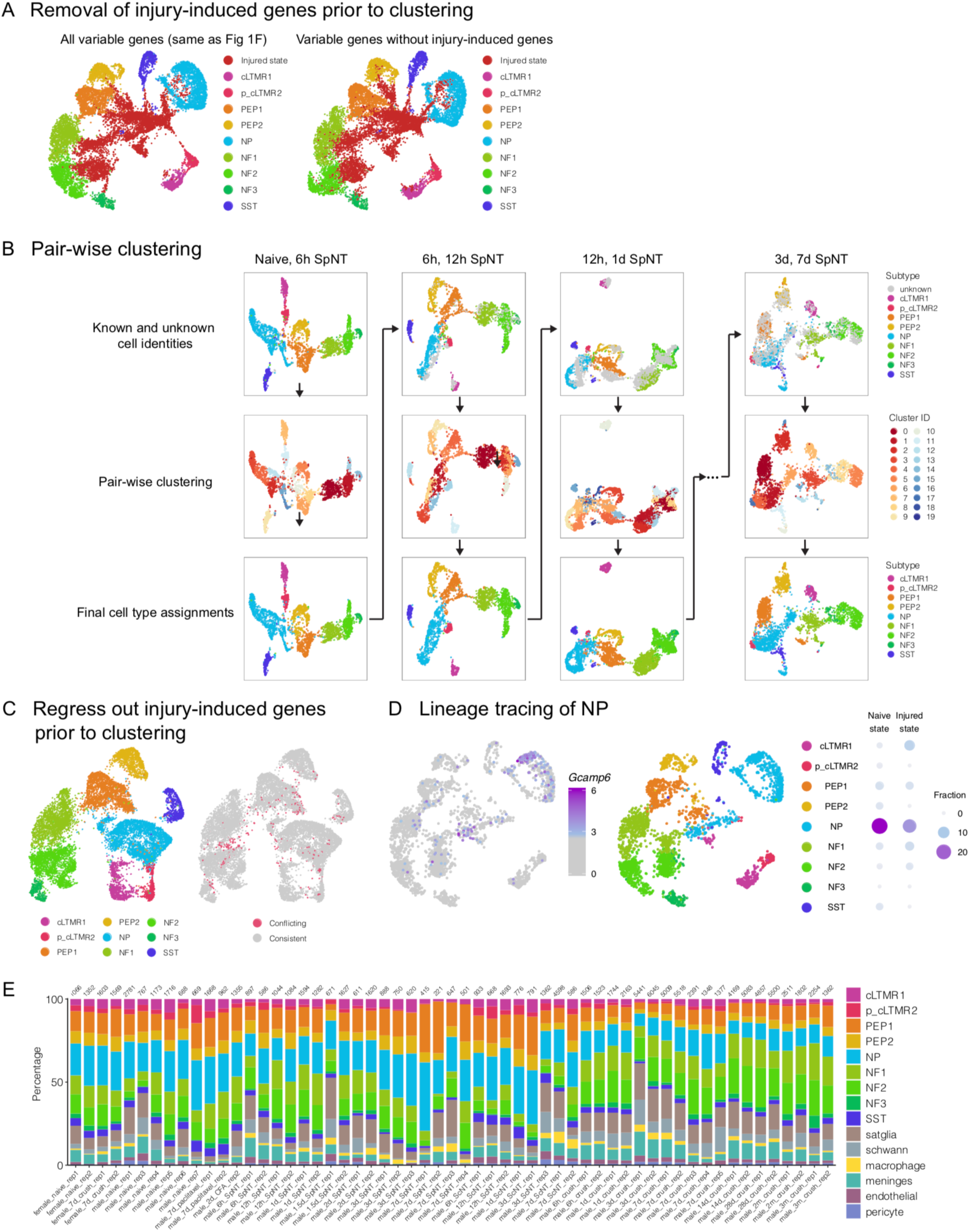
Classification of injured DRG neuronal subtypes after spinal nerve transection (SpNT). **(A)** UMAP plots showing 7,000 naive neuronal nuclei and 7,000 randomly sampled SpNT neuronal nuclei using all variable genes for clustering (left, same as Figure 1F) or after removing injury-induced genes (FDR *<* 0.01, log_2_FC > 0.5, injured state nuclei after SpNT vs. nuclei from naive animals) from the variable genes prior to clustering (right). Colors denote cell types/states. **(B)** Pairwise clustering and projection strategy to classify the neuronal subtypes of injured state nuclei after SpNT. Nuclei of known and unknown neuronal subtypes from each SpNT time point were co-clustered with the subsequent time point collected (top row). Nuclei of unknown neuronal subtype that co-clustered with clusters of marker-gene-confirmed known neuronal subtypes (middle row), were then assigned the respective neuronal subtype of that cluster (bottom row, see methods). The new injured neuronal subtype assignments were projected forward to assist in the subtype assignment of injured neurons at later time points after SpNT (long arrows). Each column shows co-clustering of nuclei from two adjacent time points. Top row colors indicate neuronal subtype with unknown injured nuclei colored gray. Middle row colors indicate cluster IDs assigned by Seurat. Bottom row colors indicate the final subtype assignment after pair-wise clustering and projection. **(C)** UMAP plot showing 7,000 randomly sampled naive neuronal nuclei and 7,000 randomly sampled SpNT neuronal nuclei. Clustering was performed after regressing out the injury-induced genes (FDR *<* 0.01, log_2_FC > 0.5, injured state nuclei after SpNT vs. nuclei from naive animals) from the mRNA counts tables (see methods). Colors denote the independent neuronal subtype assignment using regression-based clustering (left) or the concordance between injured neuronal subtype assignments using the two complementary approaches: pairwise clustering and projection or regression-based clustering (right). **(D)** Lineage tracing to experimentally test neuronal subtype bioinformatic assignments of non-peptidergic nociceptors (NP). UMAP plots of neurons from *Mrgprd-Cre^ERT2^*; *Gcamp6f* reporter mice after tamoxifen treatment. Nuclei are colored by their log_2_-normalized expression of *Gcamp6f* (left, nuclei with *Gcamp6f* expression ≤ median expression of are colored grey), or by their assigned subtypes from pairwise clustering and projection (middle). Fraction of nuclei expressing *Gcamp6f* greater than the median expression are calculated for each naive/injured neuronal subtype (right). Median expression is determined from nuclei with > 0 counts of *Gcamp6f* transcript. **(E)** Fraction of each cell type within individual biological samples sequenced after pairwise clustering and projection was used to classify the neuronal subtypes of nuclei in the “injured state.” The number above of each bar shows total number of nuclei for each sample that passed quality control. cLTMR = C-fiber low threshold mechanoreceptor; PEP = peptidergic nociceptor; NP = non-peptidergic nociceptor; NF = *Nefh+* A-fiber low threshold mechanoreceptors; SST = *Sst+* pruriceptors.

**Figure S4, related to Figure 4.**
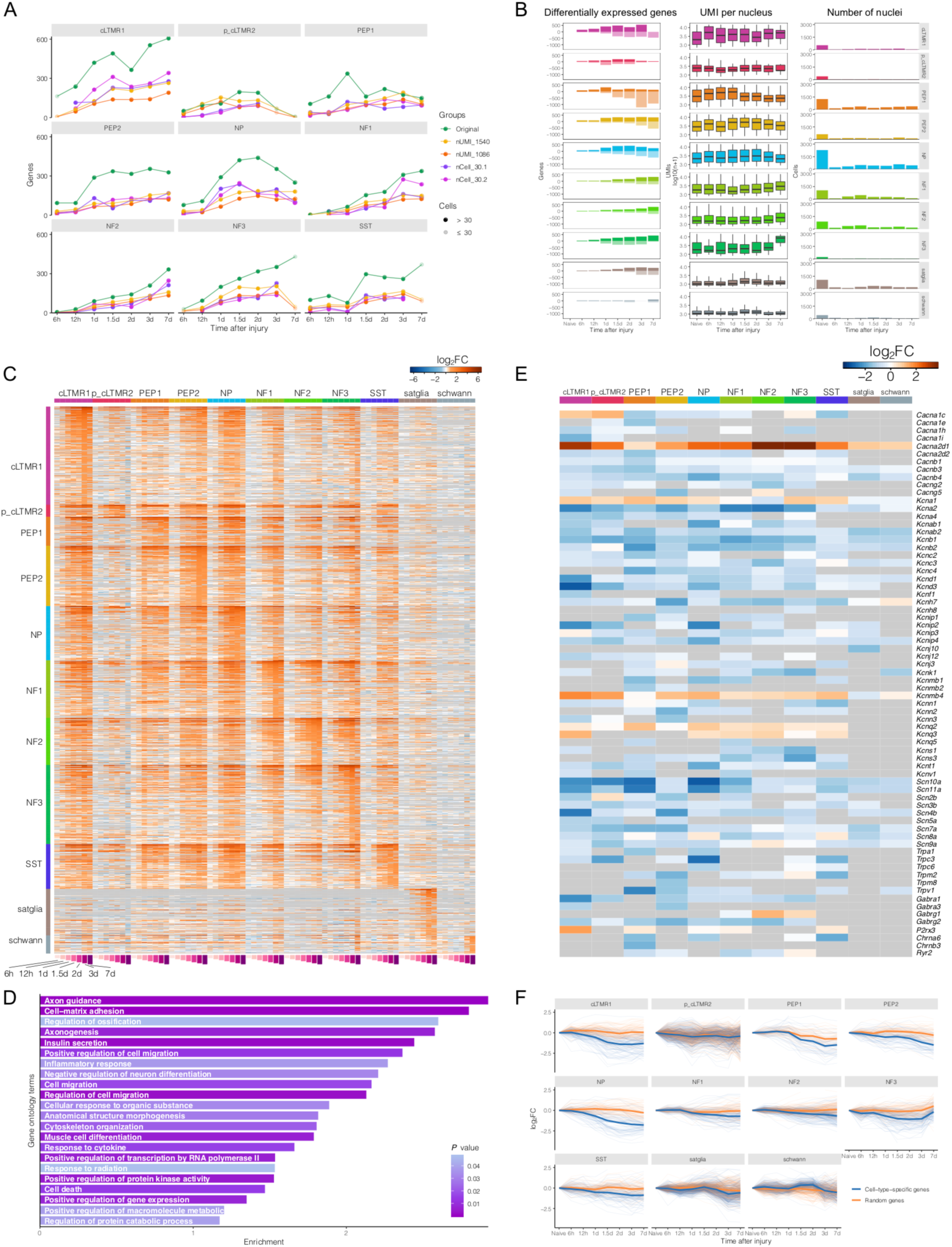
Cell-type specific transcriptional changes in DRG neurons after spinal nerve transection (SpNT). **(A)** Number of significant differentially-expressed genes (FDR < 0.01, log_2_FC > 1) in each neuronal subtype and time point after SpNT compared to naive nuclei of the respective subtype. Lines: original = differential expression including all sequenced nuclei in a given neuronal subtype (green). nUMI_1540 = prior to differential expression, nuclei from all time points and neuronal subtypes are downsampled to an average of 1540 UMI (the lowest average UMI in the SpNT time course, see methods). nUMI_1086 = prior to differential expression, nuclei from all time points and neuronal subtypes are downsampled to an average of 1086 UMI. nCell_30.1 and nCell_30.2 are two independent downsamplings of each neuronal subtype to 30 nuclei prior to differential expression. Solid circles = time points with ≥ 30 nuclei sequenced. Faded circles = time points with < 30 nuclei sequenced. **(B)** Summary of the number of significant differentially-expressed genes (left, positive number indicates significantly upregulated genes with FDR < 0.01 and log_2_FC > 1, and negative number denotes significantly down-regulated genes with FDR < 0.01 and log_2_FC < −1) in each neuronal subtype and time point after SpNT compared to naive nuclei of the respective cell type, UMI per nucleus (middle log_10_ transformed), and total number of nuclei (right) at each time point after SpNT. Boxes indicate quartiles and whiskers are 1.5-times the interquartile range (Q1-Q3). Data outside 1.5-times the interquartile range are omitted for clarity. The median is a black line inside each box. **(C)** Heatmap of log_2_FC of significantly upregulated genes at both 3 and 7 days after SpNT compared to naive nuclei of the respective cell type (FDR < 0.01, log_2_FC > 1). Significantly regulated genes are grouped by cell type, and genes that are significantly regulated in multiple cell types are repeated. Genes that are not expressed in a cell type are colored gray. **(D)** Gene ontology analysis (topGO) of the 438 genes that are commonly induced in >= 5 neuronal subtypes after SpNT compared to naive neurons. The gene ontology terms displayed in the graph are terms that have > 10 annotated significant genes and *P*-value < 0.05. **(E)** Heatmap of the log_2_FC (3d and 7d SpNT nuclei compared to naive nuclei for each cell type) of select genes encoding ion channels. Genes shown on the heatmap are significantly regulated (FDR < 0.01, log_2_FC > |1|) in at least one cell type after SpNT. **(F)** Line plots showing regulation of the cell-type-specific genes within each cell type and time point after SpNT. Cell-type-specific genes are those genes that are expressed significantly higher in one naive cell type compared to all other naive cell types (see methods). For comparison, an equal number of expression-matched randomly-selected genes in each naive cell type are displayed. Bolded lines represent the average log_2_FCs of cell-type-specific genes (blue) or expression-matched random genes (orange). cLTMR = C-fiber low threshold mechanoreceptor; PEP = peptidergic nociceptor; NP = non-peptidergic nociceptor; NF = *Nefh+* A-fiber low threshold mechanoreceptors; SST = *Sst+* pruriceptors.

**Figure S5, related to Figure 5.**
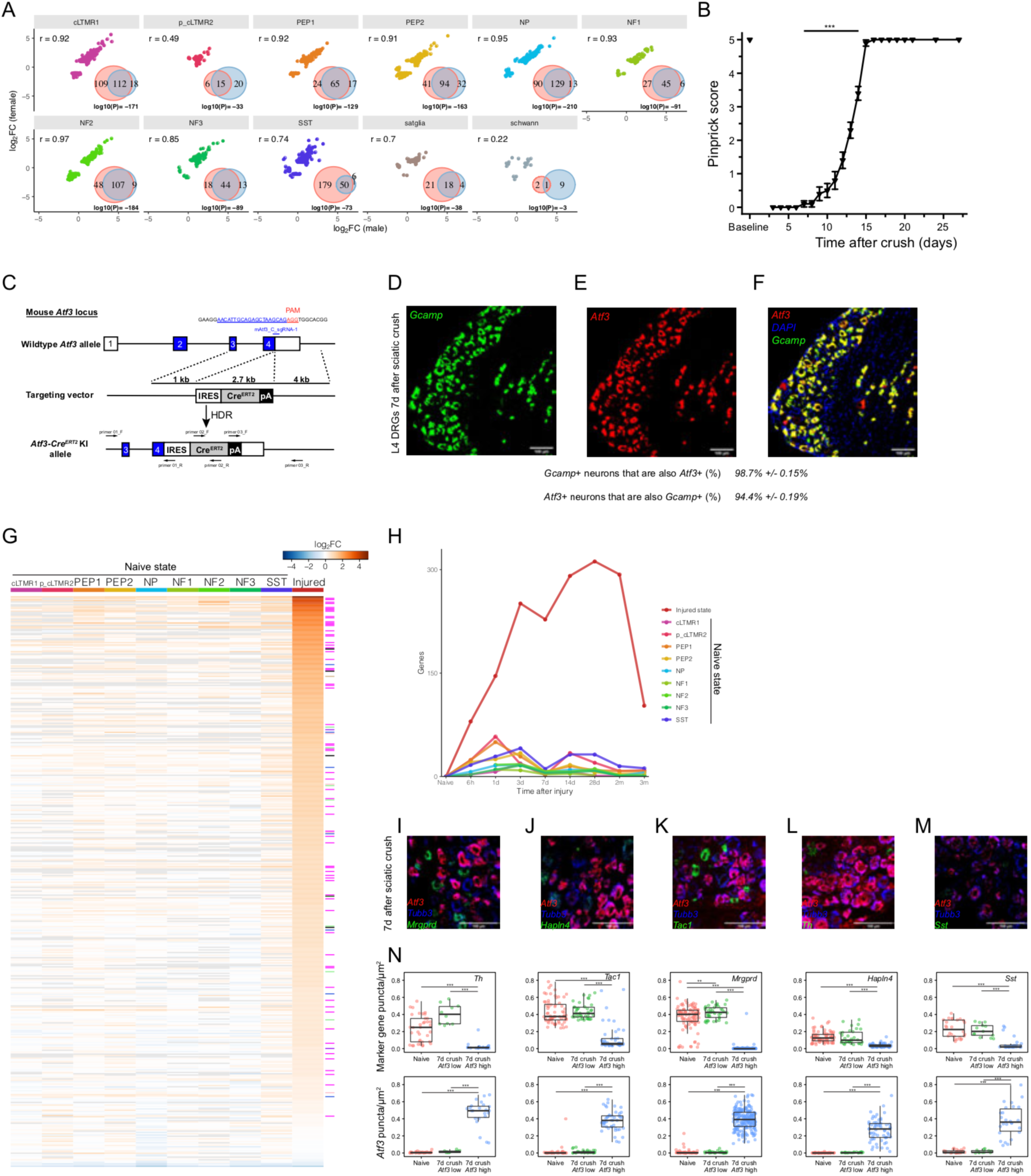
Molecular characterization of DRG neurons after sciatic nerve crush. **(A)** Sex differences in gene expression after sciatic nerve crush. Scatter plot displays the log_2_FC (1 week after sciatic nerve crush vs. naive controls) in male (on the x-axis) and female (on the y-axis) mice of the set of genes that are significantly regulated by sciatic nerve crush in either males or females (FDR < 0.01, log_2_FC > |1|, 1 week after sciatic nerve crush vs. naive) in each cell type. Pearson correlations are displayed. Venn diagrams of the above injury-regulated genes in male and female after sciatic nerve crush. Hypergeometric test *P*-values are displayed. **(B)** Recovery of sensory function after sciatic nerve crush as measured by the pinprick assay in C57/Bl6 mice. Pinprick responses recover to baseline 15 days after sciatic crush (n=11 female mice, 1-way repeated measured ANOVA, *F*(1, 10) = 1180, *P* = 1×10^-11^, Bonferroni post-hoc, \*\*\**P* < 0.001). **(C)** Diagram of the *Atf3* locus in the *Atf3-Cre^ERT2^* transgenic mouse. An IRES_*Cre^ERT2^*_pA cassette was inserted at the 3’UTR of the mouse *Atf3* locus to avoid interfering with endogenous *Atf3* expression. **(D-F)** Fluorescent *in situ* hybridization (FISH) images of an L4 *Atf3-Cre^ERT2^*;G*camp6f* mouse DRG 1 week after sciatic nerve crush stained with probes against *Gcamp6f* (green, D), *Atf3* (red, E), and colocalization of DAPI (blue), *Gcamp6f* (green) and *Atf3* (red) (F). There is a very high degree of colocalization of *Atf3* and the *Gcamp6f* reporter, suggesting this mouse is a reliable injury reporter. **(G)** Heatmap displaying the log_2_FC (sciatic crush compared to naive) of the 438 common injury-induced genes identified in Figure 4D (rows) in each neuronal subtype (columns). Differential expression for the neuronal subtypes in the “naive state” at any time point after sciatic crush was performed by comparing these nuclei to their respective naive neuronal subtype. Differential expression for the nuclei in the “injured state” at any time point sciatic crush was performed by comparing these nuclei to all naive nuclei. Gray color indicates a gene is not expressed in that cell type. Genes that have previously been described as regeneration-associated genes (Chandran et al., 2016) are labeled by the color of their gene module described in that study (e.g. magenta box denotes the gene is a member of the magenta cluster). **(H)** Time course of the number of significantly upregulated genes (FDR < 0.01, log_2_FC > 1) in each neuronal subtype after sciatic nerve crush. Nuclei after sciatic nerve crush that were considered to be in the “naive state” were compared to naive neurons of the corresponding subtype. Neurons classified as injured after sciatic nerve crush were compared to all naive neurons. Colors of each line correspond to the cell type indicated in the legend. **(I-M)** FISH images of L4 mouse DRGs stained with probes against *Atf3* (I-M, red), *Tubb3* (I-M, blue) and *Mrgprd* (I, green), *Hapln4* (J, green), *Tac1* (K, green), *Th* (L, green) or *Sst* (M, green). **(N)** Quantification of FISH puncta from Figures S4I-M. DRG neurons were first identified by *Tubb3* fluorescence, then divided into *Atf3*-high (injured) and *Atf3*-low (naive) populations (see methods). On the box plot, each dot represents an individual cell, boxes indicate quartiles, and whiskers are 1.5-times the interquartile range (Q1-Q3). The median is a black line inside each box. Significance testing by 1-way ANOVAs were all *P <* 0.001: Th (n = 36 [naive], 10 [7d *Atf3* low], 23 [7d *Atf3* high]), *F*(2, 66) = 38.34, *Atf3*(on Th slides), *F*(2, 66) = 209.09; Tac1 (n = 68 [naive], 40 [7d *Atf3* low], 45 [7d *Atf3* high]), *F*(2, 150) = 85.03, *Atf3*(on Tac1 slides), *F*(2, 150) = 420.46; Mrgprd (n = 100 [naive], 41 [7d *Atf3* low], 209 [7d *Atf3* high]), *F*(2, 347) = 899.72, *Atf3*(on Mrgprd slides), *F*(2, 347) = 780.13; Hapln4 (n = 80 [naive], 32 [7d *Atf3* low], 62 [7d *Atf3* high]), *F*(2, 171) = 57.81, *Atf3*(on Hapln4 slides), *F*(2, 171) = 235.85; Sst (n = 26 [naive], 13 [7d *Atf3* low], 26 [7d *Atf3* high]), *F*(2, 62) = 29.31, *Atf3*(on Sst slides), *F*(2, 62) = 74.81; Tukey HSD post-hoc testing (***: p < 0.001, **: p < 0.01, *: p < 0.05). Neurons expressing each marker gene are abundant in *Atf3*-low DRG neurons 1 week after sciatic crush, whereas *Atf3*-high DRG neurons contain significantly fewer marker gene puncta. cLTMR = C-fiber low threshold mechanoreceptor; PEP = peptidergic nociceptor; NP = non-peptidergic nociceptor; NF = *Nefh+* A-fiber low threshold mechanoreceptors; SST = *Sst+* pruriceptors.

**Figure S6, related to Figure 5.**
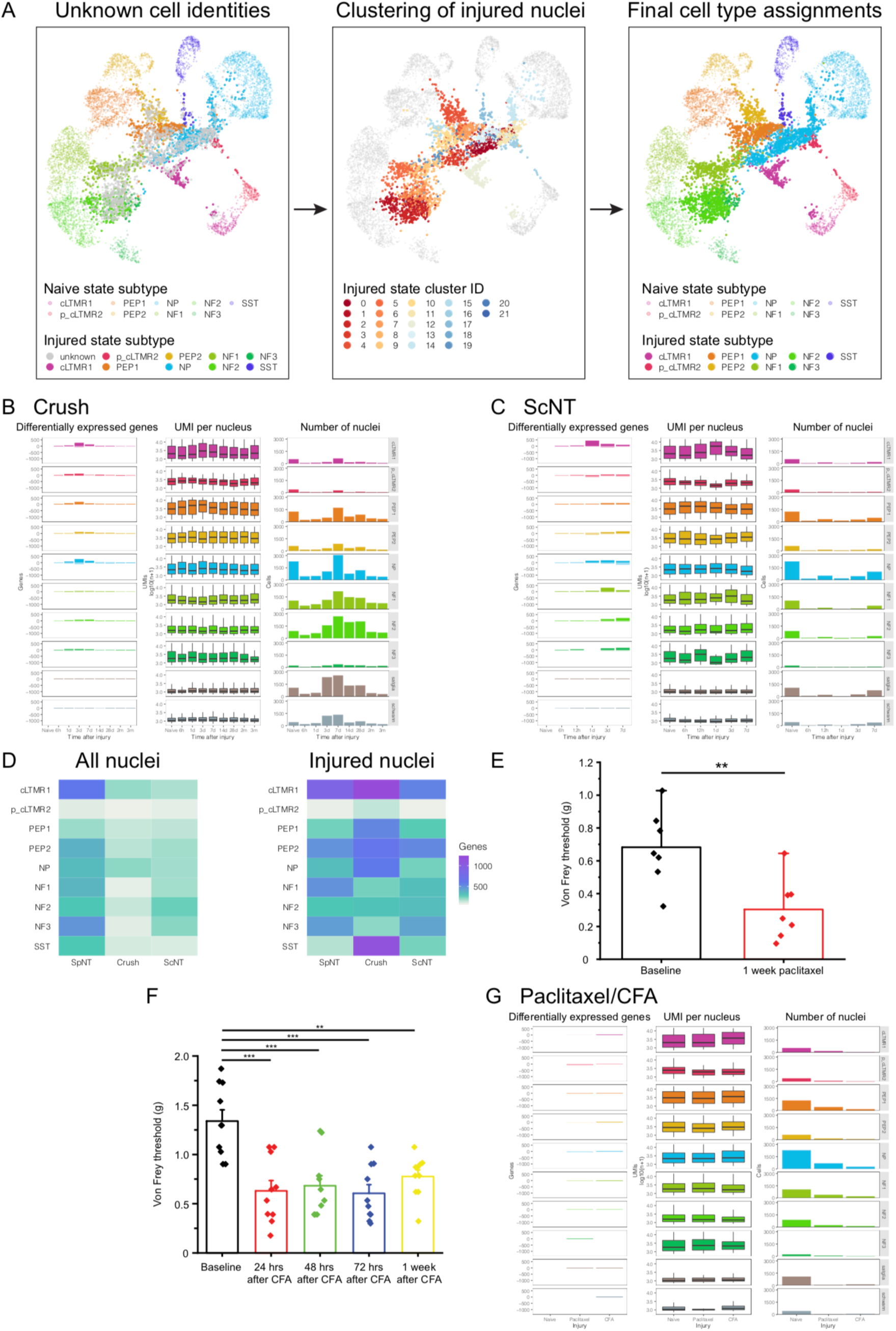
Comparison of transcriptional changes induced by axotomy and other animal models of pain in DRG neurons. **(A)** Co-clustering of known injured neuronal subtypes after spinal nerve transection (SpNT) with sciatic crush and sciatic nerve transection (ScNT) “injured state” nuclei of unknown subtype. UMAP plots displaying 2,500 neurons randomly sampled from naive, and 2,500 neurons randomly sampled from each of the three injury models after they were clustered together. Nuclei of unknown neuronal subtype that co-clustered with clusters of known neuronal subtypes from SpNT (middle, nuclei colored by clusterID), were then assigned the respective neuronal subtype of that cluster (right, see methods). Nuclei are colored by their neuronal subtype (left, right) with “naive state” faded and “injured state” bolded. **(B, C, and G)** Summary of the number of significant differentially expressed genes (left, positive number indicates significantly upregulated genes with FDR < 0.01 and log_2_FC > 1, and negative number denotes significantly down-regulated genes with FDR < 0.01 and log_2_FC < −1), UMI (middle log_10_ transformed), and total number of nuclei for cell type (right) at each time point in sciatic nerve crush (B), ScNT (C), and paclitaxel or Complete Freund’s Adjuvant (CFA) treatments (F). Boxes indicate quartiles and whiskers are 1.5-times the interquartile range (Q1-Q3). Data outside 1.5-times the interquartile range are omitted for clarity. The median is a black line inside each box. **(D)** Heatmap of the number of significant (FDR < 0.01, log_2_FC > 1) injury-induced genes for each cell type and injury model. Differential expression analyses were performed either by comparing all nuclei 3d and 7d after injury vs. nuclei from the respective neuronal subtype in naive animals (left) or by comparing only nuclei in the “injured state” 3d and 7d after injury to the respective neuronal subtype from naive mice (right). The advantage of performing differential expression on all nuclei (left) is that we can identify cell-type-specific gene expression changes at early time points after injury prior to the emergence of the “injured state,” although these analyses are limited by the inclusion of unaxotomized neurons in the analysis. The advantage of performing differential expression specifically on injured nuclei is that it allows us to more directly compare gene expression programs between injury models without including unaxotomized neurons. Because the SpNT model axotomizes most neurons, while crush and ScNT only axotomize ∼50% of neurons, the similar number of gene expression changes between “injured state” neurons across the three models suggest the gene expression program at the level of an individual injured neuron is quite similar between distal and proximal axonal injury. The number of nuclei used for differential expression analysis in each neuronal subtype was equal across injury models and set to the number of nuclei in the injury model with the fewest number of nuclei sequenced. **(E)** Von Frey behavioral measurement of mechanical sensitivity in C57/Bl6 mice at baseline or 1 week after every-other-day treatment with 4mg/kg paclitaxel. Paclitaxel treatment causes a significant mechanical allodynia 1 week after start of treatment (n=7 mice, paired two-tailed Student’s t-test, \*\**P* = 0.006). **(F)** Von Frey behavioral measurement of mechanical sensitivity in C57/Bl6 mice after hindpaw injection of 20µL CFA. CFA treatment causes significant mechanical allodynia 24 hours after treatment that persists for at least 7 days after treatment (n=10 mice, 1-way repeated measured ANOVA, *F*(4, 36) = 12.3, p=0.005, Bonferroni post-hoc \*\**P* < 0.01, *** *P* < 0.001). cLTMR = C-fiber low threshold mechanoreceptor; PEP = peptidergic nociceptor; NP = non-peptidergic nociceptor; NF = *Nefh+* A-fiber low threshold mechanoreceptors; SST = *Sst+* pruriceptors.

**Figure S7, related to Figure 7.**
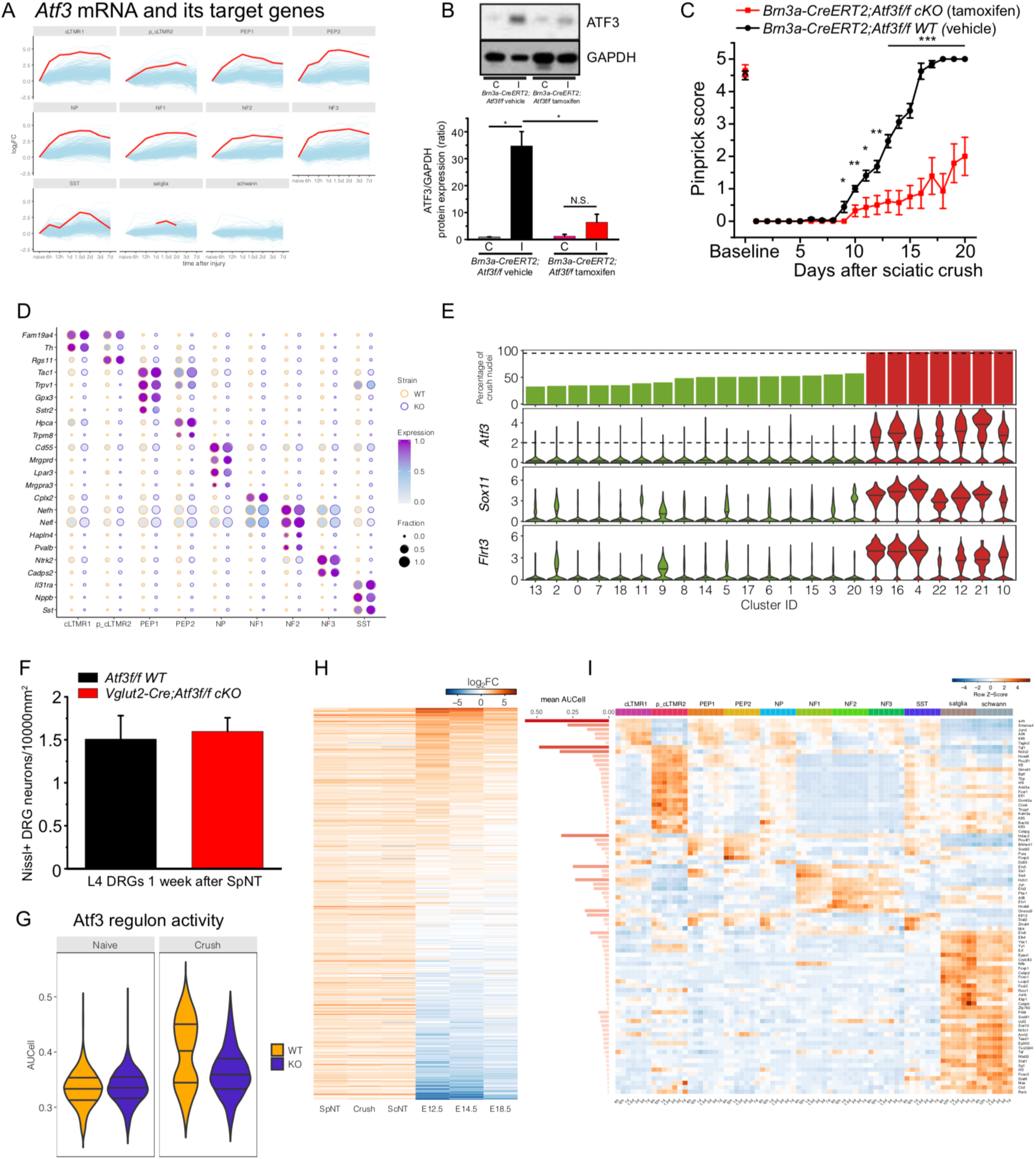
Transcription factor analysis of the injury-induced gene expression program. **(A)** Log_2_FC (spinal nerve transection [SpNT] compared to naive) of *Atf3* mRNA (red line) and ATF3 target genes (light blue lines) at each time point and DRG cell type. Each line represents regulation of one gene over time. A break in the line occurs if the gene is below the expression threshold at a specific time point. **(B)** Representative Western Blot (top) and quantification (bottom) of ATF3 protein in DRG protein extract from ipsilateral and contralateral L3-L5 DRG neurons from *Brn3a-Cre^ERT2^;Atf3f/f* mice 1 week after sciatic nerve crush. ATF3 is significantly induced in ipsilateral injured but not in uninjured contralateral DRG neurons in *Brn3a-Cre^ERT2^;Atf3f/f* mice treated with vehicle (retaining Atf3) (p=0.04, n=2, two-tailed Student’s t-test). In *Brn3a-Cre^ERT2^;Atf3f/f* mice treated with tamoxifen (which causes loss of Atf3), Atf3 is not significantly induced in ipsilateral L3-L5 DRG neurons 1 week after sciatic nerve crush (p=0.23, n=2, two-tailed Student’s t-test). For quantification (bottom), the ratio of ATF3/GAPDH protein levels was calculated from the Western Blot data. Data are mean±SEM. **(C)** Recovery of sensory function as measured by the pinprick assay in vehicle and tamoxifen treated *Brn3a-Cre^ERT2^;Atf3f/f* mice after sciatic nerve crush. Sciatic nerve crush causes a loss of sensory responses in the ipsilateral hindpaw, followed by a recovery over time associated with sensory neuron regeneration. The pinprick responses of vehicle treated *Brn3a-Cre^ERT2^;Atf3f/f* mice (n=8, black line) recover to baseline within 16 days after sciatic nerve crush (1-way repeated measures within subjects ANOVA, lower bound *F*(1,7) = 343, *P* = 3.3×10^-7^). The pinprick responses of tamoxifen treated *Brn3a-Cre^ERT2^;Atf3f/f* DRG mice (n=7, red line) show a significant delay in the time course of sensory function recovery (2-way repeated measures between subjects ANOVA, *F*(1, 13) = 40.2, *P* = 2.6×10^-5^, Bonferroni post-hoc, *** *P* < 0.001), suggesting a slower rate of sensory neuron regeneration. **(D)** Dot plot of neuronal subtype-specific marker genes (rows) in neuronal subtypes (columns) from naive *Atf3f/f* (WT, orange circles) or *Vglut2-Cre;Atf3f/f* (cKO, purple circles) DRGs. The fraction of nuclei expressing a marker gene is calculated as the number of nuclei in each cell type that express a gene (> 0 counts) divided by the total number of naive nuclei in the respective cell type. Expression in each cell type is calculated as the mean scaled counts of the marker gene relative to the highest mean-scaled counts of that gene across cell types. **(E)** Bar plot showing the percent of nuclei 7 days after sciatic crush [100 * crush nuclei / (naive + crush nuclei)] within each neuronal cluster (top row) and violin plots showing log_2_-normalized expression of selected injury-induced genes in each cluster (second to fourth rows). Note that sciatic crush only injures approximately 50% of lumbar DRG neurons sequenced. Cluster ID (x-axis) corresponds to cluster number assignment from Seurat (see methods). Clusters are classified as “injured state” (red) if they are comprised of > 95% nuclei from sciatic crush mice and have a median normalized *Atf3* expression > 0.8 SD from mean (corresponding to > log_2_-normalized expression of 2). All other clusters are classified as “naive state” (green). **(F)** Quantification of Nissl+ DRG neurons in L4 DRG sections from *Vglut2-Cre;Atf3f/f* cKO (n=4 sections, red) and *Atf3f/f* WT (n=4 sections, black) mice 1 week after SpNT. There is no significant difference in DRG neuron density (*P* = 0.71, two-tailed Student’s t-test), suggesting there is no DRG neuron death at this time point. Data are mean ± SEM. **(G)** Violin plot of ATF3 regulon enrichment (AUCell score, see methods). All neuronal nuclei are grouped by genotype (WT or cKO) and injury (naive or crush). Lines in the violins indicate the lower quartile, median, and upper quartile. One-way ANOVA: *F*(3, 11737) = 1391.28, *P* < 0.001; Tukey HSD post-hoc testing *P* > 0.05 for naive cKO vs. naive WT, p < 0.001 for all other pair-wise comparisons. **(H)** Regulation of the 438 common injury-induced genes (rows, from Figure 4C) after SpNT, crush, ScNT, and embryonic development. Heatmap shows the log_2_FC from differential expression analysis of “injured state” nuclei in each injury model compared to all naive nuclei as well as the log_2_FC between RET+ DRG neurons at 3 embryonic time points (E12.5, E14.5, E18.5) compared to adult RET+ DRG neurons (see methods). **(I)** Heatmap displays the transcription factors (rows) identified by SCENIC analysis (see methods) as having their consensus binding sites enriched within expressed genes in naive and SpNT cell types at all time points (columns). Colors on the heatmap represent row-normalized average AUCell scores for nuclei in each cell type and time point. AUCell scores are a SCENIC metric of the activity of a transcription factor in each cell; higher AUCell scores indicate greater predicted activity of a transcription factor on its target genes in a given cell. The horizontal bar plots for each transcription factor indicates the mean AUCell score (not row-normalized) across all cell types and time points. cLTMR = C-fiber low threshold mechanoreceptor; PEP = peptidergic nociceptor; NP = non-peptidergic nociceptor; NF = *Nefh+* A-fiber low threshold mechanoreceptors; SST = *Sst+* pruriceptors.

