## Supplementary figures and images for "Transcriptional reprogramming of distinct peripheral sensory neuron subtypes after axonal injury"

### High-resolution Figures 1-7

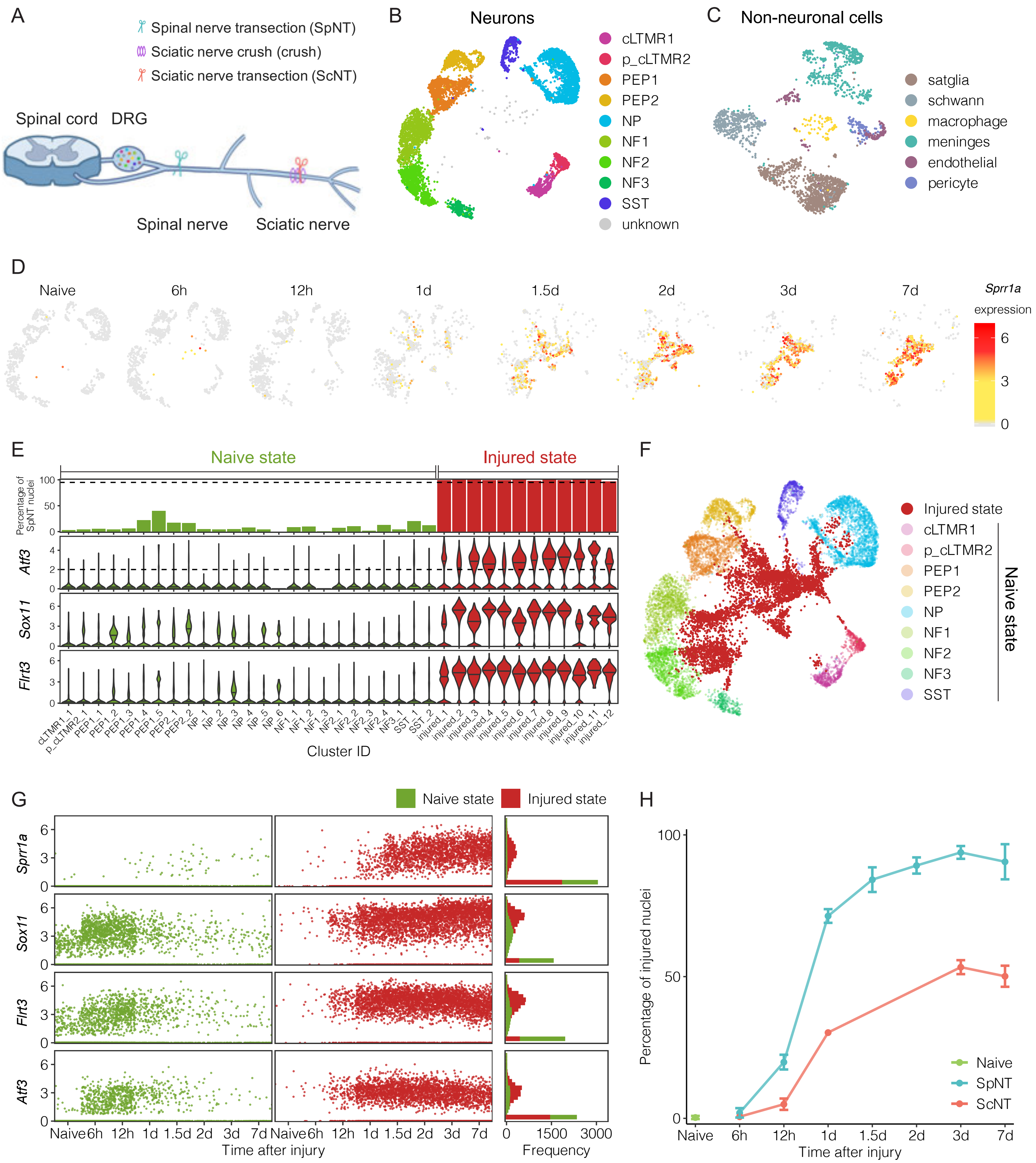

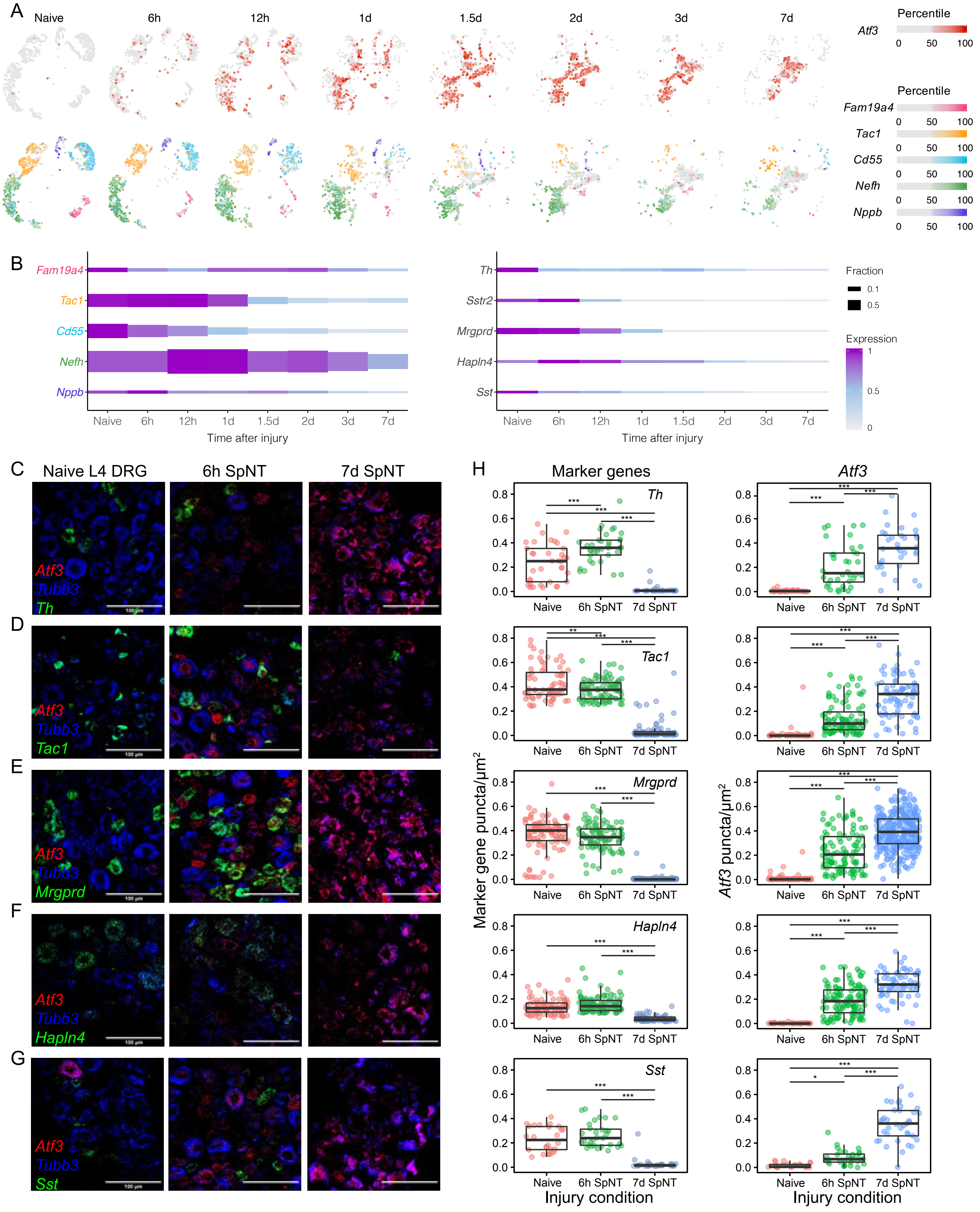

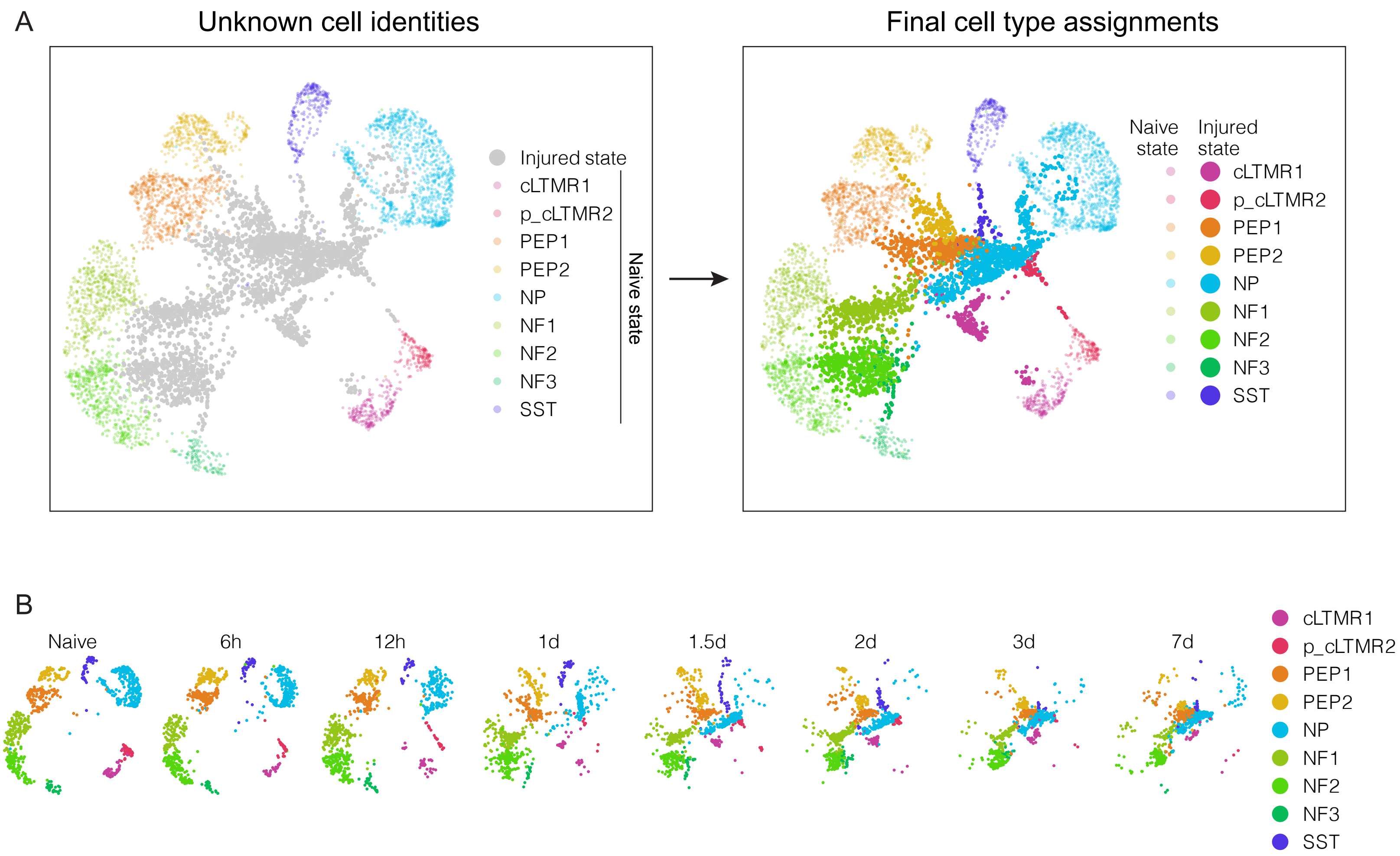

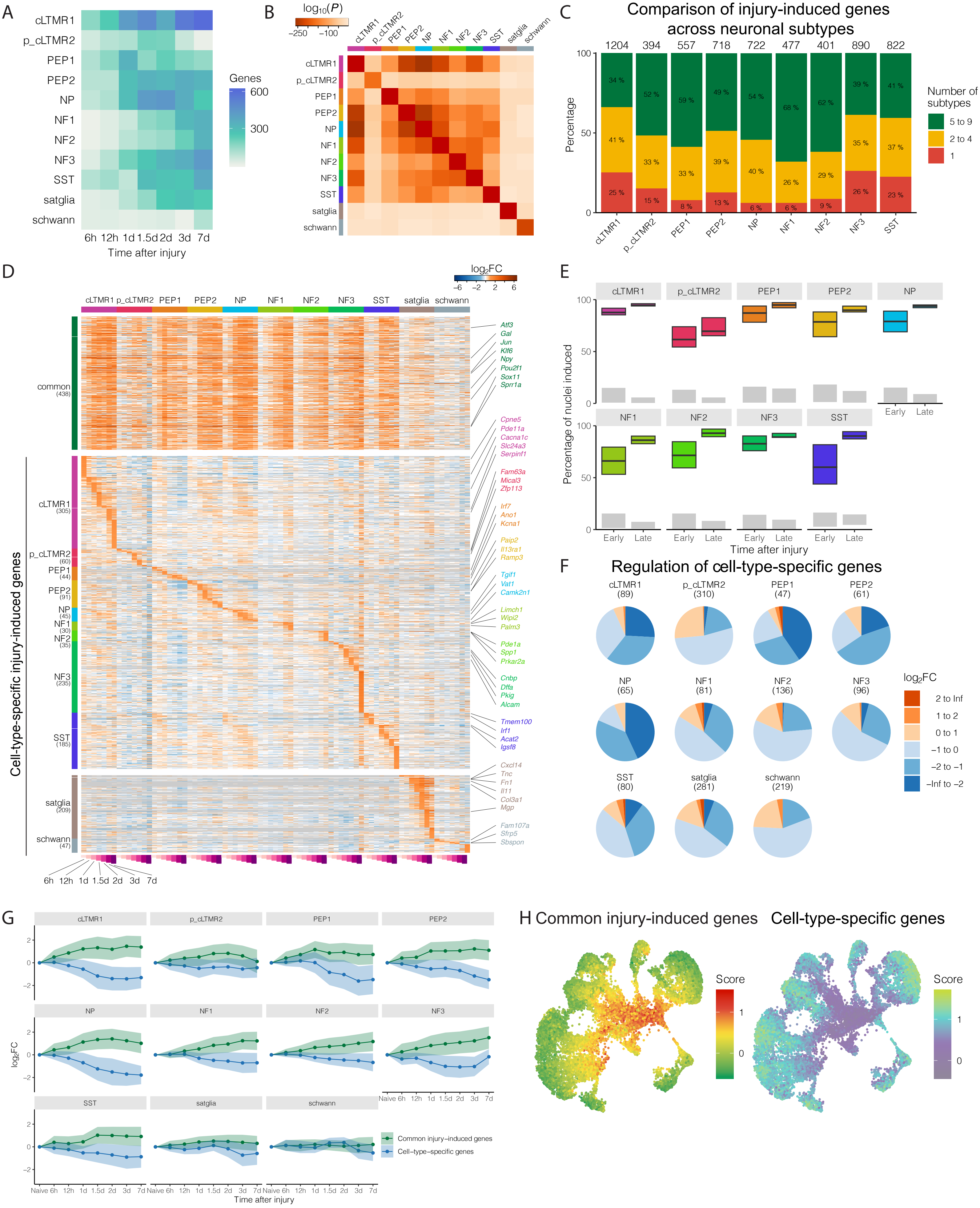

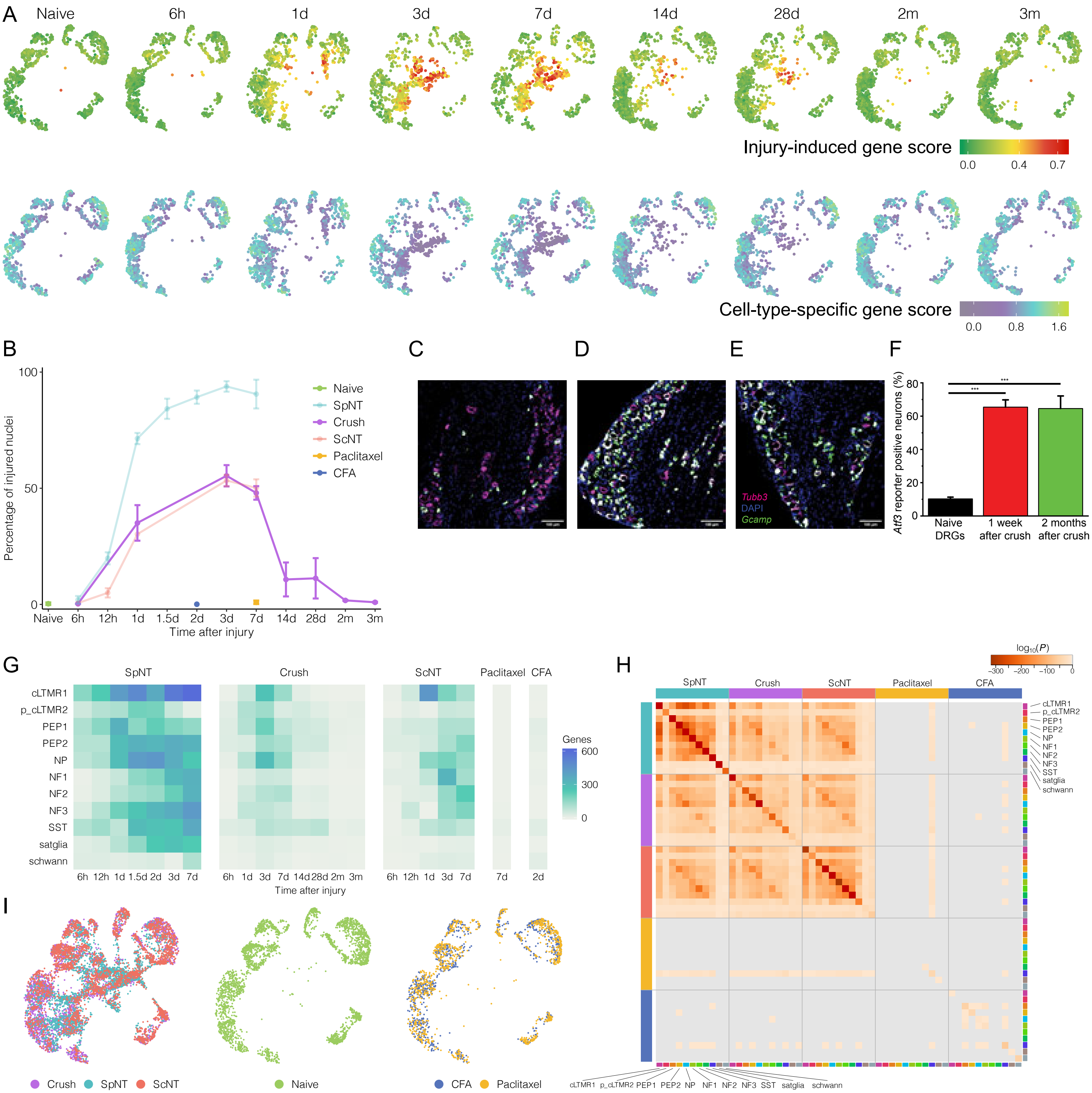

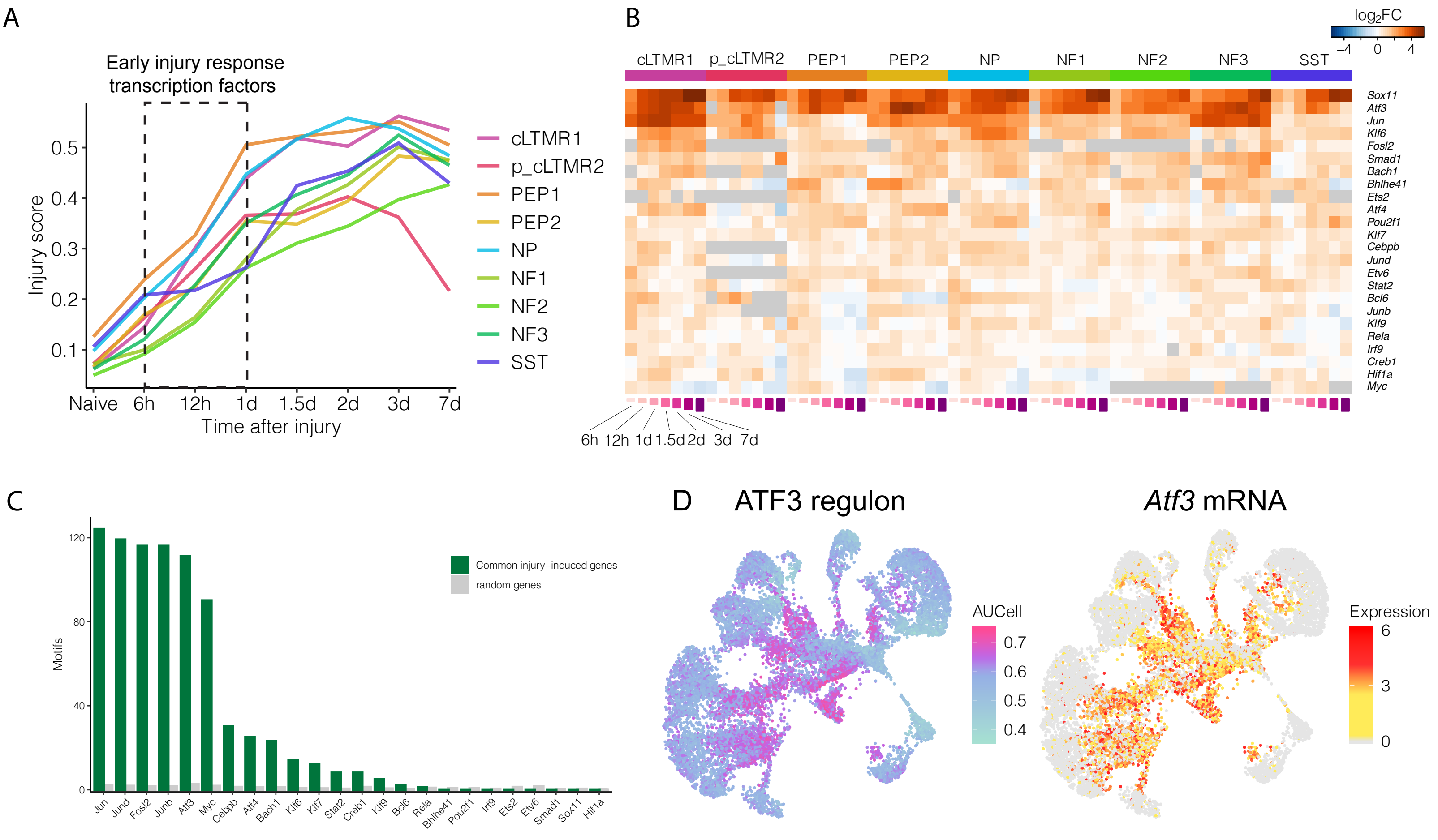

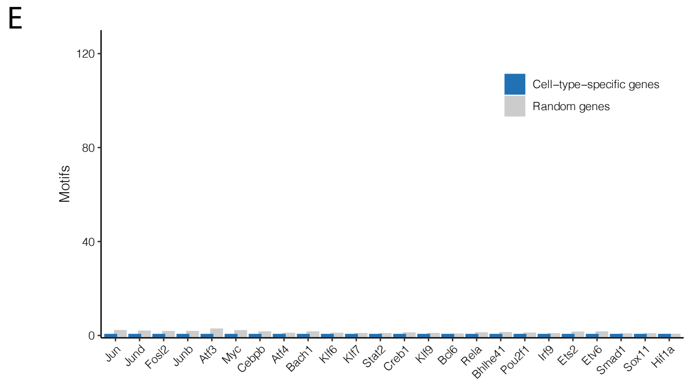

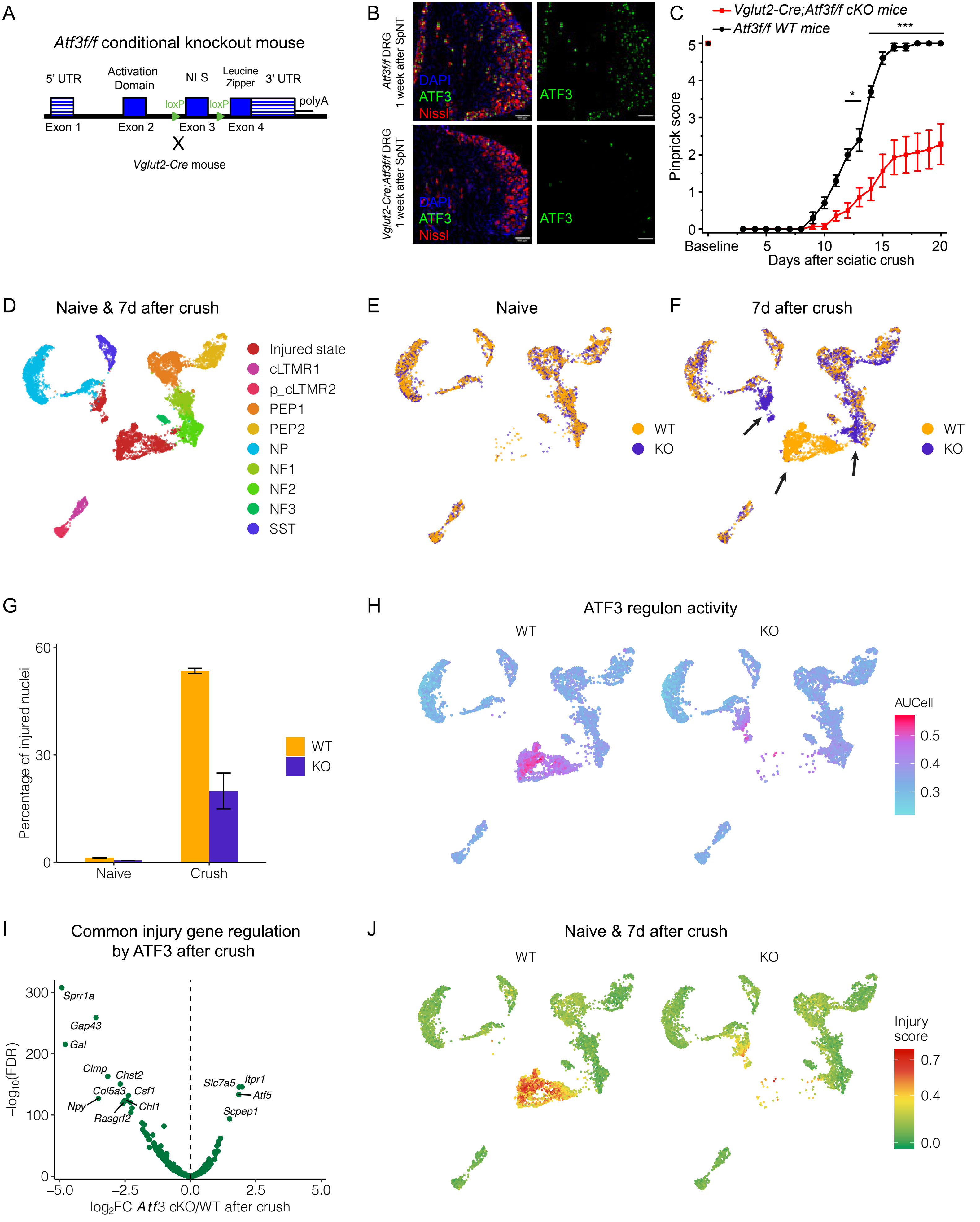
