## Supplementary material for "Transcriptional reprogramming of distinct peripheral sensory neuron subtypes after axonal injury": High-resolution Figures S1-S7

Supplementary Figure 1 -- related to Figure 1

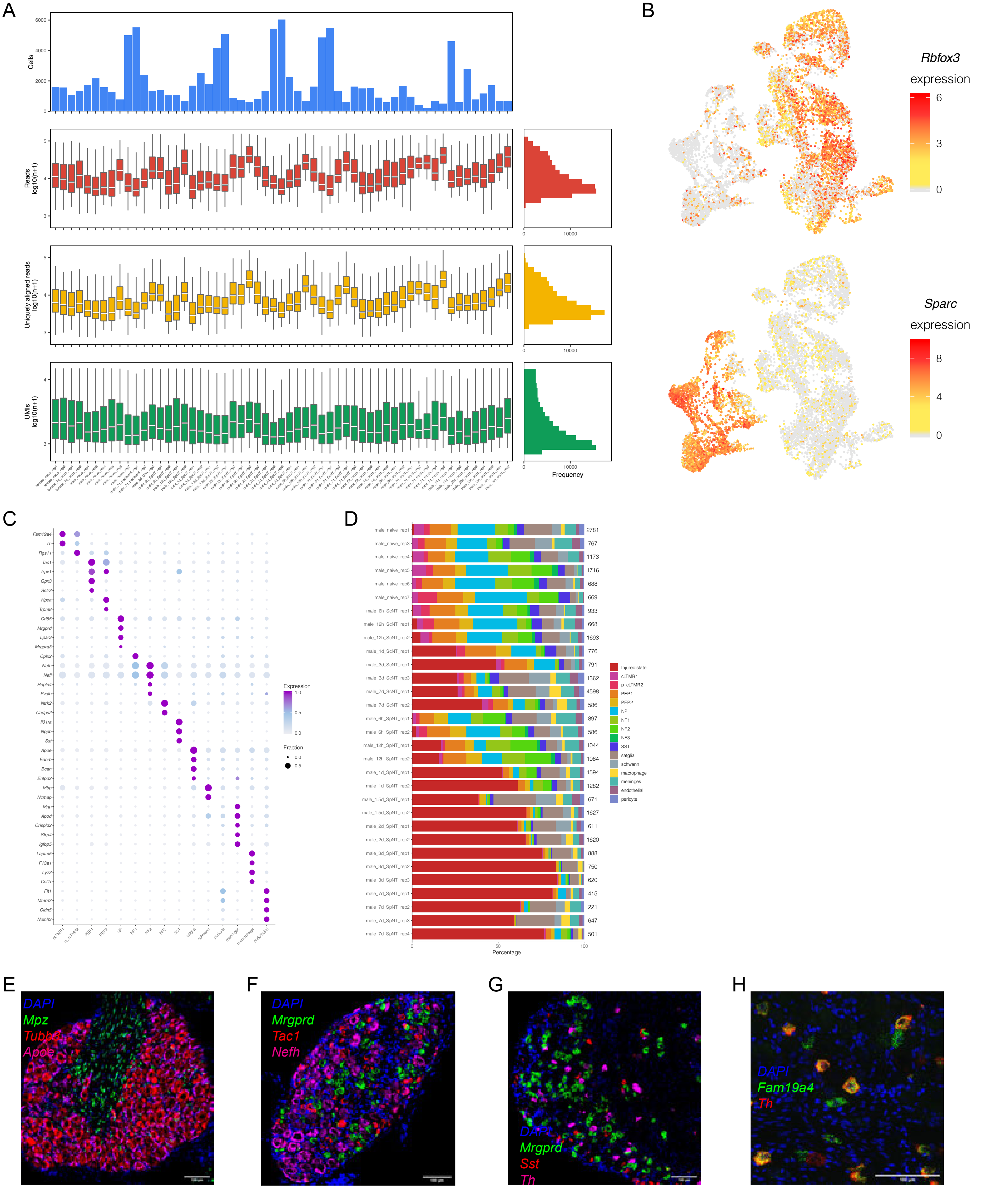

Supplementary Figure 2 -- related to Figure 1

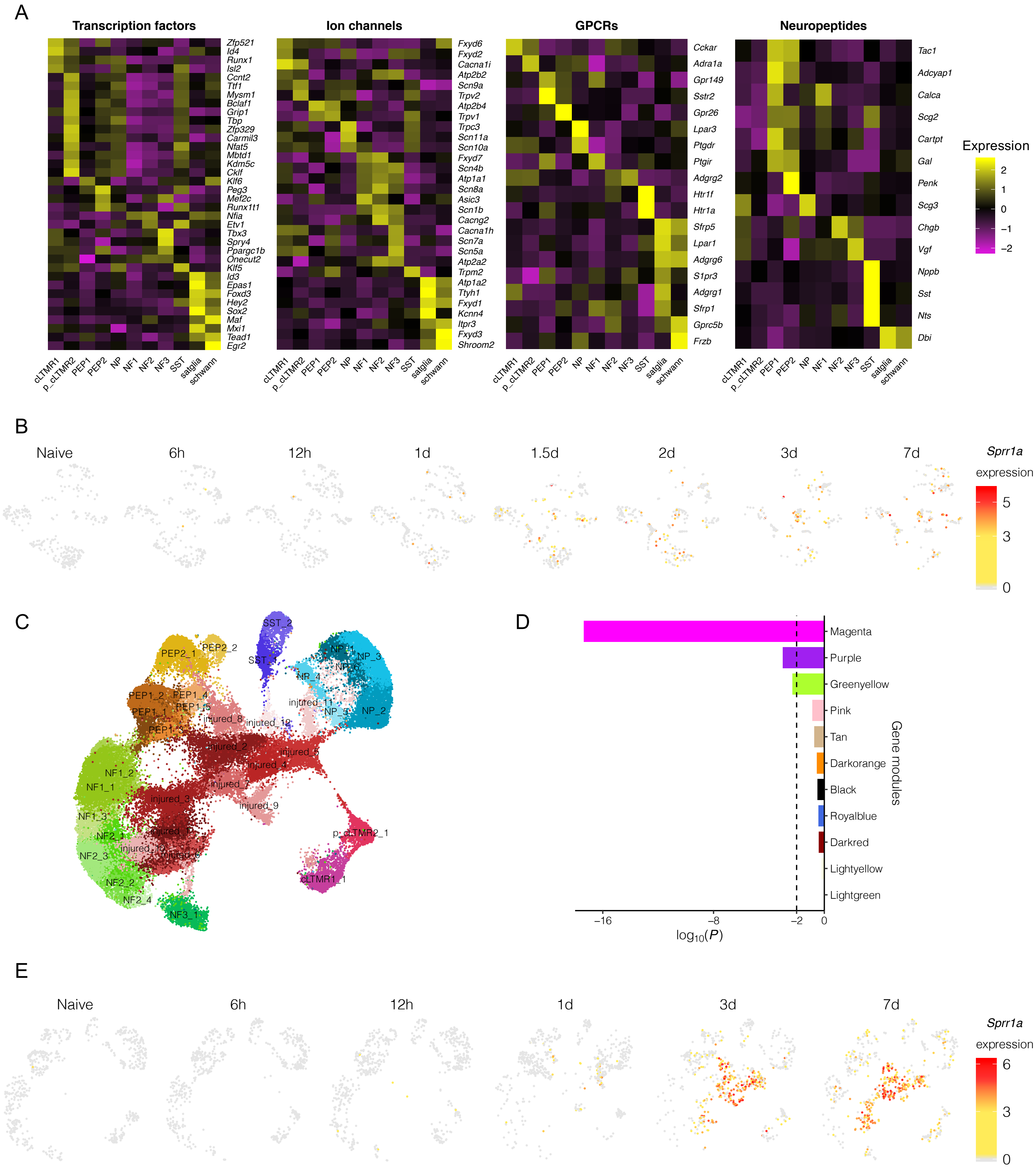

Supplementary Figure 3 -- related to Figure 3

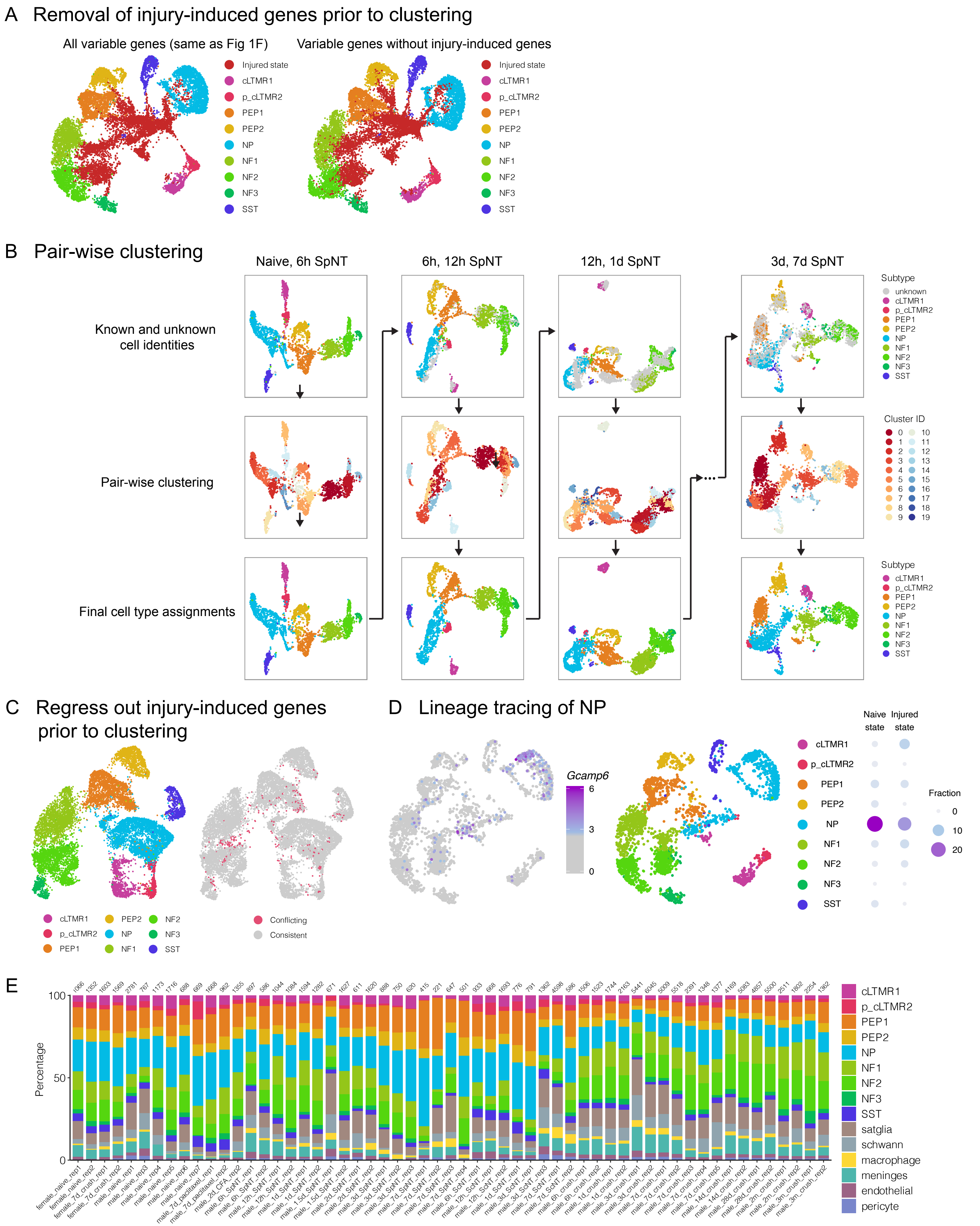

Supplementary Figure 4 -- related to Figure 4

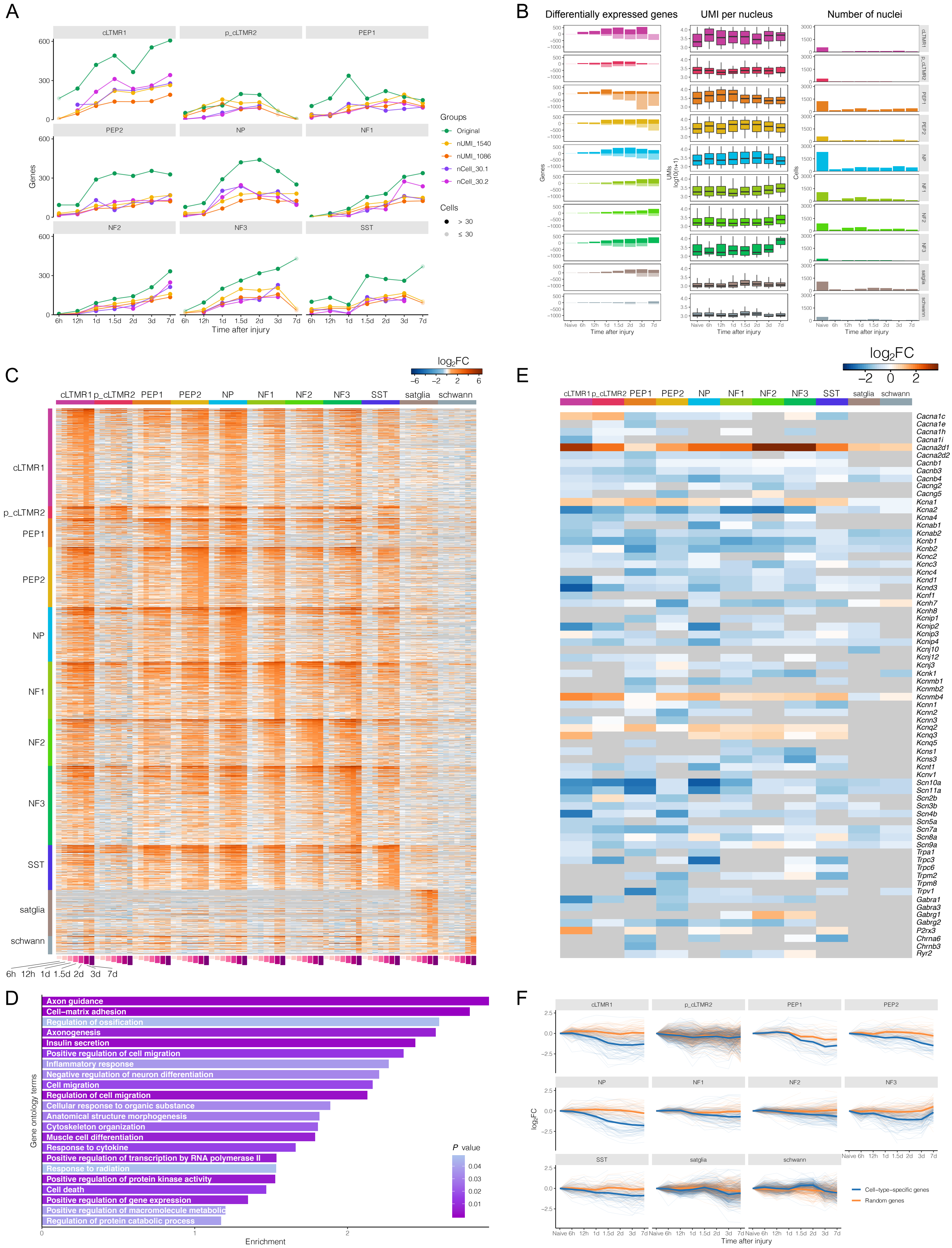

### Supplementary Figure 5 -- related to Figure 5

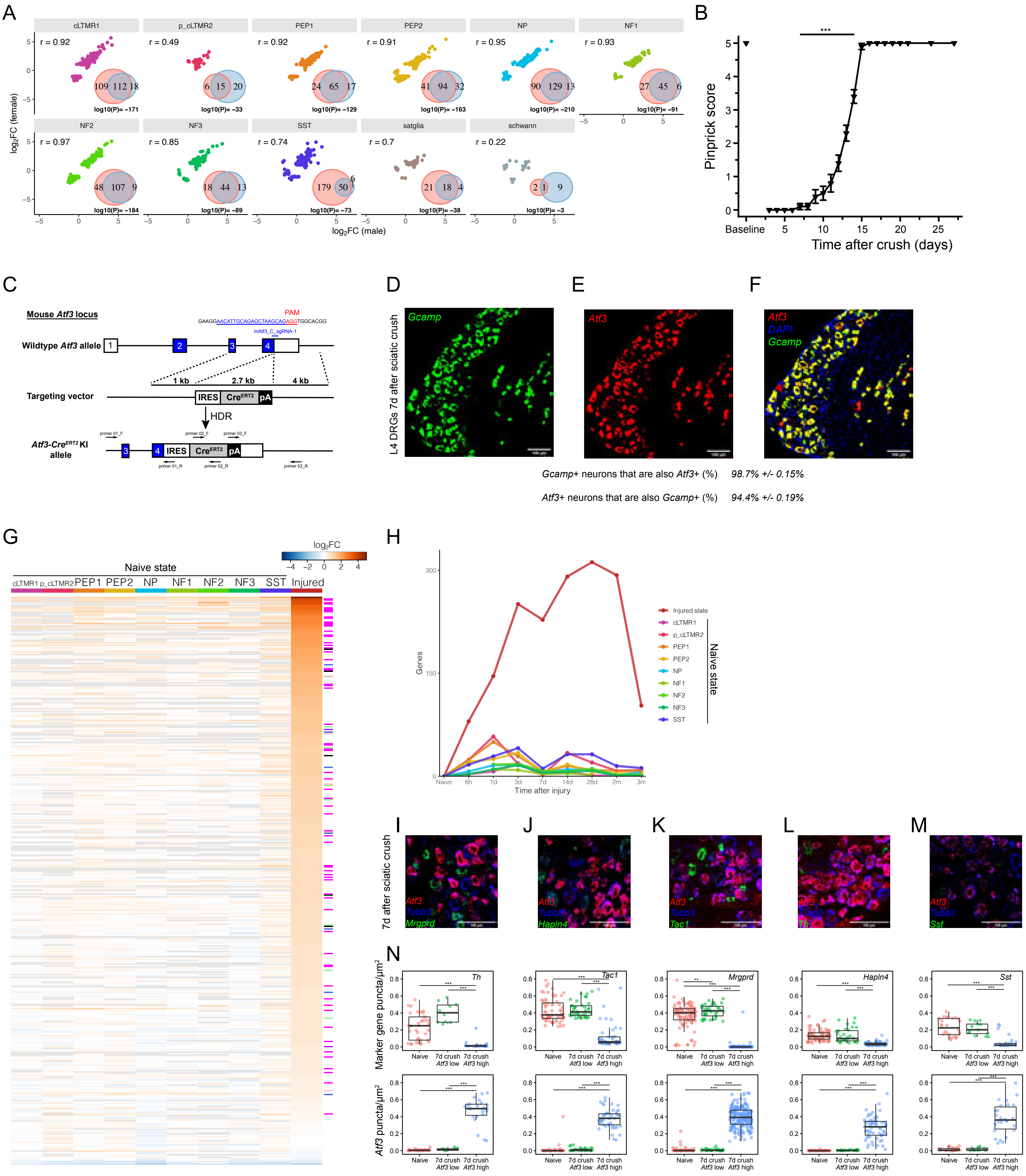

#### Supplementary Figure 6 -- related to Figure 5

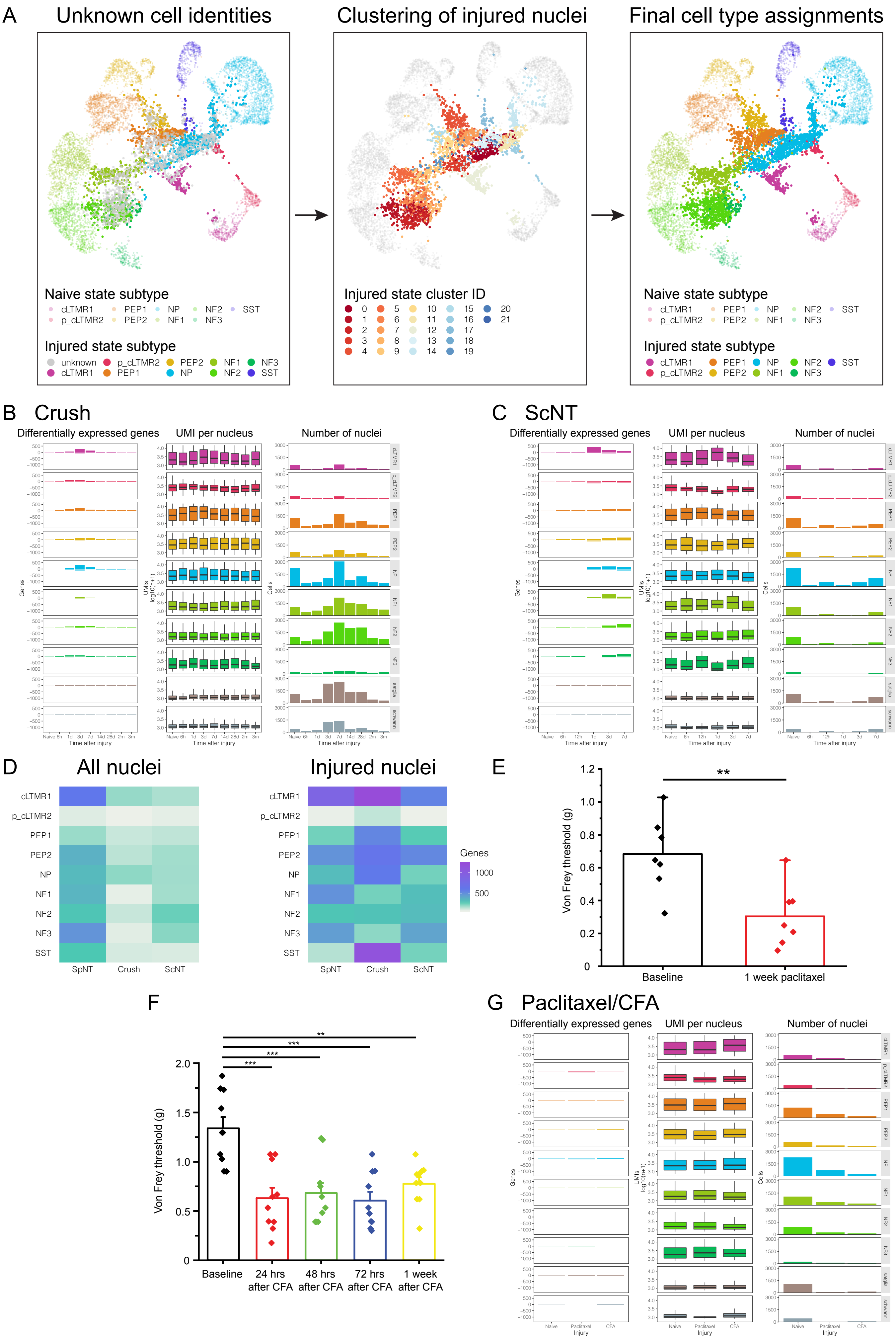

### Supplementary Figure 7 -- related to Figures 6 and 7

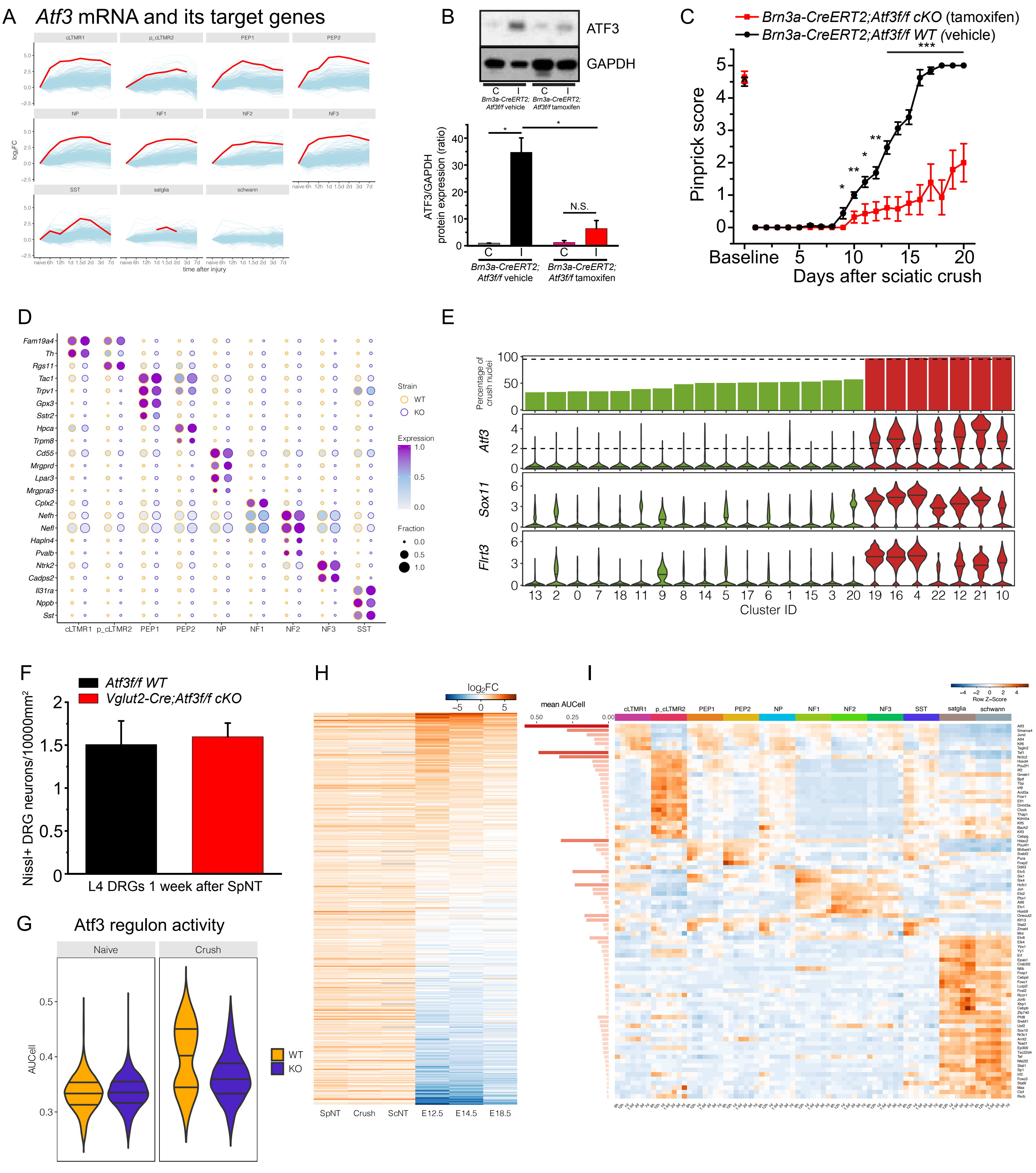
